# OTUs clustering should be avoided for defining oral microbiome

**DOI:** 10.1101/2021.08.09.455616

**Authors:** A Regueira-Iglesias, L Vázquez-González, C Balsa-Castro, T Blanco-Pintos, VM Arce, MJ Carreira, I Tomás

**Affiliations:** Oral Sciences Research Group, Special Needs Unit, Department of Surgery and Medical-Surgical Specialties, School of Medicine and Dentistry, Universidade de Santiago de Compostela, Health Research Institute Foundation of Santiago (FIDIS); Santiago de Compostela, Spain; Centro Singular de Investigación en Tecnoloxías Intelixentes and Departamento de Electrónica e Computación, Universidade de Santiago de Compostela; Health Research Institute Foundation of Santiago (FIDIS); Santiago de Compostela, Spain; Department of Physiology and Center for Disease in Molecular Medicine and Chronic Diseases, Universidade de Santiago de Compostela, Spain

**Keywords:** computational biology, DNA primers, genes, high-throughput nucleotide sequencing mouth, microbiota, rRNA

## Abstract

This *in silico* investigation aimed to: 1) evaluate a set of primer pairs with high coverage, including those most commonly used in the literature, to find the different oral species with 16S rRNA gene amplicon similarity/identity (ASI) values ≥97%; and 2) identify oral species that may be erroneously clustered in the same operational taxonomic unit (OTU) and ascertain whether they belong to distinct genera or other higher taxonomic ranks.

Thirty-nine primer pairs were employed to obtain amplicon sequence variants (ASVs) from the complete genomes of 186 bacterial and 135 archaeal species. For each primer, ASVs without mismatches were aligned using BLASTN and their similarity values were obtained. Finally, we selected ASVs from different species with an ASI value ≥97% that were covered 100% by the query sequences. For each primer, the percentage of species-level coverage with no ASI≥97% (SC-NASI≥97%) was calculated.

Based on the SC-NASI≥97% values, the best primer pairs were OP_F053-KP_R020 for bacteria (65.05%), KP_F018-KP_R002 for archaea (51.11%), and OP_F114-KP_R031 for bacteria and archaea together (52.02%). Eighty percent of the oral-bacteria and oral-archaea species shared an ASI≥97% with at least one other taxa, including *Campylobacter*, *Rothia*, *Streptococcus*, and *Tannerella*, which played conflicting roles in the oral microbiota. Moreover, around a quarter and a third of these two-by-two similarity relationships were between species from different bacteria and archaea genera, respectively. Furthermore, even taxa from distinct families, orders, and classes could be grouped in the same cluster.

Consequently, irrespective of the primer pair used, OTUs constructed with a 97% similarity provide an inaccurate description of oral-bacterial and oral-archaeal species, greatly affecting microbial diversity parameters. As a result, clustering by OTUs impacts the credibility of the associations between some oral species and certain health and disease conditions. This limits significantly the comparability of the microbial diversity findings reported in oral microbiome literature.

## INTRODUCTION

Studies in a variety of publications over the last decade have assessed the mouth’s microbiome using high throughput 16S ribosomal RNA (rRNA) gene sequencing (Zaura et al. 2021). To facilitate the analysis of complex microbial communities like the oral environment, amplicons derived from this technology are typically clustered into operational taxonomic units (OTUs) that are intended to correspond to taxonomic clades (Edgar 2013). Specifically, sequences are clustered based on a given similarity threshold, usually set at 97%, which has been conventionally regarded as the species-level correspondent (Stackebrandt and Goebel, 1994; Zaura et al. 2021).

Numerous OTU clustering algorithms have been integrated into the popular sequence-analysis pipelines, such as QIIME2 (Bolyen et al. 2019), mothur (Schloss et al. 2009), and USEARCH (Edgar 2010). Overall, existing methods for grouping 16S rRNA gene amplicons into OTUs can be categorized in three ways: *de novo*, closed-reference, and open-reference (Wei et al. 2021). However, none of these approaches produce the same results in terms of obtaining OTUs, even when using the same dataset (He et al. 2015; Westcott and Schloss 2015). Moreover, even the same method can yield distinct results after only a minor parameter change (Wei et al. 2021).

In addition, it has been reported that different species can have very highly similar 16S rRNA gene sequences (Schloss 2021; Vĕtrovský and Baldrian 2013), which may lead to the grouping of distinct taxa in the same OTU. In fact, around 25% of OTUs constructed using the widely adopted ≥97% identity threshold have been found to contain gene sequences from multiple species (Schloss 2021; Vĕtrovský and Baldrian 2013). These estimates were slightly different depending on the gene region studied but reached up to 35% for regions 4-5 (Schloss 2021). Consequently, the construction of an OTU table can be affected, as can, by extension, taxonomic assignments and microbial diversity results. To the best of our knowledge, there is no research concerning how many different oral taxa, and which specific oral species, could be erroneously clustered in the same OTU. Consequently, the objectives of the present *in silico* investigation are to: 1) evaluate a set of primer pairs with high coverage, including those most commonly used in the literature, to find the different oral prokaryotic species with 16S rRNA gene amplicon similarity/identity (ASI) values ≥97%; and 2) identify oral species that may be erroneously clustered in the same OTU and ascertain whether they belong to distinct genera or other higher taxonomic ranks.

## MATERIALS AND METHODS

The complete analysis protocol applied in the present study is detailed in Figure 1.

**Figure 1.**
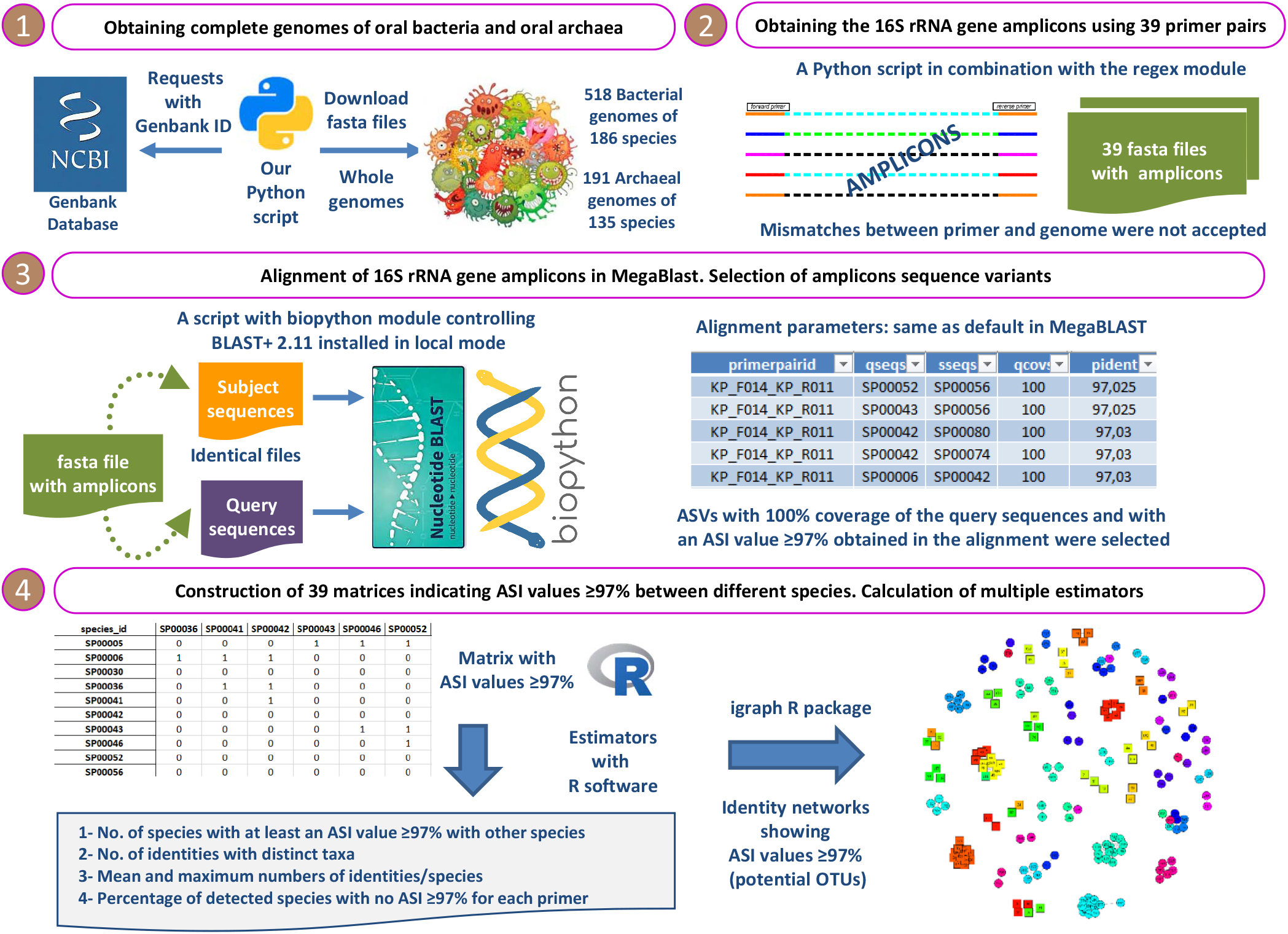
The complete analysis protocol applied in the present *in-silico* study.

### Obtaining complete genomes of oral bacteria and oral archaea

The information on the bacterial taxa present in the oral cavity was obtained from the eHOMD website (Escapa et al. 2018). Of the 2074 genomes available on the site, we only selected the 528 that had a complete sequencing status for use in the research. These complete genomes have one or more GenBank identifiers (Clark et al. 2016), which were employed to access the complete sequences stored in the NCBI database (NCBI Resource Coordinators 2016). Additionally, an initial list of 177 different oral archaea and their corresponding GenBank identifiers (Clark et al. 2016), obtained as part of previous research conducted by our group (Regueira-Iglesias et al. 2021.a), enabled us to access their complete sequences in the NCBI database. Integrating the “Entrez Programming Utilities (E-utilities)” tool (National Center for Biotechnology Information 2010) in our Python script (Python Software Foundation 2020) allowed us to acquire the URLs needed to retrieve the information of interest from various NCBI databases, including Taxonomy (Schoch et al. 2020), RefSeq (O’Leary et al. 2016), and GenBank (Clark et al. 2016). The oral-bacteria and oral-archaea genomes were then downloaded and the taxonomy of each of them was determined.

### Selecting the primer pairs and obtaining the 16S rRNA gene amplicons

Thirty-three primer pairs with the best *in silico* coverage, as identified in earlier research by our group, were selected, along with the six primer pairs used the most in the oral-microbiome literature (Regueira-Iglesias et al. 2021.a) (Appendices Tables 1 and 2).

Applying our script in combination with Python’s regex module (Barnett 2020), the direct and reverse sequences of each primer pair were used to obtain, in *silico*, the amplicons of the 16S rRNA genes identified in the selected genomes. Amplicons were included for evaluation when the sequences of both primers in a pair (forward and reverse) had no mismatches against the genome. All the amplicon sequence variants (ASVs) detected were included for the downstream analysis. Coincident ASVs within a given species were removed. Finally, the amplicons were assigned a taxonomic classification to the species level and a corresponding variant number. The taxonomic classifications and species identifiers of the 186 oral-bacterial and 135 oral-archaeal species from which we obtained the amplicons are set out in Appendices Tables 3 and 4.

### Alignment of 16S rRNA gene amplicons in MegaBlast

A script with the NcbiblastnCommandline wrapper from the BLAST (Altschul et al. 1990) module of Biopython (Cock et al. 2009) was developed to manage BLAST+ 2.11 (Camacho et al. 2009) in the local mode from Python. This enabled the data obtained in the alignments to be easily transferred for later analysis on Python. The alignment parameters were configured to be the same as the default settings in MegaBLAST (Altschul et al. 1990) since these settings were appropriate for the alignment between sequences with a similarity ≥95%.

All the retrieved ASVs without mismatches were stored in a single fasta file for each of the analyzed primer pairs. Every file was inserted both as a subject and a query in BLASTN (Chen et al. 2015) and all the sequences were then aligned. The results obtained were used to select the ASVs with 100% coverage of the query sequences with an ASI value ≥97%, i.e., the alignments with the following estimates in BLAST+ (Camacho et al. 2009): qcovs= 100%, qcovhsp= 100%, qcovus= 100%, and pident ≥97%. The following alignments were discarded: 1) between amplicons with the same unique identifier; 2) between amplicons with the same species identifier; and 3) duplicates. If two different species had more than one ASI value ≥97% in several ASVs, one of them was chosen at random. The results from the highly similar pairs of species were then stored, including the data on the taxonomic hierarchy of both taxa, using the *pandas* (McKinney 2010) and *xlsxwriter* (McNamara 2013) modules of Python.

### Construction of a matrix with ASI values ≥97% between oral species and calculation of estimators

A similarity matrix was created for each primer pair, where rows and columns had the species identifiers, and cells indicated ASI values ≥97%. We then developed a script in R (R Core Team 2016) through which we calculated the following estimates for each analyzed primer pair: 1) the number of species with at least one ASI≥97% with other taxa; 2) the total number of identities or potential OTUs with distinct taxa; 3) the mean and maximum numbers of identities per taxa. In addition, we estimated the percentage of detected species with no ASI≥97% for each primer (species coverage no ASI≥97% = SC-NASI≥97%). This parameter was then used as a criterion for selecting the primers associated with a smaller number of oral species that may be erroneously grouped in the same OTUs. Finally, we identified bacterial and archaeal species with a 97% identity threshold, paying special attention to those of different genera or higher taxonomic ranks.

## RESULTS

### Evaluation of the primer pairs against the oral-bacteria and oral-archaea genomes

If the primers used most in the literature were excluded, those with short amplicon lengths, (unlike the SC percentages) had the lowest SC-NASI≥97% values for both bacteria (S= 39.54%) and archaea (S= 40.44%) compared to the medium length and long primers (M= 45.81% and 45.37%, respectively; L= 48.38% and 44.32%, respectively).

Concerning the bacteria-specific primer pairs, the number of species with an ASI≥97% and the number of total identities ranged from 37 and 32 with the most widely used primer, KP_F031-KP_R021 (S; SC-NASI≥97%= 54.30%), to 120 and 277 with OP_F066-KP_R040 (M; SC-NASI≥97%= 24.19%), respectively (Table 1). This latter primer also had the lowest SC-NASI≥97% value, while OP_F053-KP_R020 detected the highest number of species with NASI≥97% (M; SC-NASI ≥97%= 65.05%). In addition, except for OP_F053-KP_R020, all the bacteria-specific primers had a maximum number of identities/species above five. Eleven primers produced the maximum identities/species ≥10 values, i.e., they created potential clusters that contained the highest number of very similar taxa. OP_F066_KP_R040 contained the most different species (=24) with maximum identities/species ≥10 (Figure 2; Appendix Table 5).

**Figure 2.**
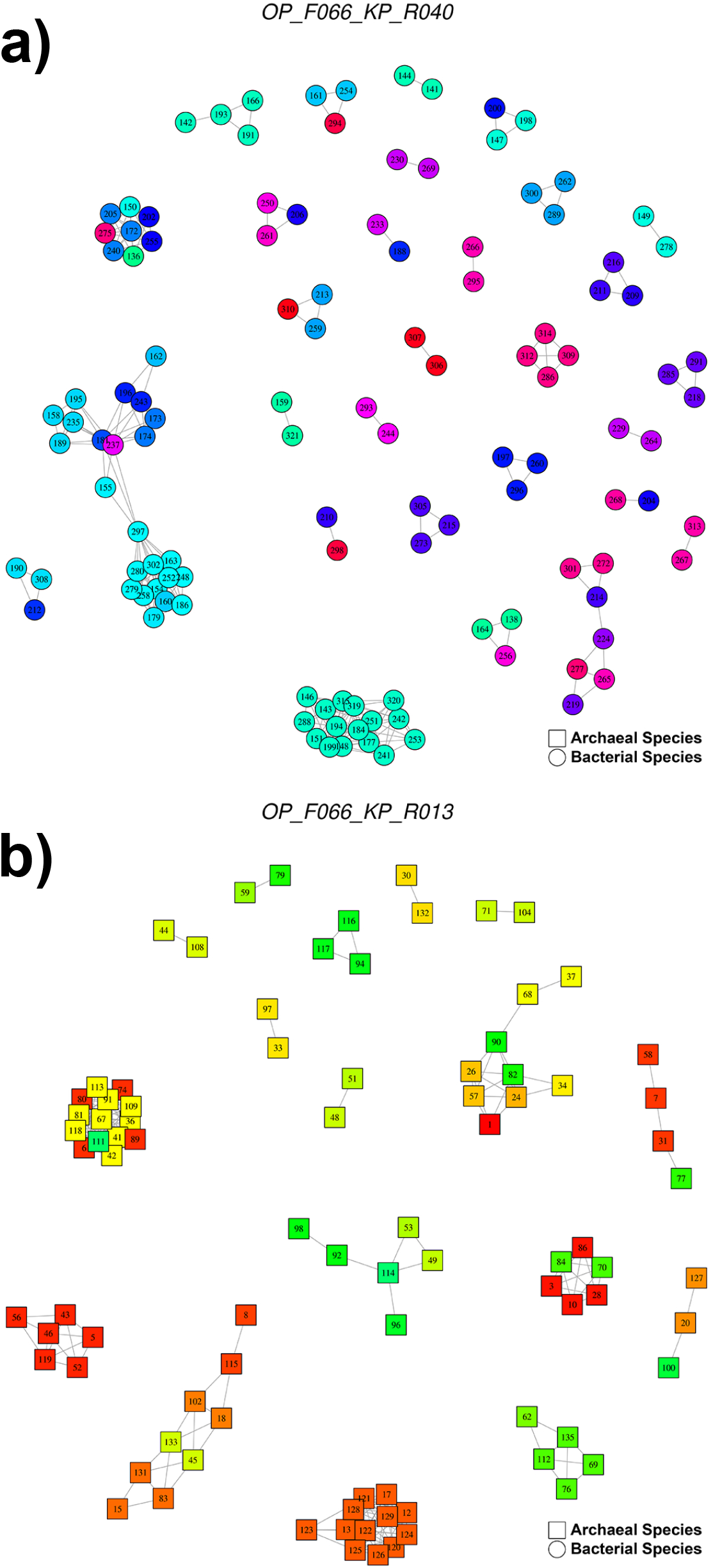
Networks showing the potential OTUs with a 97% identity threshold obtained with the primer pairs: a) OP_F066-KP_R040 for bacteria (120 species ASI >97%, 277 identities); b) OP_F066-KP_R013 for archaea (89 species ASI >97%, 240 identities). In the graphs, each node represents an oral species, the color indicates the genus and the number refers to the species identifier, whose assigned species are detailed in the Appendices Tables 3 and 4. Each edge represents the presence of a 97% similarity between different species. resulting in clusters of possible OTUs. The graphs were made using the igraph package (version 1.2.6) (Csardi and Nepusz, 2005)

**Table 1.**
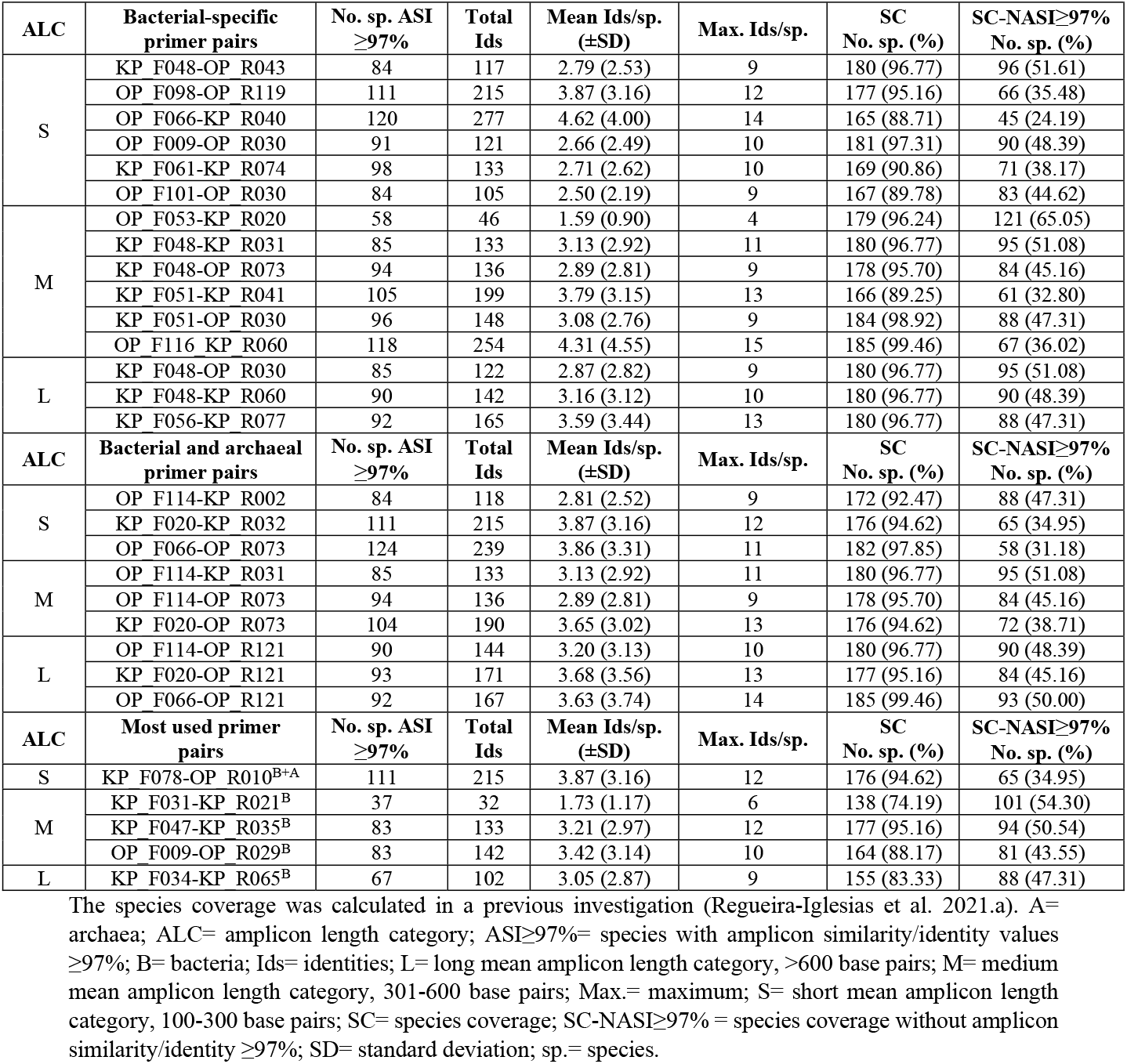
Number of species with amplicon similarity values ≥97%, the number of identities, and the SC and SC-NASI≥97% values obtained by each primer pair analyzed against the oral bacteria genomes.

Concerning the archaea-specific primer pairs, the number of species with an ASI≥97% and the number of identities ranged from 24 and 96 with the widely used KP_F014-KP_R011 (L; SC-NASI≥97%= 12.59%) to 89 and 240 with OP_F066-KP_R013 (S; SC-NASI≥97%= 29.63%), respectively (Table 2). The former primer detected the lowest number of species without an ASI≥97%, and KP_F018-KP_R002 the highest (S; SC-NASI≥97%= 51.11%). Moreover, all the archaea-specific primers had a maximum number of identities/species ≥10, meaning that all of them produced potential clusters that contained the highest number of very similar taxa. OP_F066_KP_R013 and OP_F066_OP_R016 (L) had the highest number of distinct species (=25) with maximum identities/species ≥10 (Figure 2; Appendix Table 6).

**Table 2.** Number of species with amplicon similarity values ≥97%, the number of identities, and the SC and SC-NASI≥97% values obtained by each primer pair analyzed against the oral archaea genomes.

| ALC | Archaeal-specific primer pairs | No. sp. ASI $\geq 97\%$ | Total Ids | Mean Ids/sp. (±SD) | Max. Ids/sp. | SC No. sp. (%) | SC-NASI$\geq 97\%$ No. sp. (%) |
| --- | --- | --- | --- | --- | --- | --- | --- |
| S | KP F018-KP R002 | 60 | 125 | 4.17 (2.98) | 11 | 129 (95.56) | 69 (51.11) |
|  | OP F066-KP R013 | 89 | 240 | 5.39 (4.35) | 13 | 129 (95.56) | 40 (29.63) |
| M | KP F018-KP R032 | 62 | 136 | 4.39 (3.66) | 13 | 130 (96.30) | 68 (50.37) |
|  | KP F018-OP R073 | 59 | 150 | 5.09 (4.35) | 13 | 122 (90.37) | 63 (46.67) |
|  | KP F020-KP R013 | 72 | 189 | 5.25 (4.51) | 12 | 128 (94.81) | 56 (41.48) |
|  | KP F022-KP R063 | 66 | 115 | 3.49 (3.15) | 10 | 124 (91.85) | 58 (42.96) |
| L | OP F114-KP R013 | 66 | 176 | 5.33 (4.72) | 13 | 129 (95.56) | 63 (46.67) |
|  | KP F018-KP R063 | 60 | 134 | 4.47 (3.69) | 11 | 127 (94.07) | 67 (49.63) |
|  | OP F066-OP R016 | 72 | 189 | 5.25 (4.83) | 13 | 124 (91.85) | 52 (38.52) |
| ALC | Bacterial and archaeal primer pairs | No. sp. ASI $\geq 97\%$ | Total Ids | Mean Ids/sp. (±SD) | Max. Ids/sp. | SC No. sp. (%) | SC-NASI$\geq 97\%$ No. sp. (%) |
| S | OP F114-KP R002 | 60 | 126 | 4.20 (2.95) | 11 | 134 (99.26) | 74 (54.81) |
|  | KP F020-KP R032 | 75 | 177 | 4.72 (3.92) | 12 | 134 (99.26) | 59 (43.70) |
|  | OP F066-OP R073 | 95 | 286 | 6.02 (4.14) | 14 | 126 (93.33) | 31 (22.96) |
| M | OP F114-KP R031 | 62 | 136 | 4.39 (3.66) | 13 | 134 (99.26) | 72 (53.33) |
|  | OP F114-OP R073 | 59 | 150 | 5.09 (4.35) | 13 | 126 (93.33) | 67 (49.63) |
|  | KP F020-OP R073 | 71 | 171 | 4.82 (4.03) | 12 | 125 (92.59) | 54 (40.00) |
| L | OP F114-OP R121 | 64 | 168 | 5.25 (4.55) | 13 | 129 (95.56) | 65 (48.15) |
|  | KP F020-OP R121 | 70 | 170 | 4.86 (4.31) | 12 | 129 (95.56) | 59 (43.70) |
|  | OP F066-OP R121 | 77 | 199 | 5.17 (4.68) | 13 | 130 (96.30) | 53 (39.26) |
| ALC | Most used primer pairs | No. sp. ASI $\geq 97\%$ | Total Ids | Mean Ids/sp. (±SD) | Max. Ids/sp. | SC No. sp. (%) | SC-NASI$\geq 97\%$ No. sp. (%) |
| S | KP F078-OP R010<sup>B+A</sup> | 44 | 99 | 4.50 (3.84) | 11 | 86 (63.70) | 42 (31.11) |
| L | KP F014-KP R011<sup>A</sup> | 24 | 96 | 8.00 (4.67) | 13 | 41 (30.37) | 17 (12.59) |

The species coverage was calculated in a previous investigation (Regueira-Iglesias et al. 2021.a). A= archaea; ALC= amplicon length category; ASI $\geq 97\%$ = species with amplicon similarity/identity values $\geq 97\%$ ; B= bacteria; Ids= identities; L= long mean amplicon length category, >600 base pairs; M= medium mean amplicon length category, 301-600 base pairs; Max.= maximum; S= short mean amplicon length category, 100-300 base pairs; SC= species coverage; SC-NASI $\geq 97\%$ = species coverage without amplicon similarity/identity $\geq 97\%$ ; SD= standard deviation; sp.= species.

Finally, for the domains using both the bacterial and archaeal primer pairs, the number of species with an ASI≥97% and the number of identities ranged from 144 and 244 with OP_F114-KP_R002 (S; SC-NASI≥97%= 50.47%) to 219 and 525 with OP_F066-OP_R073 (S; SC-NASI≥97%= 27.73%), respectively (Appendix Table 7). The latter primer also detected the lowest number of species without an ASI ≥97% and OP_F114-KP_R031 the highest (M; SC-NASI≥97%= 52.02%). All the bacterial and archaeal primer combinations had maximum numbers of identities/species ≥10, with OP_F066-OP_R073 having the most distinct taxa (=43; 16 bacteria, 27 archaea) with maximum identities/species ≥10 (Figure 3) (Appendices Tables 5 and 6).

**Figure 3.**
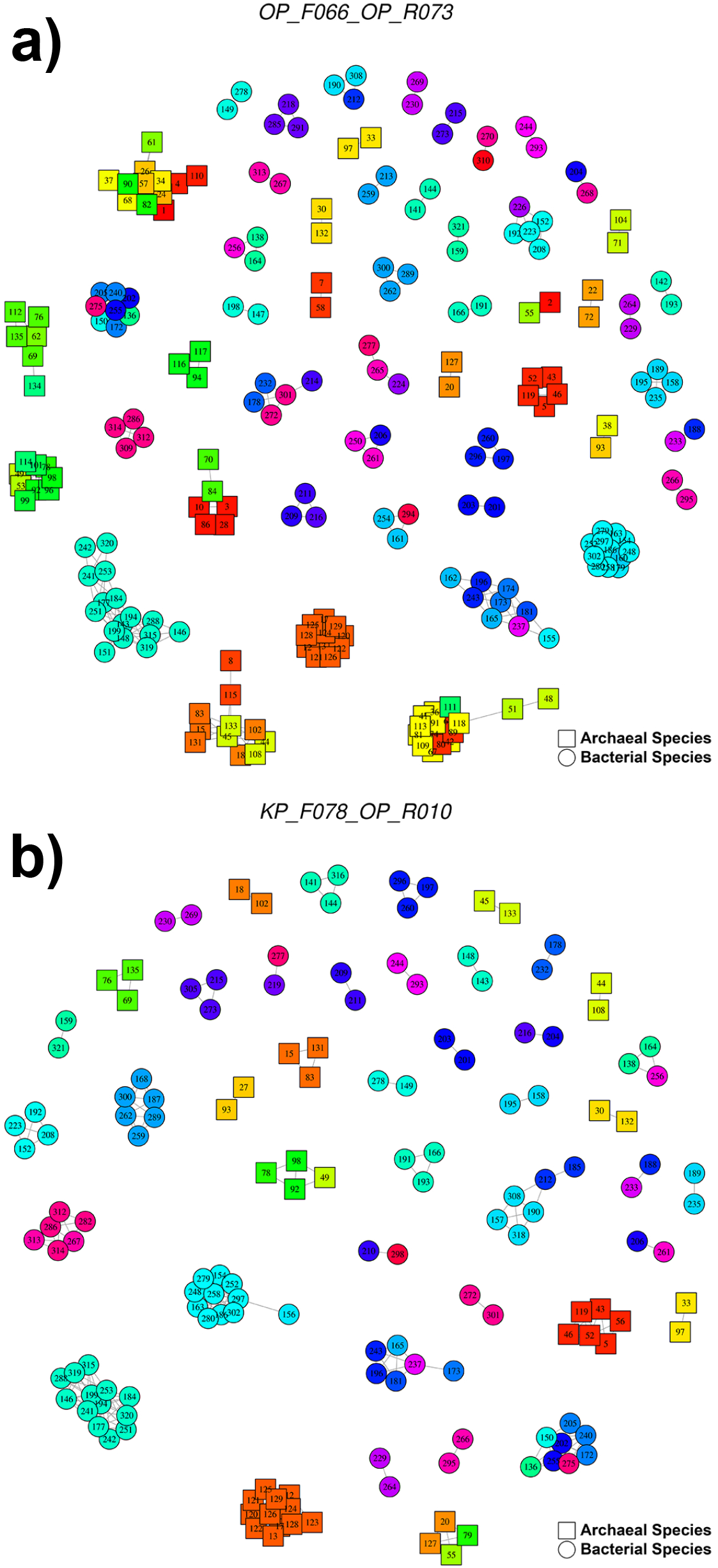
Networks showing the potential OTUs with a 97% identity threshold obtained with the primer pairs for bacteria and archaea: a) OP_F066-OP_R073 (219 species ASI >97%, 525 identities); b) KP_F078-OP_R010 (primer pair widely used in the oral microbiome literature; 155 species ASI >97%, 314 identities). In the graphs, each node represents an oral species, the color indicates the genus and the number refers to the species identifier, whose assigned species are detailed in the Appendices Tables 3 and 4. Each edge represents the presence of a 97% similarity between different species. resulting in clusters of possible OTUs. The graphs were made using the igraph package (version 1.2.6) (Csardi and Nepusz, 2005).

Appendix Table 8 and Appendix Table 9 describe the taxonomy of each of the 30 distinct bacterial and 27 distinct archaeal species, respectively, i.e., the species that could be clustered with a maximum of 10 different taxa when all the analyzed primer pairs were used.

### Taxonomy of the different oral-bacteria and oral-archaea species with an ASI≥97%

One-hundred and forty-nine (80.11%) of the oral-bacteria species analyzed had an ASI ≥97% with at least one distinct species (Appendix Table 10). Conversely, 37 (19.89%) species, including *Filifactor alocis*, *Porphyromonas gingivalis*, and *Prevotella intermedia*, did not have ASI≥97% with other taxa (Appendix Table 11).

All the primers targeting bacteria enabled us to detect 4450 two-on-two relationships between 408 distinct pairs of oral-bacteria species with an ASI≥97% (Appendix Table 12). All primer pairs analyzed identified 18 of these different taxa pairs (frequency= 29, number times that a pair of species had an ASI≥97% in the different primer pairs evaluated), which belonged to the genera *Actinomyces*, *Lactobacillus*, *Neisseria*, *Staphylococcus*, and *Streptococcus.* Conversely, 50 pairs of species were identified once by only one primer pair (frequency= 1). Although the relationships mostly involved species from the same genera (3641; 81.82%), 809 (18.18%) were constituted by taxa from different genera, e.g., the combination of species from *Kleibsella* with others from *Cronobacter*, which occurred more often (frequency= 99) (Appendix Table 13). Furthermore, the pairs of species with an ASI≥97%, such as Enterobacteriaceae and Yersiniaceae (frequency= 153), likewise belonged to distinct families (293; 6.58%); this was also the case for orders (26; 0.58%) like Bacillales and Lactobacillales or Enterobacterales and Pasteurellales (frequencies= 10 and 10, respectively) (Appendices Tables 14-16).

One-hundred and eight (80.00%) of the oral-archaea species evaluated had an ASI≥97% with at least one distinct species (Appendix Table 17). In contrast, 27 (20.00%) taxa, including *Sulfolobus acidocaldarius*, had ASI estimates below 97% (Appendix Table 18). The primers targeting archaea enabled us to detect 3232 two-on-two relationships between 340 different pairs of archaeal species with an ASI≥97% (Appendix Table 19). All primer pairs analyzed identified seven pairs of species (frequency= 20), which belonged to the genera *Methanobrevibacter* and *Methanocaldococcus*. There were 66 pairs of species detected only once by only one primer pair (frequency= 1). Again, most of the relationships were between archaeal species from the same genera (2359, 72.99%), but 873 (27.01%) involved taxa from distinct genera (Appendix Table 20). The combination of species from *Pyrococcus* and *Thermococcus* occurred the most, by far, (frequency= 428). The pairs of archaea with an ASI≥97%, such as Desulfurococcaceae and Pyrodictiaceae (frequency= 27), also belonged to distinct families (35; 1.08%), orders (3; 0.09%), or even classes (1; 0.03%) (Appendices Tables 21-23).

## DISCUSSION

The high degree of similarity between full-length 16S rRNA sequences from distinct species, or even genera, has been reported in the literature (Schloss 2021; Vĕtrovský and Baldrian 2013), leading to questions about the reliability of diversity estimates based on sequence-clustering methods. About full-length genes and the conventionally used 97% sequence-similarity threshold, some authors have detected that around a quarter of constructed OTUs contain sequences from multiple species (Schloss 2021; Vĕtrovský and Baldrian 2013) and about a tenth from distinct genera (Vĕtrovský and Baldrian 2013). These estimates were obviously higher when gene regions were assessed instead of full sequences. Schloss et al. (Schloss 2021) found that, with a 97% similarity threshold and applying the OptiClust algorithm, 31.7%, 34.3% and 34.8% of the OTUs assessed had 16S rRNA amplicons from distinct species in the V3-V4, V4, and V4-V5 regions, respectively (Schloss 2021). However, these investigations did not focus on taxa inhabiting a specific environment, despite the importance of conducting 16S rRNA gene-based research using habitat-specific databases (Escapa et al. 2020). Consequently, we used primer pairs targeting several gene regions (Regueira-Iglesias et al. 2021.a) to determine the number of different oral-bacterial and oral-archaeal species with an ASI≥97%, as well as the potential clusters that might contain distinct species. Moreover, for the first time in this kind of analysis, we identified the specific taxa of the oral environment that can be erroneously grouped in the same cluster.

In the present study, the primer pairs that targeted bacteria had a mean of 91.88 (49.40%) bacterial species with an ASI≥97% and an average of 153.46 potential OTUs containing distinct species. For those targeting archaea, these numbers were 65.60 (48.59%) and 162.26, respectively. Using SC-NASI≥97% values as a selection criterion, the optimum primer pair for detecting oral bacteria was OP_F053-KP_R020. Despite it being the primer used most in the studies contained in the oral-microbiome literature, KP_F031-KP_R021 identified slightly fewer species with an ASI≥97% (37 vs. 58) and identities (32 vs. 46); its SC-NASI≥97% was also lower than that of OP_F053-KP_R020 (54.30% vs. 65.05%). The primer pair producing the best estimates for detecting oral archaea was KP_F018-KP_R002. Again, the widely used primer KP_F014-KP_R011, although it only detected a few species with an ASI≥97% (24 vs. 60) and identities (96 vs. 125), however, also had a considerably lower SC-NASI≥97% than that of KP_F018-KP_R002 (12.59% vs. 51.11%). Lastly, we recommend the primer OP_F114-KP_R031 for detecting oral bacteria and archaea simultaneously. OP_F114-KP_R002, meanwhile, identified slightly fewer taxa with an ASI≥97% (144 vs. 147) and identities (244 vs. 269) but had a lower SC-NASI≥97% (50.47% vs. 52.02%). In addition, as previously observed (Regueira-Iglesias et al. 2021.a; Regueira-Iglesias et al. 2021.b), none of the primer combinations that are most commonly employed in sequencing-based studies of the oral microbiome were among the best. Specifically, the species coverage of KP_F078-OP_R010, a primer described by Caporaso (Caporaso et al. 2011), fell from 81.62% (Regueira-Iglesias et al 2021.b) to 33.33% when considering the species with an ASI≥97%, possibly generating as many as 314 potential clusters that contain different species.

Around 80% of the oral-bacteria and oral-archaea species analyzed had an ASI≥97% with at least another species. The widely-known bacterial periodontopathogens *Fusobacterium nucleatum* and *Treponema denticola* (Teles et al. 2013) had similar amplicons to *Fusobacterium hwasookii* and *Treponema putidum*, respectively, which have also been detected in periodontal lesions (Cho et al. 2015; Wyss et al. 2004). Interestingly, other bacteria with high amplicon similarities had antagonistic roles. Examples are: the health-associated *Campylobacter concisus* and the initially periodontitis-associated *Campylobacter curvus* (Henne et al. 2014); the health-related *Rothia mucilaginosa* (Zhang et al. 2018) and the decay-abundant *Rothia dentocariosa* (Jiang et al. 2016); the commensal *Streptococcus mitis*, *oralis*, and *salivarius*; the caries-associated *Streptococcus mutans* (Abranches et al. 2018; Teles et al. 2013); and the periodontal health-related *Tannerella sp. oral taxon HOT-286* (Vartoukian et al. 2016) and the periodontitis-related *Tannerella forsythia* (Teles et al. 2013). Furthermore, relevant oral-disease associated species, such as *Aggregatibacter actinomycetemcomitans* (Teles et al. 2013) and *Rothia dentocariosa* (Jiang et al. 2016), were among those that had an ASI≥97% with taxa from distinct genera. Regarding the archaea, we found that four Methanosarcina species found in healthy and periodontitis pockets, namely *barkeri*, *lacustris*, *mazeii*, and *vacuolata* (Deng et al. 2017), were highly similar. Moreover, *Halovivax ruber*, *Methanotorris igneus*, *Methanosalsum zhilinae*, and *Natronococcus occultus*, which are reported to be among the 10 most abundant species in both healthy and periodontitis subjects (Deng et al. 2017), had an ASI≥97% with several taxa from distinct genera.

Although Schloss (Schloss 2021) has recently stated that the risks of splitting a genome into multiple OTUs are greater than those of clustering species together, our clinical view is that the latter approach should be avoided. Amplicons from species traditionally associated with contrary health conditions, like those described above, can be grouped with a ≥97% identity threshold. This would, however, result in both an overabundance of the single species representing the cluster and an underestimation of the diversity of the community, with other species within the OTU overlooked. Consequently, despite possible difficulties, it would be better to use the lowest possible level of resolution, i.e., the variant level (Callahan et al. 2017), and databases specifically designed for taxonomic identifications of taxa at this level (Escapa et al. 2020).

The main limitation of our study is that the different ASVs from a given species can simultaneously present with very similar values to those of a number of other species, while distinct OTU clustering approaches, or even the same method, can yield uneven results for the same dataset (He et al. 2015; Wei et al. 2021; Westcott and Schloss 2015). Consequently, the results presented here are an approximation of the different oral species that could be grouped in OTUs. Furthermore, we only utilized one of all the possible similarity values ≥97% relating to ASVs from two particular species and discarded the others. Another consideration is that we were only able to evaluate 25% of the oral microorganism genomes listed on the eHOMD website, as the remainder were not fully sequenced. This absence of complete genomes reduced the number of species investigated to 35% of those set out on the site. Although the analysis could have been performed on annotations of the 16S rRNA gene sequences from oral microbes, we preferred to use complete genomes, thereby ensuring the high quality of the sequences reviewed. Thus, our results highlight only part of a much more extensive problem.

In conclusion, the tested primer pairs targeting bacteria and/or archaea identified an average of more than 150 potential OTUs that might contain different species. According to the SC-NASI≥97%, the best primer pairs were: OP_F053-KP_R020 for bacteria (region 1-3; primer pair position for *Escherichia coli* J01859.1: 9-356); KP_F018-KP_R002 for archaea (3; undefined-532); and OP_F114-KP_R031 for both (3-5; 340-801). Around 80% of the oral-bacteria and oral-archaea species analyzed had an ASI≥97% with at least one other taxa. Of these species, *Campylobacter*, *Rothia*, *Streptococcus*, and *Tannerella* play contrasting roles in the oral microbiota. Moreover, ~20% and ~30% of these two-by-two similarity relationships were established between species from different bacterial and archaeal genera, respectively. Even taxa from distinct families, orders, and classes could be grouped in the same OTU. Consequently, irrespective of the primer pair used, OTUs constructed with a 97% similarity provide an inaccurate description of oral-bacterial and oral-archaeal species, greatly affecting microbial diversity parameters. As a result, clustering by OTUs impacts the credibility of the associations between some oral species and certain health and disease conditions. This limits significantly the comparability of the microbial_diversity findings reported in oral microbiome literature.

## ACKNOWLEDGMENTS

This investigation was supported by the Instituto de Salud Carlos III (General Division of Evaluation and Research Promotion, Madrid, Spain) and co-financed by the FEDER (European Regional Development Fund, ERDF) (“A way of making Europe”) under grant ISCIII/PI17/01722; the Consellería de Cultura, Educación e Ordenación Universitaria de la Xunta de Galicia (accreditation 2019-2022 ED431G-2019/04, group with growth potential ED431B 2020-2022 GPC2020/27; A. Regueira-Iglesias support ED481A-2017/233) and the ERDF, which acknowledges the CiTIUS-Research Center in Intelligent Technologies of the Santiago de Compostela University as a Research Center of the Galician University System.

The funders had no role in study design, data collection and analysis, decision to publish, or preparation of the manuscript.

## AUTHOR CONTRIBUTIONS

Balsa-Castro C, Carreira MJ, and Tomás I contributed to the conception and design of the study, and critically revised manuscript. Regueira-Iglesias A, Vázquez-González L, Blanco-Pintos T, and Arce VM contributed to acquisition, analysis, and interpretation, and drafted the manuscript. All the authors gave final approval and agree to be accountable for all aspects of the work in ensuring that questions relatingto the accuracy or integrity of any part of the work are appropriately investigated and resolved.

