## Supplemental Table 1-23 for "OTUs clustering should be avoided for defining oral microbiome"

**TITLE:** OTU clustering should be avoided for defining oral microbiome

**Appendices**

Appendix Table 1. Selected primer pairs with high *in silico* coverage values targeting oral bacteria and/or archaea.

Appendix Table 2. Primer pairs are most used in the sequencing-based studies of the oral microbiome.

Appendix Table 3. Taxonomy classification, species identifier, and mean number of intragenomic 16S rRNA genes of the 186 oral bacteria species analyzed in the present study.

Appendix Table 4. Taxonomy classification, species identifier, and mean number of intragenomic 16S rRNA genes of the 135 oral archaea species analyzed in the present study.

Appendix Table 5. Primer pairs that produced the maximum number of identities/species ≥10 in the oral bacteria genomes.

Appendix Table 6. Primer pairs that produced the maximum number of identities/species ≥10 in the oral archaea genomes.

Appendix Table 7. Number of species with amplicon similarity values ≥97%, number of identities, and SC and SC-NASI≥97% values obtained by each primer pair analyzed against both domains.

Appendix Table 8. Taxonomy of all different bacterial taxa with a maximum number of identities/species ≥10.

Appendix Table 9. Taxonomy of all different archaeal taxa with a maximum number of identities/species ≥10.

Appendix Table 10. Oral-bacteria species with amplicon similarity values ≥97% with at least one different taxon.

Appendix Table 11. Oral-bacteria species with no amplicon similarity values ≥97% with different taxa.

Appendix Table 12. Pairs of bacterial species with amplicon similarity ≥97% using the analyzed primer pairs.

Appendix Table 13. Genus of the pairs of bacterial species with amplicon similarity ≥97%.

Appendix Table 14. Families of the pairs of bacterial species with amplicon similarity ≥97%.

Appendix Table 15. Orders of the pairs of bacterial species with amplicon similarity ≥97%.

Appendix Table 16. Classes of the pairs of bacterial species with amplicon similarity ≥97%.

Appendix Table 17. Oral-archaea species with amplicon similarity values ≥97% with at least one different taxon.

Appendix Table 18. Oral-archaea species with no amplicon similarity values ≥97% with different taxa.

Appendix Table 19. Pairs of archaeal species with amplicon similarity ≥97% using the analyzed primer pairs.

Appendix Table 20. Genus of the pairs of archaeal species with amplicon similarity ≥97%.

Appendix Table 21. Families of the pairs of archaeal species with amplicon similarity ≥97%.

Appendix Table 22. Orders of the pairs of archaeal species with amplicon similarity ≥97%.

Appendix Table 23. Classes of the pairs of archaeal species with amplicon similarity ≥97%.

Appendix Table 1. Selected primer pairs with high *in silico* coverage values targeting oral bacteria and/or archaea.

| **ALC** | **Bacterial-specific primer pair** | **F identifier** | **F Sequence 5-3** | **F First post** | **F Last Post** | **R identifier** | **R Sequence 5-3** | **R First post** | **R Last Post** | **Length (bps)** |
| --- | --- | --- | --- | --- | --- | --- | --- | --- | --- | --- |
| **S** | KP_F048-OP_R043 | KP_F048 | TACGGRAGGCAGCAG | 342 | 356 | OP_R043 | CCGCGRCTGCTGGCAC | 514 | 529 | 187 |
|  | OP_F098-OP_R119 | OP_F098 | CCAGCAGCYGCGGTAAN | 517 | 533 | OP_R119 | GGACTACCRGGGTATCTAA | 787 | 805 | 288 |
|  | OP_F066-KP_R040 | OP_F066 | GGMTTAGATACCC | 784 | 796 | KP_R040 | CCGTCAATTCMTTTGAGTTT | 906 | 925 | 141 |
|  | OP_F009-OP_R030 | OP_F009 | GGATTAGATACCCBRGTAGTC | 784 | 804 | OP_R030 | TCACRRCACGAGCTGWCGAC | 1060 | 1079 | 295 |
|  | KP_F061-KP_R074 | KP_F061 | ACTCAAAKGAATWGACGG | 908 | 925 | KP_R074 | GGGTYKCGCTCGTTR | 1099 | 1113 | 205 |
|  | OP_F101-OP_R030 | OP_F101 | GAATTGRCGGGGRCC | 916 | 930 | OP_R030 | TCACRRCACGAGCTGWCGAC | 1060 | 1079 | 163 |
| **M** | OP_F053-KP_R020 | OP_F053 | GRGTTYGATYMTGGCTCAG | 9 | 27 | KP_R020 | CTGCTGCCTYCCGTA | 342 | 356 | 347 |
|  | KP_F048-KP_R031 | KP_F048 | TACGGRAGGCAGCAG | 342 | 356 | KP_R031 | TACHVGGGTATCTAAKCC | 784 | 801 | 459 |
|  | KP_F048-OP_R073 | KP_F048 | TACGGRAGGCAGCAG | 342 | 356 | OP_R073 | CRTACTHCHCAGGYG | 879 | 893 | 551 |
|  | KP_F051-KP_R041 | KP_F051 | GTGCCAGCMGCNGCGG | 514 | 529 | KP_R041 | CGTCAATTCMTTTGAGTT | 907 | 924 | 410 |
|  | KP_F051-OP_R030 | KP_F051 | GTGCCAGCMGCNGCGG | 514 | 529 | OP_R030 | TCACRRCACGAGCTGWCGAC | 1060 | 1079 | 565 |
|  | OP_F116-KP_R060 | OP_F116 | YAACGAGCGCAACCC | 1099 | 1113 | KP_R060 | GACGGGCGGTGWGTRCA | 1390 | 1406 | 307 |
| **L** | KP_F048-OP_R030 | KP_F048 | TACGGRAGGCAGCAG | 342 | 356 | OP_R030 | TCACRRCACGAGCTGWCGAC | 1060 | 1079 | 737 |
|  | KP_F048-KP_R060 | KP_F048 | TACGGRAGGCAGCAG | 342 | 356 | KP_R060 | GACGGGCGGTGWGTRCA | 1390 | 1406 | 1064 |
|  | KP_F056-KP_R077 | KP_F056 | AYTGGGYDTAAAGNG | 572 | 576 | KP_R077 | GACGGGCGGTGTGTACAA | 1389 | 1406 | 834 |
| **ALC** | **Archaeal-specific primer pair** | **F identifier** | **F Sequence 5-3** | **F First post** | **F Last Post** | **R identifier** | **R Sequence 5-3** | **R First post** | **R Last Post** | **Length (bps)** |
| **S** | KP_F018-KP_R002 | KP_F018 | GYGCASCAGKCGMGAAW | U | U | KP_R002 | TTACCGCGGCKGCTG | 518 | 532 | - |
|  | OP_F066-KP_R013 | OP_F066 | GGMTTAGATACCC | 784 | 796 | KP_R013 | GGCCATGCACCWCCTCTC | U | U | - |
| **M** | KP_F018-KP_R032 | KP_F018 | GYGCASCAGKCGMGAAW | U | U | KP_R032 | TACNVGGGTATCTAATCC | 784 | 801 | - |
|  | KP_F018-OP_R073 | KP_F018 | GYGCASCAGKCGMGAAW | U | U | OP_R073 | CRTACTHCHCAGGYG | 879 | 893 | - |
|  | KP_F020-KP_R013 | KP_F020 | CAGCMGCCGCGGTAA | 518 | 532 | KP_R013 | GGCCATGCACCWCCTCTC | U | U | - |
|  | KP_F022-KP_R063 | KP_F022 | AGGAATTGGCGGGGGAGCA | U | U | KP_R063 | TACCTTGTTACGACTT | 1491 | 1506 | - |
| **L** | OP_F114-KP_R013 | OP_F114 | CCTAYGGGRBGCASCAG | 340 | 356 | KP_R013 | GGCCATGCACCWCCTCTC | U | U | - |
|  | KP_F018-KP_R063 | KP_F018 | GYGCASCAGKCGMGAAW | U | U | KP_R063 | TACCTTGTTACGACTT | 1491 | 1506 | - |
|  | OP_F066-OP_R016 | OP_F066 | GGMTTAGATACCC | 784 | 796 | OP_R016 | CGGTGTGTGCAAGGAG | U | U | - |
| **ALC** | **Bacterial and archaeal primer pair** | **F identifier** | **F Sequence 5-3** | **F First post** | **F Last Post** | **R identifier** | **R Sequence 5-3** | **R First post** | **R Last Post** | **Length (bps)** |
| S | OP_F114-KP_R002 | OP_F114 | CCTAYGGGRBGCASCAG | 340 | 356 | KP_R002 | TTACCGCGGCKGCTG | 518 | 532 | 192 |
|  | KP_F020-KP_R032 | KP_F020 | CAGCMGCCGCGGTAA | 518 | 532 | KP_R032 | TACNVGGGTATCTAATCC | 784 | 801 | 283 |
|  | OP_F066-OP_R073 | OP_F066 | GGMTTAGATACCC | 784 | 796 | OP_R073 | CRTACTHCHCAGGYG | 879 | 893 | 109 |
| M | OP_F114-KP_R031 | OP_F114 | CCTAYGGGRBGCASCAG | 340 | 356 | KP_R031 | TACHVGGGTATCTAAKCC | 784 | 801 | 461 |
|  | OP_F114-OP_R073 | OP_F114 | CCTAYGGGRBGCASCAG | 340 | 356 | OP_R073 | CRTACTHCHCAGGYG | 879 | 893 | 553 |
|  | KP_F020-OP_R073 | KP_F020 | CAGCMGCCGCGGTAA | 518 | 532 | OP_R073 | CRTACTHCHCAGGYG | 879 | 893 | 375 |
| L | OP_F114-OP_R121 | OP_F114 | CCTAYGGGRBGCASCAG | 340 | 356 | OP_R121 | ACGGGCGGTGWGTRC | 1391 | 1405 | 1065 |
|  | KP_F020-OP_R121 | KP_F020 | CAGCMGCCGCGGTAA | 518 | 532 | OP_R121 | ACGGGCGGTGWGTRC | 1391 | 1405 | 887 |
|  | OP_F066-OP_R121 | OP_F066 | GGMTTAGATACCC | 784 | 796 | OP_R121 | ACGGGCGGTGWGTRC | 1391 | 1405 | 621 |

Primer pairs were selected based on the species coverage values (number of species detected /total species evaluated) in a previous investigation (15). They were individually evaluated through regular expressions against *Escherichia coli* J01859 to define their positions. The U values represent a mismatch on the assessment and, therefore, the position cannot be confirmed with a guarantee. ALC= amplicon length category; bps= base pairs; F= forward; KP= Klindworth primer; L= long mean amplicon length category, >600 base pairs; M= medium mean amplicon length category, 301-600 base pairs; OP= oral primer; Post= position; R= reverse; S= short mean amplicon length category, 100-300 base pairs.

Appendix Table 2. Primer pairs are most used in the sequencing-based studies of the oral microbiome.

| **ALC** | **Most used primer pair** | **F identifier** | **F Sequence 5-3** | **F First post** | **F Last Post** | **R identifier** | **R Sequence 5-3** | **R First post** | **R Last Post** | **Length (bps)** |
| --- | --- | --- | --- | --- | --- | --- | --- | --- | --- | --- |
| S | KP_F078-OP_R010^B+A^ | KP_F078 | GTGCCAGCMGCCGCGGTAA | 514 | 532 | OP_R010 | GGACTACHVGGGTWTCTAAT | 786 | 805 | 291 |
| M | KP_F031-KP_R021^B^ | KP_F031 | AGAGTTTGATCCTGGCTCAG | 8 | 27 | KP_R021 | TTACCGCGGCTGCTGGCAC | 515 | 532 | 524 |
|  | KP_F047_KP_R035^B^ | KP_F047 | CCTACGGGNGGCWGCAG | 340 | 356 | KP_R035 | GACTACHVGGGTATCTAATCC | 784 | 804 | 464 |
|  | OP_F009_OP_R029^B^ | OP_F009 | GGATTAGATACCCBRGTAGTC | 784 | 868 | OP_R029 | ACGTCRTCCCCDCCTTCCTC | 1174 | 1193 | 409 |
| L | KP_F014_KP_R011^A^ | KP_F014 | TCCAGGCCCTACGGG | U | U | KP_R011 | YCCGGCGTTGAMTCCAATT | U | U | - |
|  | KP_F034_KP_R065^B^ | KP_F034 | AGAGTTTGATCMTGGCTCAG | 8 | 27 | KP_R065 | TACGGYTACCTTGTTACGACTT | 1491 | 1512 | 1504 |

Primer pairs were selected based on the species coverage values (number of species detected /total species evaluated) in a previous investigation (15). They were individually evaluated through regular expressions against *Escherichia coli* J01859 to define their positions. The U values represent a mismatch on the assessment and, therefore, the position cannot be confirmed with a guarantee. The target domain of each primer pair is indicated as a superscript. A= archaea; ALC= amplicon length category; B= bacteria; bps= base pairs; F= forward; KP= Klindworth primer; L= long mean amplicon length category, >600 base pairs; M= medium mean amplicon length category, 301-600 base pairs; OP= oral primer; Post= position; R= reverse; S= short mean amplicon length category, 100-300 base pairs; U= unidentified; WUOL= widely used in the oral literature.

Appendix Table 3. Taxonomy classification, species identifier and mean number of intragenomic 16S rRNA genes of the 186 oral bacteria species analyzed in the present study.

| **ID** | **Num genes**  **(mean)** | **Phylum** | **Class** | **Order** | **Family** | **Genus** | **Species** |
| --- | --- | --- | --- | --- | --- | --- | --- |
| SP00136 | 7.00 | Proteobacteria | Gammaproteobacteria | Enterobacterales | Enterobacteriaceae | Escherichia | coli |
| SP00137 | 2.00 | Proteobacteria | Epsilonproteobacteria | Campylobacterales | Helicobacteraceae | Helicobacter | pylori |
| SP00138 | 1.00 | Actinobacteria | Actinomycetia | Corynebacteriales | Mycobacteriaceae | Mycobacterium | tuberculosis |
| SP00139 | 2.00 | Spirochaetes | Spirochaetia | Spirochaetales | Spirochaetaceae | Treponema | pallidum |
| SP00140 | 1.00 | Chlamydiae | Chlamydiia | Chlamydiales | Chlamydiaceae | Chlamydia | pneumoniae |
| SP00141 | 4.00 | Proteobacteria | Betaproteobacteria | Neisseriales | Neisseriaceae | Neisseria | meningitidis |
| SP00142 | 4.00 | Proteobacteria | Gammaproteobacteria | Pseudomonadales | Pseudomonadaceae | Pseudomonas | aeruginosa |
| SP00143 | 6.00 | Firmicutes | Bacilli | Lactobacillales | Streptococcaceae | Streptococcus | pyogenes |
| SP00144 | 4.00 | Proteobacteria | Betaproteobacteria | Neisseriales | Neisseriaceae | Neisseria | gonorrhoeae |
| SP00145 | 6.00 | Firmicutes | Bacilli | Lactobacillales | Streptococcaceae | Lactococcus | lactis |
| SP00146 | 4.00 | Firmicutes | Bacilli | Lactobacillales | Streptococcaceae | Streptococcus | pneumoniae |
| SP00147 | 4.00 | Proteobacteria | Alphaproteobacteria | Hyphomicrobiales | Rhizobiaceae | Agrobacterium | fabrum |
| SP00148 | 7.00 | Firmicutes | Bacilli | Lactobacillales | Streptococcaceae | Streptococcus | agalactiae |
| SP00149 | 4.00 | Fusobacteria | Fusobacteriia | Fusobacteriales | Fusobacteriaceae | Fusobacterium | nucleatum |
| SP00150 | 7.00 | Proteobacteria | Gammaproteobacteria | Enterobacterales | Yersiniaceae | Yersinia | pestis |
| SP00151 | 5.00 | Firmicutes | Bacilli | Lactobacillales | Streptococcaceae | Streptococcus | mutans |
| SP00152 | 4.00 | Actinobacteria | Actinomycetia | Bifidobacteriales | Bifidobacteriaceae | Bifidobacterium | longum |
| SP00153 | 4.00 | Bacteroidetes | Bacteroidia | Bacteroidales | Porphyromonadaceae | Porphyromonas | gingivalis |
| SP00154 | 6.00 | Firmicutes | Bacilli | Bacillales | Staphylococcaceae | Staphylococcus | epidermidis |
| SP00155 | 4.00 | Firmicutes | Bacilli | Lactobacillales | Enterococcaceae | Enterococcus | faecalis |
| SP00156 | 11.00 | Firmicutes | Bacilli | Bacillales | Bacillaceae | Bacillus | anthracis |
| SP00157 | 6.00 | Proteobacteria | Gammaproteobacteria | Pasteurellales | Pasteurellaceae | Haemophilus | ducreyi |
| SP00158 | 5.00 | Firmicutes | Bacilli | Lactobacillales | Lactobacillaceae | Lactobacillus | johnsonii |
| SP00159 | 2.00 | Spirochaetes | Spirochaetia | Spirochaetales | Spirochaetaceae | Treponema | denticola |
| SP00160 | 6.00 | Firmicutes | Bacilli | Bacillales | Listeriaceae | Listeria | monocytogenes |
| SP00161 | 3.00 | Actinobacteria | Actinomycetia | Propionibacteriales | Propionibacteriaceae | Cutibacterium | acnes |
| SP00162 | 7.00 | Firmicutes | Bacilli | Lactobacillales | Lactobacillaceae | Ligilactobacillus | salivarius |
| SP00163 | 5.00 | Firmicutes | Bacilli | Bacillales | Staphylococcaceae | Staphylococcus | aureus |
| SP00164 | 1.00 | Actinobacteria | Actinomycetia | Corynebacteriales | Mycobacteriaceae | Mycobacterium | leprae |
| SP00165 | 5.00 | Firmicutes | Bacilli | Lactobacillales | Lactobacillaceae | Lactiplantibacillus | plantarum |
| SP00166 | 6.00 | Proteobacteria | Gammaproteobacteria | Pseudomonadales | Pseudomonadaceae | Pseudomonas | fluorescens |
| SP00167 | 4.00 | Proteobacteria | Gammaproteobacteria | Xanthomonadales | Xanthomonadaceae | Stenotrophomonas | maltophilia |
| SP00168 | 3.00 | Actinobacteria | Actinomycetia | Corynebacteriales | Corynebacteriaceae | Corynebacterium | urealyticum |
| SP00169 | 7.00 | Proteobacteria | Gammaproteobacteria | Enterobacterales | Morganellaceae | Proteus | mirabilis |
| SP00170 | 2.00 | Proteobacteria | Alphaproteobacteria | Hyphomicrobiales | Phyllobacteriaceae | Mesorhizobium | japonicum |
| SP00171 | 7.00 | Firmicutes | Bacilli | Bacillales | Bacillaceae | Alkalihalobacillus | clausii |
| SP00172 | 8.00 | Proteobacteria | Gammaproteobacteria | Enterobacterales | Enterobacteriaceae | Klebsiella | pneumoniae |
| SP00173 | 6.00 | Firmicutes | Bacilli | Lactobacillales | Lactobacillaceae | Limosilactobacillus | reuteri |
| SP00174 | 5.00 | Firmicutes | Bacilli | Lactobacillales | Lactobacillaceae | Limosilactobacillus | fermentum |
| SP00175 | 4.00 | Firmicutes | Tissierellia | Tissierellales | Peptoniphilaceae | Finegoldia | magna |
| SP00176 | 2.00 | Tenericutes | Mollicutes | Mycoplasmatales | Mycoplasmataceae | Mycoplasmopsis | fermentans |
| SP00177 | 4.00 | Firmicutes | Bacilli | Lactobacillales | Streptococcaceae | Streptococcus | intermedius |
| SP00178 | 3.00 | Actinobacteria | Actinomycetia | Micrococcales | Micrococcaceae | Rothia | mucilaginosa |
| SP00179 | 9.00 | Firmicutes | Bacilli | Bacillales | Bacillaceae | Bacillus | subtilis |
| SP00180 | 2.00 | Chloroflexi | Anaerolineae | Anaerolineales | Anaerolineaceae | Anaerolinea | thermophila |
| SP00181 | 5.00 | Firmicutes | Bacilli | Lactobacillales | Lactobacillaceae | Levilactobacillus | brevis |
| SP00182 | 1.00 | Tenericutes | Mollicutes | Mycoplasmatales | Mycoplasmataceae | Mycoplasma | pneumoniae |
| SP00183 | 2.00 | Chloroflexi | Caldilineae | Caldilineales | Caldilineaceae | Caldilinea | aerophila |
| SP00184 | 4.00 | Firmicutes | Bacilli | Lactobacillales | Streptococcaceae | Streptococcus | anginosus |
| SP00185 | 6.00 | Proteobacteria | Gammaproteobacteria | Pasteurellales | Pasteurellaceae | Aggregatibacter | actinomycetemcomitans |
| SP00186 | 5.00 | Firmicutes | Bacilli | Bacillales | Staphylococcaceae | Staphylococcus | schleiferi |
| SP00187 | 5.00 | Actinobacteria | Actinomycetia | Corynebacteriales | Corynebacteriaceae | Corynebacterium | diphtheriae |
| SP00188 | 3.00 | Proteobacteria | Betaproteobacteria | Burkholderiales | Alcaligenaceae | Bordetella | pertussis |
| SP00189 | 4.00 | Firmicutes | Bacilli | Lactobacillales | Lactobacillaceae | Lactobacillus | acidophilus |
| SP00190 | 6.00 | Proteobacteria | Gammaproteobacteria | Pasteurellales | Pasteurellaceae | Haemophilus | influenzae |
| SP00191 | 5.00 | Proteobacteria | Gammaproteobacteria | Pseudomonadales | Pseudomonadaceae | Pseudomonas | protegens |
| SP00192 | 3.00 | Actinobacteria | Actinomycetia | Bifidobacteriales | Bifidobacteriaceae | Bifidobacterium | breve |
| SP00193 | 4.00 | Proteobacteria | Gammaproteobacteria | Pseudomonadales | Pseudomonadaceae | Pseudomonas | stutzeri |
| SP00194 | 4.00 | Firmicutes | Bacilli | Lactobacillales | Streptococcaceae | Streptococcus | sanguinis |
| SP00195 | 6.00 | Firmicutes | Bacilli | Lactobacillales | Lactobacillaceae | Lactobacillus | gasseri |
| SP00196 | 5.00 | Firmicutes | Bacilli | Lactobacillales | Lactobacillaceae | Lacticaseibacillus | paracasei |
| SP00197 | 6.00 | Proteobacteria | Gammaproteobacteria | Pseudomonadales | Moraxellaceae | Acinetobacter | baumannii |
| SP00198 | 3.00 | Proteobacteria | Alphaproteobacteria | Hyphomicrobiales | Rhizobiaceae | Agrobacterium | radiobacter |
| SP00199 | 4.00 | Firmicutes | Bacilli | Lactobacillales | Streptococcaceae | Streptococcus | gordonii |
| SP00200 | 4.00 | Proteobacteria | Alphaproteobacteria | Hyphomicrobiales | Brucellaceae | Brucella | anthropi |
| SP00201 | 3.00 | Proteobacteria | Epsilonproteobacteria | Campylobacterales | Campylobacteraceae | Campylobacter | curvus |
| SP00202 | 7.00 | Proteobacteria | Gammaproteobacteria | Enterobacterales | Enterobacteriaceae | Cronobacter | sakazakii |
| SP00203 | 3.00 | Proteobacteria | Epsilonproteobacteria | Campylobacterales | Campylobacteraceae | Campylobacter | concisus |
| SP00204 | 5.00 | Proteobacteria | Betaproteobacteria | Burkholderiales | Comamonadaceae | Delftia | acidovorans |
| SP00205 | 8.00 | Proteobacteria | Gammaproteobacteria | Enterobacterales | Enterobacteriaceae | Klebsiella | variicola |
| SP00206 | 4.00 | Proteobacteria | Betaproteobacteria | Burkholderiales | Burkholderiaceae | Ralstonia | pickettii |
| SP00207 | 2.00 | Chlorobi | Chlorobia | Chlorobiales | Chlorobiaceae | Chlorobium | limicola |
| SP00208 | 4.00 | Actinobacteria | Actinomycetia | Bifidobacteriales | Bifidobacteriaceae | Bifidobacterium | animalis |
| SP00209 | 3.00 | Proteobacteria | Betaproteobacteria | Burkholderiales | Comamonadaceae | Comamonas | thiooxydans |
| SP00210 | 4.00 | Proteobacteria | Alphaproteobacteria | Rhodobacterales | Rhodobacteraceae | Rhodobacter | capsulatus |
| SP00211 | 3.00 | Proteobacteria | Betaproteobacteria | Burkholderiales | Comamonadaceae | Acidovorax | ebreus |
| SP00212 | 6.00 | Proteobacteria | Gammaproteobacteria | Pasteurellales | Pasteurellaceae | Aggregatibacter | aphrophilus |
| SP00213 | 3.00 | Actinobacteria | Actinomycetia | Corynebacteriales | Corynebacteriaceae | Corynebacterium | kroppenstedtii |
| SP00214 | 1.00 | Actinobacteria | Actinomycetia | Micrococcales | Micrococcaceae | Micrococcus | luteus |
| SP00215 | 4.00 | Bacteroidetes | Flavobacteriia | Flavobacteriales | Flavobacteriaceae | Capnocytophaga | ochracea |
| SP00216 | 2.00 | Proteobacteria | Betaproteobacteria | Burkholderiales | Comamonadaceae | Variovorax | paradoxus |
| SP00217 | 3.00 | Actinobacteria | Coriobacteriia | Eggerthellales | Eggerthellaceae | Cryptobacterium | curtum |
| SP00218 | 5.00 | Fusobacteria | Fusobacteriia | Fusobacteriales | Leptotrichiaceae | Leptotrichia | buccalis |
| SP00219 | 2.00 | Actinobacteria | Actinomycetia | Micrococcales | Kytococcaceae | Kytococcus | sedentarius |
| SP00220 | 4.00 | Firmicutes | Tissierellia | Tissierellales | Peptoniphilaceae | Anaerococcus | prevotii |
| SP00221 | 1.00 | Actinobacteria | Coriobacteriia | Coriobacteriales | Atopobiaceae | Lancefieldella | parvulum |
| SP00222 | 3.00 | Actinobacteria | Coriobacteriia | Eggerthellales | Eggerthellaceae | Eggerthella | lenta |
| SP00223 | 4.00 | Actinobacteria | Actinomycetia | Bifidobacteriales | Bifidobacteriaceae | Bifidobacterium | dentium |
| SP00224 | 4.00 | Actinobacteria | Actinomycetia | Micrococcales | Sanguibacteraceae | Sanguibacter | keddieii |
| SP00225 | 4.00 | Firmicutes | Negativicutes | Veillonellales | Veillonellaceae | Veillonella | parvula |
| SP00226 | 2.00 | Actinobacteria | Actinomycetia | Bifidobacteriales | Bifidobacteriaceae | Gardnerella | vaginalis |
| SP00227 | 4.00 | Proteobacteria | Gammaproteobacteria | Pseudomonadales | Moraxellaceae | Moraxella | catarrhalis |
| SP00228 | 4.00 | Actinobacteria | Actinomycetia | Actinomycetales | Actinomycetaceae | Arcanobacterium | haemolyticum |
| SP00229 | 1.00 | Actinobacteria | Coriobacteriia | Coriobacteriales | Atopobiaceae | Olsenella | uli |
| SP00230 | 4.00 | Bacteroidetes | Bacteroidia | Bacteroidales | Prevotellaceae | Prevotella | melaninogenica |
| SP00231 | 5.00 | Firmicutes | Clostridia | Eubacteriales | Eubacteriaceae | Eubacterium | callanderi |
| SP00232 | 3.00 | Actinobacteria | Actinomycetia | Micrococcales | Micrococcaceae | Rothia | dentocariosa |
| SP00233 | 3.00 | Proteobacteria | Betaproteobacteria | Burkholderiales | Alcaligenaceae | Achromobacter | xylosoxidans |
| SP00234 | 4.00 | Firmicutes | Clostridia | Eubacteriales | Peptostreptococcaceae | Filifactor | alocis |
| SP00235 | 4.00 | Firmicutes | Bacilli | Lactobacillales | Lactobacillaceae | Lactobacillus | amylovorus |
| SP00236 | 4.00 | Bacteroidetes | Bacteroidia | Bacteroidales | Prevotellaceae | Prevotella | denticola |
| SP00237 | 5.00 | Firmicutes | Bacilli | Lactobacillales | Lactobacillaceae | Lentilactobacillus | buchneri |
| SP00238 | 2.00 | Bacteroidetes | Bacteroidia | Bacteroidales | Porphyromonadaceae | Porphyromonas | asaccharolytica |
| SP00239 | 2.00 | Actinobacteria | Actinomycetia | Propionibacteriales | Propionibacteriaceae | Pseudopropionibacterium | propionicum |
| SP00240 | 8.00 | Proteobacteria | Gammaproteobacteria | Enterobacterales | Enterobacteriaceae | Klebsiella | aerogenes |
| SP00241 | 4.00 | Firmicutes | Bacilli | Lactobacillales | Streptococcaceae | Streptococcus | parasanguinis |
| SP00242 | 6.00 | Firmicutes | Bacilli | Lactobacillales | Streptococcaceae | Streptococcus | salivarius |
| SP00243 | 5.00 | Firmicutes | Bacilli | Lactobacillales | Lactobacillaceae | Lacticaseibacillus | rhamnosus |
| SP00244 | 2.00 | Bacteroidetes | Bacteroidia | Bacteroidales | Tannerellaceae | Tannerella | forsythia |
| SP00245 | 1.00 | Ignavibacteriae | Ignavibacteria | Ignavibacteriales | Ignavibacteriaceae | Ignavibacterium | album |
| SP00246 | 4.00 | Bacteroidetes | Bacteroidia | Bacteroidales | Prevotellaceae | Prevotella | intermedia |
| SP00247 | 1.00 | Ignavibacteriae | Ignavibacteria | Ignavibacteriales | Melioribacteraceae | Melioribacter | roseus |
| SP00248 | 5.00 | Firmicutes | Bacilli | Bacillales | Staphylococcaceae | Staphylococcus | warneri |
| SP00249 | 1.00 | Tenericutes | Mollicutes | Mycoplasmatales | Mycoplasmataceae | Mycoplasma | genitalium |
| SP00250 | 2.00 | Proteobacteria | Betaproteobacteria | Burkholderiales | Burkholderiaceae | Burkholderia | cepacia |
| SP00251 | 4.00 | Firmicutes | Bacilli | Lactobacillales | Streptococcaceae | Streptococcus | constellatus |
| SP00252 | 5.00 | Firmicutes | Bacilli | Bacillales | Staphylococcaceae | Staphylococcus | pasteuri |
| SP00253 | 4.00 | Firmicutes | Bacilli | Lactobacillales | Streptococcaceae | Streptococcus | cristatus |
| SP00254 | 3.00 | Actinobacteria | Actinomycetia | Propionibacteriales | Propionibacteriaceae | Cutibacterium | avidum |
| SP00255 | 7.00 | Proteobacteria | Gammaproteobacteria | Enterobacterales | Enterobacteriaceae | Cronobacter | malonaticus |
| SP00256 | 2.00 | Actinobacteria | Actinomycetia | Corynebacteriales | Mycobacteriaceae | Mycolicibacterium | neoaurum |
| SP00257 | 1.00 | C.Saccharibacteria | C.Saccharimonia | C.Nanosynbacterales | C.Nanosynbacteraceae | C.Nanosynbacter | lyticus |
| SP00258 | 6.00 | Firmicutes | Bacilli | Bacillales | Staphylococcaceae | Staphylococcus | capitis |
| SP00259 | 5.00 | Actinobacteria | Actinomycetia | Corynebacteriales | Corynebacteriaceae | Corynebacterium | sp. ATCC 6931 |
| SP00260 | 7.00 | Proteobacteria | Gammaproteobacteria | Pseudomonadales | Moraxellaceae | Acinetobacter | johnsonii |
| SP00261 | 4.00 | Proteobacteria | Betaproteobacteria | Burkholderiales | Burkholderiaceae | Cupriavidus | gilardii |
| SP00262 | 4.00 | Actinobacteria | Actinomycetia | Corynebacteriales | Corynebacteriaceae | Corynebacterium | singulare |
| SP00263 | 3.00 | Fusobacteria | Fusobacteriia | Fusobacteriales | Leptotrichiaceae | Sneathia | vaginalis |
| SP00264 | 2.00 | Actinobacteria | Coriobacteriia | Coriobacteriales | Atopobiaceae | Olsenella | sp. oral taxon 807 |
| SP00265 | 2.00 | Actinobacteria | Actinomycetia | Micrococcales | Intrasporangiaceae | Arsenicicoccus | sp. oral taxon 190 |
| SP00266 | 4.00 | Firmicutes | Negativicutes | Selenomonadales | Selenomonadaceae | Selenomonas | sp. oral taxon 478 |
| SP00267 | 3.00 | Actinobacteria | Actinomycetia | Actinomycetales | Actinomycetaceae | Schaalia | meyeri |
| SP00268 | 3.00 | Proteobacteria | Betaproteobacteria | Burkholderiales | Comamonadaceae | Ottowia | sp. oral taxon 894 |
| SP00269 | 3.00 | Bacteroidetes | Bacteroidia | Bacteroidales | Prevotellaceae | Prevotella | fusca |
| SP00270 | 2.00 | Actinobacteria | Actinomycetia | Corynebacteriales | Lawsonellaceae | Lawsonella | clevelandensis |
| SP00271 | 4.00 | Fusobacteria | Fusobacteriia | Fusobacteriales | Leptotrichiaceae | Leptotrichia | sp. oral taxon 212 |
| SP00272 | 3.00 | Actinobacteria | Actinomycetia | Micrococcales | Micrococcaceae | Kocuria | palustris |
| SP00273 | 4.00 | Bacteroidetes | Flavobacteriia | Flavobacteriales | Flavobacteriaceae | Capnocytophaga | sp. oral taxon 323 |
| SP00274 | 3.00 | Actinobacteria | Actinomycetia | Actinomycetales | Actinomycetaceae | Actinomyces | sp. oral taxon 414 |
| SP00275 | 7.00 | Proteobacteria | Gammaproteobacteria | Enterobacterales | Yersiniaceae | Serratia | marcescens |
| SP00276 | 3.00 | Bacteroidetes | Bacteroidia | Bacteroidales | Prevotellaceae | Prevotella | enoeca |
| SP00277 | 2.00 | Actinobacteria | Actinomycetia | Micrococcales | Intrasporangiaceae | Janibacter | indicus |
| SP00278 | 5.00 | Fusobacteria | Fusobacteriia | Fusobacteriales | Fusobacteriaceae | Fusobacterium | hwasookii |
| SP00279 | 6.00 | Firmicutes | Bacilli | Bacillales | Staphylococcaceae | Staphylococcus | haemolyticus |
| SP00280 | 6.00 | Firmicutes | Bacilli | Bacillales | Staphylococcaceae | Staphylococcus | lugdunensis |
| SP00281 | 7.00 | Bacteroidetes | Flavobacteriia | Flavobacteriales | Flavobacteriaceae | Capnocytophaga | haemolytica |
| SP00282 | 3.00 | Actinobacteria | Actinomycetia | Actinomycetales | Actinomycetaceae | Actinomyces | radicidentis |
| SP00283 | 3.00 | Proteobacteria | Deltaproteobacteria | Desulfovibrionales | Desulfovibrionaceae | Desulfovibrio | fairfieldensis |
| SP00284 | 2.00 | Proteobacteria | Deltaproteobacteria | Desulfovibrionales | Desulfomicrobiaceae | Desulfomicrobium | orale |
| SP00285 | 5.00 | Fusobacteria | Fusobacteriia | Fusobacteriales | Leptotrichiaceae | Leptotrichia | sp. oral taxon 847 |
| SP00286 | 3.00 | Actinobacteria | Actinomycetia | Actinomycetales | Actinomycetaceae | Actinomyces | oris |
| SP00287 | 4.00 | Proteobacteria | Gammaproteobacteria | Pseudomonadales | Moraxellaceae | Moraxella | osloensis |
| SP00288 | 4.00 | Firmicutes | Bacilli | Lactobacillales | Streptococcaceae | Streptococcus | sp. oral taxon 431 |
| SP00289 | 4.00 | Actinobacteria | Actinomycetia | Corynebacteriales | Corynebacteriaceae | Corynebacterium | simulans |
| SP00290 | 5.00 | Proteobacteria | Alphaproteobacteria | Rhodospirillales | Acetobacteraceae | Roseomonas | gilardii |
| SP00291 | 5.00 | Fusobacteria | Fusobacteriia | Fusobacteriales | Leptotrichiaceae | Leptotrichia | sp. oral taxon 498 |
| SP00292 | 5.00 | Firmicutes | Negativicutes | Veillonellales | Veillonellaceae | Dialister | pneumosintes |
| SP00293 | 2.00 | Bacteroidetes | Bacteroidia | Bacteroidales | Tannerellaceae | Tannerella | sp. oral taxon HOT-286 |
| SP00294 | 3.00 | Actinobacteria | Actinomycetia | Propionibacteriales | Propionibacteriaceae | Propionibacterium | sp. oral taxon 193 |
| SP00295 | 4.00 | Firmicutes | Negativicutes | Selenomonadales | Selenomonadaceae | Selenomonas | sp. oral taxon 920 |
| SP00296 | 6.00 | Proteobacteria | Gammaproteobacteria | Pseudomonadales | Moraxellaceae | Acinetobacter | junii |
| SP00297 | 6.00 | Firmicutes | Bacilli | Bacillales | Staphylococcaceae | Staphylococcus | cohnii |
| SP00298 | 3.00 | Proteobacteria | Alphaproteobacteria | Rhodobacterales | Rhodobacteraceae | Paracoccus | yeei |
| SP00299 | 9.00 | Firmicutes | Bacilli | Bacillales | Bacillaceae | Anoxybacillus | flavithermus |
| SP00300 | 4.00 | Actinobacteria | Actinomycetia | Corynebacteriales | Corynebacteriaceae | Corynebacterium | striatum |
| SP00301 | 3.00 | Actinobacteria | Actinomycetia | Micrococcales | Micrococcaceae | Kocuria | rhizophila |
| SP00302 | 6.00 | Firmicutes | Bacilli | Bacillales | Staphylococcaceae | Staphylococcus | pettenkoferi |
| SP00303 | 2.00 | Firmicutes | Clostridia | Eubacteriales | Oscillospiraceae | Fastidiosipila | sanguinis |
| SP00304 | 2.00 | Firmicutes | Clostridia | Eubacteriales | Clostridiales Family XIII. I. Sedis | Mogibacterium | diversum |
| SP00305 | 4.00 | Bacteroidetes | Flavobacteriia | Flavobacteriales | Flavobacteriaceae | Capnocytophaga | sp. oral taxon 878 |
| SP00306 | 4.00 | Bacteroidetes | Bacteroidia | Bacteroidales | Bacteroidaceae | Bacteroides | zoogleoformans |
| SP00307 | 4.00 | Bacteroidetes | Bacteroidia | Bacteroidales | Bacteroidaceae | Bacteroides | heparinolyticus |
| SP00308 | 6.00 | Proteobacteria | Gammaproteobacteria | Pasteurellales | Pasteurellaceae | Haemophilus | sp. oral taxon 036 |
| SP00309 | 3.00 | Actinobacteria | Actinomycetia | Actinomycetales | Actinomycetaceae | Actinomyces | sp. oral taxon 897 |
| SP00310 | 4.00 | Actinobacteria | Actinomycetia | Corynebacteriales | Dietziaceae | Dietzia | sp. oral taxon 368 |
| SP00311 | 8.00 | Firmicutes | Bacilli | Bacillales | Paenibacillaceae | Paenibacillus | glucanolyticus |
| SP00312 | 3.00 | Actinobacteria | Actinomycetia | Actinomycetales | Actinomycetaceae | Actinomyces | sp. oral taxon 171 |
| SP00313 | 3.00 | Actinobacteria | Actinomycetia | Actinomycetales | Actinomycetaceae | Schaalia | odontolytica |
| SP00314 | 3.00 | Actinobacteria | Actinomycetia | Actinomycetales | Actinomycetaceae | Actinomyces | sp. oral taxon 169 |
| SP00315 | 4.00 | Firmicutes | Bacilli | Lactobacillales | Streptococcaceae | Streptococcus | mitis |
| SP00316 | 4.00 | Proteobacteria | Betaproteobacteria | Neisseriales | Neisseriaceae | Neisseria | lactamica |
| SP00317 | 2.00 | Tenericutes | Mollicutes | Mycoplasmatales | Mycoplasmataceae | Mycoplasma | hominis |
| SP00318 | 6.00 | Proteobacteria | Gammaproteobacteria | Pasteurellales | Pasteurellaceae | Haemophilus | parainfluenzae |
| SP00319 | 4.00 | Firmicutes | Bacilli | Lactobacillales | Streptococcaceae | Streptococcus | oralis |
| SP00320 | 6.00 | Firmicutes | Bacilli | Lactobacillales | Streptococcaceae | Streptococcus | thermophilus |
| SP00321 | 2.00 | Spirochaetes | Spirochaetia | Spirochaetales | Spirochaetaceae | Treponema | putidum |

The mean number of intragenomic 16S rRNA genes of these species were calculated in a previous investigation of our group (Regueira-Iglesias et al. 2021.b). C.= candidatus; I.= Incertae; ID= species identifier; Num= number; sp.= species.

Appendix Table 4. Taxonomy classification, species identifier and mean number of intragenomic 16S rRNA genes of the 135 oral archaea species analyzed in the present study.

| **ID** | **Num genes**  **(mean)** | **Phylum** | **Class** | **Order** | **Family** | **Genus** | **Species** |
| --- | --- | --- | --- | --- | --- | --- | --- |
| SP00001 | 1.00 | Crenarchaeota | Thermoprotei | Desulfurococcales | Desulfurococcaceae | Ignisphaera | aggregans |
| SP00002 | 3.00 | Euryarchaeota | Methanomicrobia | Methanosarcinales | Methanosarcinaceae | Methanolobus | psychrophilus |
| SP00003 | 1.00 | Crenarchaeota | Thermoprotei | Thermoproteales | Thermoproteaceae | Pyrobaculum | oguniense |
| SP00004 | 1.00 | Crenarchaeota | Thermoprotei | Desulfurococcales | Desulfurococcaceae | Aeropyrum | pernix |
| SP00005 | 2.00 | Euryarchaeota | Methanococci | Methanococcales | Methanocaldococcaceae | Methanocaldococcus | jannaschii |
| SP00006 | 1.00 | Euryarchaeota | Thermococci | Thermococcales | Thermococcaceae | Pyrococcus | horikoshii |
| SP00007 | 3.00 | Euryarchaeota | Methanococci | Methanococcales | Methanococcaceae | Methanococcus | maripaludis |
| SP00008 | 1.00 | Euryarchaeota | Halobacteria | Halobacteriales | Halobacteriaceae | Halobacterium | salinarum |
| SP00009 | 1.00 | C.Thermoplasmatota | Thermoplasmata | Thermoplasmatales | Thermoplasmataceae | Thermoplasma | volcanium |
| SP00010 | 1.00 | Crenarchaeota | Thermoprotei | Thermoproteales | Thermoproteaceae | Pyrobaculum | aerophilum |
| SP00011 | 1.00 | Euryarchaeota | Methanopyri | Methanopyrales | Methanopyraceae | Methanopyrus | kandleri |
| SP00012 | 3.00 | Euryarchaeota | Methanomicrobia | Methanosarcinales | Methanosarcinaceae | Methanosarcina | acetivorans |
| SP00013 | 3.00 | Euryarchaeota | Methanomicrobia | Methanosarcinales | Methanosarcinaceae | Methanosarcina | mazei |
| SP00014 | 1.00 | C.Thermoplasmatota | Thermoplasmata | Thermoplasmatales | Picrophilaceae | Picrophilus | torridus |
| SP00015 | 3.00 | Euryarchaeota | Halobacteria | Halobacteriales | Haloarculaceae | Haloarcula | marismortui |
| SP00016 | 2.00 | Crenarchaeota | Thermoprotei | Sulfolobales | Sulfolobaceae | Sulfolobus | acidocaldarius |
| SP00017 | 3.00 | Euryarchaeota | Methanomicrobia | Methanosarcinales | Methanosarcinaceae | Methanosarcina | barkeri |
| SP00018 | 1.00 | Euryarchaeota | Halobacteria | Halobacteriales | Haloarculaceae | Natronomonas | pharaonis |
| SP00019 | 4.00 | Euryarchaeota | Methanomicrobia | Methanomicrobiales | Methanospirillaceae | Methanospirillum | hungatei |
| SP00020 | 3.00 | Euryarchaeota | Methanomicrobia | Methanosarcinales | Methanosarcinaceae | Methanococcoides | burtonii |
| SP00021 | 2.00 | Euryarchaeota | Halobacteria | Haloferacales | Haloferacaceae | Haloquadratum | walsbyi |
| SP00022 | 3.00 | Euryarchaeota | Methanomicrobia | Methanosarcinales | Methanotrichaceae | Methanothrix | thermoacetophila |
| SP00023 | 1.00 | Crenarchaeota | Thermoprotei | Thermoproteales | Thermofilaceae | Thermofilum | pendens |
| SP00024 | 1.00 | Crenarchaeota | Thermoprotei | Desulfurococcales | Pyrodictiaceae | Hyperthermus | butylicus |
| SP00025 | 3.00 | Euryarchaeota | Methanomicrobia | Methanomicrobiales | Methanocorpusculaceae | Methanocorpusculum | labreanum |
| SP00026 | 1.00 | Crenarchaeota | Thermoprotei | Desulfurococcales | Desulfurococcaceae | Staphylothermus | marinus |
| SP00027 | 1.00 | Euryarchaeota | Methanomicrobia | Methanomicrobiales | Methanomicrobiaceae | Methanoculleus | marisnigri |
| SP00028 | 1.00 | Crenarchaeota | Thermoprotei | Thermoproteales | Thermoproteaceae | Pyrobaculum | arsenaticum |
| SP00029 | 3.00 | Euryarchaeota | Methanomicrobia | Methanocellales | Methanocellaceae | Methanocella | arvoryzae |
| SP00030 | 2.00 | Euryarchaeota | Methanobacteria | Methanobacteriales | Methanobacteriaceae | Methanobrevibacter | smithii |
| SP00031 | 4.00 | Euryarchaeota | Methanococci | Methanococcales | Methanococcaceae | Methanococcus | vannielii |
| SP00032 | 2.00 | Euryarchaeota | Methanococci | Methanococcales | Methanococcaceae | Methanococcus | aeolicus |
| SP00033 | 1.00 | Euryarchaeota | Methanomicrobia | Methanomicrobiales | Methanoregulaceae | Methanoregula | boonei |
| SP00034 | 1.00 | Crenarchaeota | Thermoprotei | Desulfurococcales | Desulfurococcaceae | Ignicoccus | hospitalis |
| SP00035 | 1.00 | Crenarchaeota | Thermoprotei | Sulfolobales | Sulfolobaceae | Acidianus | hospitalis |
| SP00036 | 1.00 | Euryarchaeota | Thermococci | Thermococcales | Thermococcaceae | Thermococcus | onnurineus |
| SP00037 | 1.00 | Crenarchaeota | Thermoprotei | Desulfurococcales | Desulfurococcaceae | Desulfurococcus | amylolyticus |
| SP00038 | 3.00 | Euryarchaeota | Methanomicrobia | Methanomicrobiales | Methanoregulaceae | Methanosphaerula | palustris |
| SP00039 | 3.00 | Euryarchaeota | Halobacteria | Haloferacales | Halorubraceae | Halorubrum | lacusprofundi |
| SP00040 | 1.00 | Crenarchaeota | Thermoprotei | Sulfolobales | Sulfolobaceae | Sulfolobus | islandicus |
| SP00041 | 1.00 | Euryarchaeota | Thermococci | Thermococcales | Thermococcaceae | Thermococcus | gammatolerans |
| SP00042 | 1.00 | Euryarchaeota | Thermococci | Thermococcales | Thermococcaceae | Thermococcus | sibiricus |
| SP00043 | 2.00 | Euryarchaeota | Methanococci | Methanococcales | Methanocaldococcaceae | Methanocaldococcus | fervens |
| SP00044 | 1.00 | Euryarchaeota | Halobacteria | Halobacteriales | Haloarculaceae | Halorhabdus | utahensis |
| SP00045 | 3.00 | Euryarchaeota | Halobacteria | Halobacteriales | Haloarculaceae | Halomicrobium | mukohataei |
| SP00046 | 2.00 | Euryarchaeota | Methanococci | Methanococcales | Methanocaldococcaceae | Methanocaldococcus | vulcanius |
| SP00047 | 2.00 | Euryarchaeota | Methanomicrobia | Methanocellales | Methanocellaceae | Methanocella | paludicola |
| SP00048 | 1.00 | Euryarchaeota | Archaeoglobi | Archaeoglobales | Archaeoglobaceae | Archaeoglobus | profundus |
| SP00049 | 3.00 | Euryarchaeota | Halobacteria | Natrialbales | Natrialbaceae | Haloterrigena | turkmenica |
| SP00050 | 2.00 | Euryarchaeota | Methanobacteria | Methanobacteriales | Methanobacteriaceae | Methanobrevibacter | ruminantium |
| SP00051 | 1.00 | Euryarchaeota | Archaeoglobi | Archaeoglobales | Archaeoglobaceae | Ferroglobus | placidus |
| SP00052 | 2.00 | Euryarchaeota | Methanococci | Methanococcales | Methanocaldococcaceae | Methanocaldococcus | sp. FS406-22 |
| SP00053 | 3.00 | Euryarchaeota | Halobacteria | Natrialbales | Natrialbaceae | Natrialba | magadii |
| SP00054 | 2.00 | Euryarchaeota | Halobacteria | Haloferacales | Haloferacaceae | Haloferax | volcanii |
| SP00055 | 3.00 | Euryarchaeota | Methanomicrobia | Methanosarcinales | Methanosarcinaceae | Methanohalophilus | mahii |
| SP00056 | 2.00 | Euryarchaeota | Methanococci | Methanococcales | Methanocaldococcaceae | Methanocaldococcus | infernus |
| SP00057 | 1.00 | Crenarchaeota | Thermoprotei | Desulfurococcales | Desulfurococcaceae | Staphylothermus | hellenicus |
| SP00058 | 2.00 | Euryarchaeota | Methanococci | Methanococcales | Methanococcaceae | Methanococcus | voltae |
| SP00059 | 2.00 | Euryarchaeota | Methanomicrobia | Methanosarcinales | Methanosarcinaceae | Methanohalobium | evestigatum |
| SP00060 | 1.00 | Euryarchaeota | Halobacteria | Halobacteriales | Halobacteriaceae | Halalkalicoccus | jeotgali |
| SP00061 | 1.00 | Crenarchaeota | Thermoprotei | Acidilobales | Acidilobaceae | Acidilobus | saccharovorans |
| SP00062 | 2.00 | Euryarchaeota | Methanobacteria | Methanobacteriales | Methanobacteriaceae | Methanothermobacter | marburgensis |
| SP00063 | 2.00 | Euryarchaeota | Methanomicrobia | Methanomicrobiales | Methanomicrobiaceae | Methanolacinia | petrolearia |
| SP00064 | 1.00 | Crenarchaeota | Thermoprotei | Thermoproteales | Thermoproteaceae | Vulcanisaeta | distributa |
| SP00065 | 2.00 | Euryarchaeota | Methanobacteria | Methanobacteriales | Methanothermaceae | Methanothermus | fervidus |
| SP00066 | 3.00 | Euryarchaeota | Halobacteria | Haloferacales | Haloferacaceae | Halogeometricum | borinquense |
| SP00067 | 1.00 | Euryarchaeota | Thermococci | Thermococcales | Thermococcaceae | Thermococcus | barophilus |
| SP00068 | 1.00 | Crenarchaeota | Thermoprotei | Desulfurococcales | Desulfurococcaceae | Desulfurococcus | mucosus |
| SP00069 | 2.00 | Euryarchaeota | Methanobacteria | Methanobacteriales | Methanobacteriaceae | Methanobacterium | lacus |
| SP00070 | 1.00 | Crenarchaeota | Thermoprotei | Thermoproteales | Thermoproteaceae | Thermoproteus | uzoniensis |
| SP00071 | 1.00 | Euryarchaeota | Archaeoglobi | Archaeoglobales | Archaeoglobaceae | Archaeoglobus | veneficus |
| SP00072 | 3.00 | Euryarchaeota | Methanomicrobia | Methanosarcinales | Methanotrichaceae | Methanothrix | soehngenii |
| SP00073 | 1.00 | Crenarchaeota | Thermoprotei | Sulfolobales | Sulfolobaceae | Metallosphaera | cuprina |
| SP00074 | 1.00 | Euryarchaeota | Thermococci | Thermococcales | Thermococcaceae | Pyrococcus | sp. NA2 |
| SP00075 | 2.00 | Euryarchaeota | Methanococci | Methanococcales | Methanocaldococcaceae | Methanotorris | igneus |
| SP00076 | 3.00 | Euryarchaeota | Methanobacteria | Methanobacteriales | Methanobacteriaceae | Methanobacterium | paludis |
| SP00077 | 2.00 | Euryarchaeota | Methanococci | Methanococcales | Methanococcaceae | Methanothermococcus | okinawensis |
| SP00078 | 3.00 | Euryarchaeota | Halobacteria | Natrialbales | Natrialbaceae | Halopiger | xanaduensis |
| SP00079 | 3.00 | Euryarchaeota | Methanomicrobia | Methanosarcinales | Methanosarcinaceae | Methanosalsum | zhilinae |
| SP00080 | 1.00 | Euryarchaeota | Thermococci | Thermococcales | Thermococcaceae | Pyrococcus | yayanosii |
| SP00081 | 1.00 | Euryarchaeota | Thermococci | Thermococcales | Thermococcaceae | Thermococcus | sp. 4557 |
| SP00082 | 1.00 | Crenarchaeota | Thermoprotei | Desulfurococcales | Pyrodictiaceae | Pyrolobus | fumarii |
| SP00083 | 3.00 | Euryarchaeota | Halobacteria | Halobacteriales | Haloarculaceae | Haloarcula | hispanica |
| SP00084 | 1.00 | Crenarchaeota | Thermoprotei | Thermoproteales | Thermoproteaceae | Thermoproteus | tenax |
| SP00085 | 1.00 | Euryarchaeota | Methanomicrobia | Methanosarcinales | Methanotrichaceae | Methanothrix | harundinacea |
| SP00086 | 1.00 | Crenarchaeota | Thermoprotei | Thermoproteales | Thermoproteaceae | Pyrobaculum | ferrireducens |
| SP00087 | 2.00 | Euryarchaeota | Methanomicrobia | Methanocellales | Methanocellaceae | Methanocella | conradii |
| SP00088 | 1.00 | Crenarchaeota | Thermoprotei | Fervidicoccales | Fervidicoccaceae | Fervidicoccus | fontis |
| SP00089 | 1.00 | Euryarchaeota | Thermococci | Thermococcales | Thermococcaceae | Pyrococcus | sp. ST04 |
| SP00090 | 1.00 | Crenarchaeota | Thermoprotei | Desulfurococcales | Desulfurococcaceae | Thermogladius | calderae |
| SP00091 | 1.00 | Euryarchaeota | Thermococci | Thermococcales | Thermococcaceae | Thermococcus | cleftensis |
| SP00092 | 3.00 | Euryarchaeota | Halobacteria | Natrialbales | Natrialbaceae | Natrinema | sp. J7-2 |
| SP00093 | 1.00 | Euryarchaeota | Methanomicrobia | Methanomicrobiales | Methanomicrobiaceae | Methanoculleus | bourgensis |
| SP00094 | 1.00 | Thaumarchaeota | Nitrososphaeria | Nitrososphaerales | Nitrososphaeraceae | Nitrososphaera | gargensis (c.) |
| SP00095 | 1.00 | Crenarchaeota | Thermoprotei | Acidilobales | Caldisphaeraceae | Caldisphaera | lagunensis |
| SP00096 | 3.00 | Euryarchaeota | Halobacteria | Natrialbales | Natrialbaceae | Natronobacterium | gregoryi |
| SP00097 | 1.00 | Euryarchaeota | Methanomicrobia | Methanomicrobiales | Methanoregulaceae | Methanoregula | formicica |
| SP00098 | 3.00 | Euryarchaeota | Halobacteria | Natrialbales | Natrialbaceae | Natrinema | pellirubrum |
| SP00099 | 2.00 | Euryarchaeota | Halobacteria | Natrialbales | Natrialbaceae | Halovivax | ruber |
| SP00100 | 2.00 | Euryarchaeota | Methanomicrobia | Methanosarcinales | Methanosarcinaceae | Methanomethylovorans | hollandica |
| SP00101 | 3.00 | Euryarchaeota | Halobacteria | Natrialbales | Natrialbaceae | Natronococcus | occultus |
| SP00102 | 1.00 | Euryarchaeota | Halobacteria | Halobacteriales | Haloarculaceae | Natronomonas | moolapensis |
| SP00103 | 1.00 | C.Thermoplasmatota | Thermoplasmata | Methanomassiliicoccales | C.Methanomethylophilaceae | C.Methanomethylophilus | alvus |
| SP00104 | 1.00 | Euryarchaeota | Archaeoglobi | Archaeoglobales | Archaeoglobaceae | Archaeoglobus | sulfaticallidus |
| SP00105 | 1.00 | C.Thermoplasmatota | Thermoplasmata | Methanomassiliicoccales | Methanomassiliicoccaceae | Methanomassiliicoccus | intestinalis (c.) |
| SP00106 | 3.00 | Euryarchaeota | Methanobacteria | Methanobacteriales | Methanobacteriaceae | Methanobrevibacter | sp. AbM4 |
| SP00107 | 1.00 | C.Thermoplasmatota | Thermoplasmata | Thermoplasmatales | Ferroplasmaceae | Ferroplasma | acidarmanus |
| SP00108 | 1.00 | Euryarchaeota | Halobacteria | Halobacteriales | Haloarculaceae | Halorhabdus | tiamatea |
| SP00109 | 1.00 | Euryarchaeota | Thermococci | Thermococcales | Thermococcaceae | Thermococcus | litoralis |
| SP00110 | 1.00 | Crenarchaeota | Thermoprotei | Desulfurococcales | Desulfurococcaceae | Aeropyrum | camini |
| SP00111 | 2.00 | Euryarchaeota | Thermococci | Thermococcales | Thermococcaceae | Palaeococcus | pacificus |
| SP00112 | 4.00 | Euryarchaeota | Methanobacteria | Methanobacteriales | Methanobacteriaceae | Methanobacterium | formicicum |
| SP00113 | 1.00 | Euryarchaeota | Thermococci | Thermococcales | Thermococcaceae | Thermococcus | paralvinellae |
| SP00114 | 2.00 | Euryarchaeota | Halobacteria | Natrialbales | Natrialbaceae | Halostagnicola | larsenii |
| SP00115 | 1.00 | Euryarchaeota | Halobacteria | Halobacteriales | Halobacteriaceae | Halobacterium | sp. DL1 |
| SP00116 | 1.00 | Thaumarchaeota | Nitrososphaeria | Nitrososphaerales | Nitrososphaeraceae | Nitrososphaera | evergladensis (c.) |
| SP00117 | 1.00 | Thaumarchaeota | Nitrososphaeria | Nitrososphaerales | Nitrososphaeraceae | Nitrososphaera | viennensis |
| SP00118 | 1.00 | Euryarchaeota | Thermococci | Thermococcales | Thermococcaceae | Thermococcus | eurythermalis |
| SP00119 | 2.00 | Euryarchaeota | Methanococci | Methanococcales | Methanocaldococcaceae | Methanocaldococcus | bathoardescens |
| SP00120 | 3.00 | Euryarchaeota | Methanomicrobia | Methanosarcinales | Methanosarcinaceae | Methanosarcina | thermophila |
| SP00121 | 3.00 | Euryarchaeota | Methanomicrobia | Methanosarcinales | Methanosarcinaceae | Methanosarcina | sp. WWM596 |
| SP00122 | 3.00 | Euryarchaeota | Methanomicrobia | Methanosarcinales | Methanosarcinaceae | Methanosarcina | sp. WH1 |
| SP00123 | 3.00 | Euryarchaeota | Methanomicrobia | Methanosarcinales | Methanosarcinaceae | Methanosarcina | sp. MTP4 |
| SP00124 | 3.00 | Euryarchaeota | Methanomicrobia | Methanosarcinales | Methanosarcinaceae | Methanosarcina | siciliae |
| SP00125 | 3.00 | Euryarchaeota | Methanomicrobia | Methanosarcinales | Methanosarcinaceae | Methanosarcina | lacustris |
| SP00126 | 3.00 | Euryarchaeota | Methanomicrobia | Methanosarcinales | Methanosarcinaceae | Methanosarcina | horonobensis |
| SP00127 | 3.00 | Euryarchaeota | Methanomicrobia | Methanosarcinales | Methanosarcinaceae | Methanococcoides | methylutens |
| SP00128 | 3.00 | Euryarchaeota | Methanomicrobia | Methanosarcinales | Methanosarcinaceae | Methanosarcina | vacuolata |
| SP00129 | 3.00 | Euryarchaeota | Methanomicrobia | Methanosarcinales | Methanosarcinaceae | Methanosarcina | sp. Kolksee |
| SP00130 | 1.00 | C.Thermoplasmatota | Thermoplasmata | Methanomassiliicoccales | Methanomassiliicoccaceae | C.Methanoplasma | termitum |
| SP00131 | 3.00 | Euryarchaeota | Halobacteria | Halobacteriales | Haloarculaceae | Haloarcula | sp. CBA1115 |
| SP00132 | 2.00 | Euryarchaeota | Methanobacteria | Methanobacteriales | Methanobacteriaceae | Methanobrevibacter | millerae |
| SP00133 | 2.00 | Euryarchaeota | Halobacteria | Halobacteriales | Haloarculaceae | Halomicrobium | sp. ZPS1 |
| SP00134 | 4.00 | Euryarchaeota | Methanobacteria | Methanobacteriales | Methanobacteriaceae | Methanosphaera | stadtmanae |
| SP00135 | 3.00 | Euryarchaeota | Methanobacteria | Methanobacteriales | Methanobacteriaceae | Methanobacterium | congolense |

The mean number of intragenomic 16S rRNA genes of these species were calculated in a previous investigation of our group (Regueira-Iglesias et al. 2021.b). C.= candidatus; ID= species identifier; Num= number; sp.= species.

Appendix Table 5. Primer pairs that produced the maximum number of identities/species ≥10 in the oral bacteria genomes.

| **ALC** | **Bacterial-specific primer pairs** | **ID** | **Number of Ids/sp.** |
| --- | --- | --- | --- |
| S | OP_F098_OP_R119 | SP00194 | 12 |
|  |  | SP00199 | 12 |
|  |  | SP00241 | 12 |
|  |  | SP00253 | 12 |
|  |  | SP00297 | 10 |
|  | OP_F066_KP_R040 | SP00184 | 14 |
|  |  | SP00177 | 13 |
|  |  | SP00251 | 13 |
|  |  | SP00315 | 13 |
|  |  | SP00319 | 13 |
|  |  | SP00148 | 12 |
|  |  | SP00194 | 12 |
|  |  | SP00199 | 12 |
|  |  | SP00297 | 12 |
|  |  | SP00160 | 11 |
|  |  | SP00163 | 11 |
|  |  | SP00181 | 11 |
|  |  | SP00237 | 11 |
|  |  | SP00248 | 11 |
|  |  | SP00252 | 11 |
|  |  | SP00258 | 11 |
|  |  | SP00279 | 11 |
|  |  | SP00280 | 11 |
|  |  | SP00302 | 11 |
|  |  | SP00143 | 11 |
|  |  | SP00154 | 11 |
|  |  | SP00151 | 10 |
|  |  | SP00179 | 10 |
|  |  | SP00186 | 10 |
|  | OP_F009_OP_R030 | SP00280 | 10 |
|  | KP_F061_KP_R074 | SP00163 | 10 |
|  |  | SP00280 | 10 |
| M | KP_F048_KP_R031 | SP00194 | 11 |
|  | KP_F051_KP_R041 | SP00199 | 13 |
|  |  | SP00253 | 12 |
|  |  | SP00194 | 11 |
|  |  | SP00241 | 11 |
|  | OP_F116_KP_R060 | SP00143 | 15 |
|  |  | SP00177 | 15 |
|  |  | SP00184 | 15 |
|  |  | SP00242 | 15 |
|  |  | SP00251 | 15 |
|  |  | SP00320 | 15 |
|  |  | SP00148 | 15 |
|  |  | SP00146 | 14 |
|  |  | SP00194 | 14 |
|  |  | SP00199 | 14 |
|  |  | SP00241 | 14 |
|  |  | SP00253 | 14 |
|  |  | SP00288 | 14 |
|  |  | SP00315 | 14 |
|  |  | SP00319 | 14 |
| L | KP_F048_KP_R060 | SP00194 | 10 |
|  | KP_F056_KP_R077 | SP00199 | 13 |
|  |  | SP00194 | 12 |
|  |  | SP00177 | 11 |
|  |  | SP00253 | 11 |
|  |  | SP00251 | 10 |
| **ALC** | **Bacterial and archaeal primer pairs** | **ID** | **Number of Ids/sp.** |
| S | KP_F020_KP_R032 | SP00194 | 12 |
|  |  | SP00199 | 12 |
|  |  | SP00241 | 12 |
|  |  | SP00253 | 12 |
|  |  | SP00297 | 10 |
|  | OP_F066_OP_R073 | SP00160 | 11 |
|  |  | SP00163 | 11 |
|  |  | SP00179 | 11 |
|  |  | SP00186 | 11 |
|  |  | SP00248 | 11 |
|  |  | SP00252 | 11 |
|  |  | SP00258 | 11 |
|  |  | SP00279 | 11 |
|  |  | SP00280 | 11 |
|  |  | SP00297 | 11 |
|  |  | SP00302 | 11 |
|  |  | SP00154 | 11 |
|  |  | SP00148 | 10 |
|  |  | SP00194 | 10 |
|  |  | SP00199 | 10 |
|  |  | SP00143 | 10 |
| M | OP_F114_KP_R031 | SP00194 | 11 |
|  | KP_F020_OP_R073 | SP00199 | 13 |
|  |  | SP00194 | 11 |
|  |  | SP00253 | 11 |
|  |  | SP00241 | 10 |
| L | OP_F114_OP_R121 | SP00194 | 10 |
|  | KP_F020_OP_R121 | SP00177 | 13 |
|  |  | SP00199 | 13 |
|  |  | SP00194 | 12 |
|  |  | SP00253 | 12 |
|  |  | SP00251 | 10 |
|  | OP_F066_OP_R121 | SP00143 | 14 |
|  |  | SP00199 | 12 |
|  |  | SP00177 | 12 |
|  |  | SP00194 | 11 |
|  |  | SP00241 | 11 |
|  |  | SP00251 | 11 |
|  |  | SP00146 | 10 |
|  |  | SP00288 | 10 |
|  |  | SP00315 | 10 |
|  |  | SP00319 | 10 |
| **ALC** | **Most used primer pairs** | **ID** | **Number of Ids/sp.** |
| S | KP_F078_OP_R010 ^B+A^ | SP00194 | 12 |
|  |  | SP00199 | 12 |
|  |  | SP00241 | 12 |
|  |  | SP00253 | 12 |
|  |  | SP00297 | 10 |
| M | KP_F047_KP_R035^B^ | SP00194 | 12 |
|  | OP_F009_OP_R029^B^ | SP00194 | 10 |

A= archaea; ALC= amplicon length category; B= bacteria; ID= species identifier; Ids= identities; L= long mean amplicon length category, >600 base pairs; M= medium mean amplicon length category, 301-600 base pairs; S= short mean amplicon length category, 100-300 base pairs; sp.= species.

Appendix Table 6. Primer pairs that produced maximum number of identities/species ≥10 in the oral archaea genomes.

| **ALC** | **Archaeal-specific primer pairs** | **ID** | **Number of Ids/sp.** |
| --- | --- | --- | --- |
| S | KP_F018_KP_R002 | SP00036 | 11 |
|  | OP_F066_KP_R013 | SP00036 | 13 |
|  |  | SP00041 | 13 |
|  |  | SP00042 | 13 |
|  |  | SP00067 | 13 |
|  |  | SP00074 | 13 |
|  |  | SP00080 | 13 |
|  |  | SP00081 | 13 |
|  |  | SP00089 | 13 |
|  |  | SP00091 | 13 |
|  |  | SP00109 | 13 |
|  |  | SP00111 | 13 |
|  |  | SP00113 | 13 |
|  |  | SP00118 | 13 |
|  |  | SP00006 | 13 |
|  |  | SP00013 | 11 |
|  |  | SP00121 | 11 |
|  |  | SP00122 | 11 |
|  |  | SP00125 | 11 |
|  |  | SP00017 | 10 |
|  |  | SP00120 | 10 |
|  |  | SP00124 | 10 |
|  |  | SP00126 | 10 |
|  |  | SP00128 | 10 |
|  |  | SP00129 | 10 |
|  |  | SP00012 | 10 |
| M | KP_F018_KP_R032 | SP00036 | 13 |
|  |  | SP00017 | 10 |
|  |  | SP00124 | 10 |
|  |  | SP00126 | 10 |
|  |  | SP00128 | 10 |
|  |  | SP00129 | 10 |
|  |  | SP00118 | 10 |
|  |  | SP00012 | 10 |
|  | KP_F018_OP_R073 | SP00036 | 13 |
|  |  | SP00041 | 13 |
|  |  | SP00091 | 13 |
|  |  | SP00067 | 12 |
|  |  | SP00113 | 12 |
|  |  | SP00013 | 10 |
|  |  | SP00017 | 10 |
|  |  | SP00120 | 10 |
|  |  | SP00121 | 10 |
|  |  | SP00122 | 10 |
|  |  | SP00124 | 10 |
|  |  | SP00125 | 10 |
|  |  | SP00126 | 10 |
|  |  | SP00128 | 10 |
|  |  | SP00129 | 10 |
|  |  | SP00118 | 10 |
|  |  | SP00012 | 10 |
|  | KP_F020_KP_R013 | SP00036 | 12 |
|  |  | SP00041 | 12 |
|  |  | SP00042 | 12 |
|  |  | SP00067 | 12 |
|  |  | SP00074 | 12 |
|  |  | SP00080 | 12 |
|  |  | SP00081 | 12 |
|  |  | SP00089 | 12 |
|  |  | SP00091 | 12 |
|  |  | SP00109 | 12 |
|  |  | SP00111 | 12 |
|  |  | SP00113 | 12 |
|  |  | SP00006 | 12 |
|  |  | SP00013 | 11 |
|  |  | SP00121 | 11 |
|  |  | SP00122 | 11 |
|  |  | SP00125 | 11 |
|  |  | SP00128 | 11 |
|  |  | SP00129 | 11 |
|  |  | SP00017 | 10 |
|  |  | SP00120 | 10 |
|  |  | SP00124 | 10 |
|  |  | SP00126 | 10 |
|  |  | SP00012 | 10 |
|  | KP_F022_KP_R063 | SP00013 | 10 |
|  |  | SP00121 | 10 |
|  |  | SP00122 | 10 |
|  |  | SP00124 | 10 |
|  |  | SP00126 | 10 |
|  |  | SP00012 | 10 |
| L | OP_F114_KP_R013 | SP00036 | 13 |
|  |  | SP00041 | 13 |
|  |  | SP00067 | 13 |
|  |  | SP00091 | 13 |
|  |  | SP00109 | 13 |
|  |  | SP00113 | 13 |
|  |  | SP00118 | 13 |
|  |  | SP00042 | 12 |
|  |  | SP00080 | 12 |
|  |  | SP00006 | 12 |
|  |  | SP00074 | 11 |
|  |  | SP00081 | 11 |
|  |  | SP00013 | 10 |
|  |  | SP00017 | 10 |
|  |  | SP00089 | 10 |
|  |  | SP00120 | 10 |
|  |  | SP00121 | 10 |
|  |  | SP00122 | 10 |
|  |  | SP00124 | 10 |
|  |  | SP00125 | 10 |
|  |  | SP00126 | 10 |
|  |  | SP00128 | 10 |
|  |  | SP00129 | 10 |
|  |  | SP00012 | 10 |
|  | KP_F018_KP_R063 | SP00113 | 11 |
|  |  | SP00013 | 10 |
|  |  | SP00017 | 10 |
|  |  | SP00036 | 10 |
|  |  | SP00120 | 10 |
|  |  | SP00121 | 10 |
|  |  | SP00122 | 10 |
|  |  | SP00124 | 10 |
|  |  | SP00125 | 10 |
|  |  | SP00126 | 10 |
|  |  | SP00128 | 10 |
|  |  | SP00129 | 10 |
|  |  | SP00012 | 10 |
|  | OP_F066_OP_R016 | SP00036 | 13 |
|  |  | SP00041 | 13 |
|  |  | SP00067 | 13 |
|  |  | SP00080 | 13 |
|  |  | SP00081 | 13 |
|  |  | SP00089 | 13 |
|  |  | SP00091 | 13 |
|  |  | SP00109 | 13 |
|  |  | SP00111 | 13 |
|  |  | SP00113 | 13 |
|  |  | SP00118 | 13 |
|  |  | SP00006 | 13 |
|  |  | SP00042 | 12 |
|  |  | SP00074 | 12 |
|  |  | SP00013 | 10 |
|  |  | SP00017 | 10 |
|  |  | SP00120 | 10 |
|  |  | SP00121 | 10 |
|  |  | SP00122 | 10 |
|  |  | SP00124 | 10 |
|  |  | SP00125 | 10 |
|  |  | SP00126 | 10 |
|  |  | SP00128 | 10 |
|  |  | SP00129 | 10 |
|  |  | SP00012 | 10 |
| **ALC** | **Bacterial and archaeal primer pairs** | **ID** | **Number of Ids/sp.** |
| S | OP_F114_KP_R002 | SP00036 | 11 |
|  | KP_F020_KP_R032 | SP00036 | 12 |
|  |  | SP00041 | 12 |
|  |  | SP00067 | 12 |
|  |  | SP00091 | 12 |
|  |  | SP00113 | 12 |
|  |  | SP00013 | 11 |
|  |  | SP00017 | 11 |
|  |  | SP00124 | 11 |
|  |  | SP00126 | 11 |
|  |  | SP00128 | 11 |
|  |  | SP00129 | 11 |
|  |  | SP00012 | 11 |
|  |  | SP00120 | 10 |
|  |  | SP00121 | 10 |
|  |  | SP00122 | 10 |
|  |  | SP00125 | 10 |
|  | OP_F066_OP_R073 | SP00118 | 14 |
|  |  | SP00036 | 13 |
|  |  | SP00041 | 13 |
|  |  | SP00042 | 13 |
|  |  | SP00074 | 13 |
|  |  | SP00080 | 13 |
|  |  | SP00081 | 13 |
|  |  | SP00089 | 13 |
|  |  | SP00091 | 13 |
|  |  | SP00006 | 13 |
|  |  | SP00067 | 12 |
|  |  | SP00109 | 12 |
|  |  | SP00113 | 12 |
|  |  | SP00013 | 10 |
|  |  | SP00017 | 10 |
|  |  | SP00026 | 10 |
|  |  | SP00057 | 10 |
|  |  | SP00111 | 10 |
|  |  | SP00120 | 10 |
|  |  | SP00121 | 10 |
|  |  | SP00122 | 10 |
|  |  | SP00124 | 10 |
|  |  | SP00125 | 10 |
|  |  | SP00126 | 10 |
|  |  | SP00128 | 10 |
|  |  | SP00129 | 10 |
|  |  | SP00012 | 10 |
| M | OP_F114_KP_R031 | SP00036 | 13 |
|  |  | SP00017 | 10 |
|  |  | SP00124 | 10 |
|  |  | SP00126 | 10 |
|  |  | SP00128 | 10 |
|  |  | SP00129 | 10 |
|  |  | SP00118 | 10 |
|  |  | SP00012 | 10 |
|  | OP_F114_OP_R073 | SP00036 | 13 |
|  |  | SP00041 | 13 |
|  |  | SP00091 | 13 |
|  |  | SP00067 | 12 |
|  |  | SP00113 | 12 |
|  |  | SP00013 | 10 |
|  |  | SP00017 | 10 |
|  |  | SP00120 | 10 |
|  |  | SP00121 | 10 |
|  |  | SP00122 | 10 |
|  |  | SP00124 | 10 |
|  |  | SP00125 | 10 |
|  |  | SP00126 | 10 |
|  |  | SP00128 | 10 |
|  |  | SP00129 | 10 |
|  |  | SP00118 | 10 |
|  |  | SP00012 | 10 |
|  | KP_F020_OP_R073 | SP00036 | 12 |
|  |  | SP00041 | 12 |
|  |  | SP00067 | 12 |
|  |  | SP00091 | 12 |
|  |  | SP00109 | 12 |
|  |  | SP00113 | 12 |
|  |  | SP00128 | 11 |
|  |  | SP00129 | 11 |
|  |  | SP00012 | 11 |
|  |  | SP00013 | 10 |
|  |  | SP00017 | 10 |
|  |  | SP00120 | 10 |
|  |  | SP00121 | 10 |
|  |  | SP00122 | 10 |
|  |  | SP00124 | 10 |
|  |  | SP00125 | 10 |
|  |  | SP00126 | 10 |
| L | OP_F114_OP_R121 | SP00036 | 13 |
|  |  | SP00041 | 13 |
|  |  | SP00067 | 13 |
|  |  | SP00091 | 13 |
|  |  | SP00109 | 13 |
|  |  | SP00113 | 13 |
|  |  | SP00118 | 13 |
|  |  | SP00013 | 10 |
|  |  | SP00017 | 10 |
|  |  | SP00074 | 10 |
|  |  | SP00080 | 10 |
|  |  | SP00089 | 10 |
|  |  | SP00120 | 10 |
|  |  | SP00121 | 10 |
|  |  | SP00122 | 10 |
|  |  | SP00124 | 10 |
|  |  | SP00125 | 10 |
|  |  | SP00126 | 10 |
|  |  | SP00128 | 10 |
|  |  | SP00129 | 10 |
|  |  | SP00006 | 10 |
|  |  | SP00012 | 10 |
|  | KP_F020_OP_R121 | SP00036 | 12 |
|  |  | SP00041 | 12 |
|  |  | SP00067 | 12 |
|  |  | SP00080 | 12 |
|  |  | SP00091 | 12 |
|  |  | SP00109 | 12 |
|  |  | SP00113 | 12 |
|  |  | SP00089 | 11 |
|  |  | SP00111 | 11 |
|  |  | SP00013 | 10 |
|  |  | SP00017 | 10 |
|  |  | SP00042 | 10 |
|  |  | SP00074 | 10 |
|  |  | SP00120 | 10 |
|  |  | SP00121 | 10 |
|  |  | SP00122 | 10 |
|  |  | SP00124 | 10 |
|  |  | SP00125 | 10 |
|  |  | SP00126 | 10 |
|  |  | SP00128 | 10 |
|  |  | SP00129 | 10 |
|  |  | SP00006 | 10 |
|  |  | SP00012 | 10 |
|  | OP_F066_OP_R121 | SP00036 | 13 |
|  |  | SP00041 | 13 |
|  |  | SP00067 | 13 |
|  |  | SP00080 | 13 |
|  |  | SP00081 | 13 |
|  |  | SP00089 | 13 |
|  |  | SP00091 | 13 |
|  |  | SP00109 | 13 |
|  |  | SP00111 | 13 |
|  |  | SP00113 | 13 |
|  |  | SP00118 | 13 |
|  |  | SP00006 | 13 |
|  |  | SP00042 | 12 |
|  |  | SP00074 | 12 |
|  |  | SP00013 | 10 |
|  |  | SP00017 | 10 |
|  |  | SP00120 | 10 |
|  |  | SP00121 | 10 |
|  |  | SP00122 | 10 |
|  |  | SP00124 | 10 |
|  |  | SP00125 | 10 |
|  |  | SP00126 | 10 |
|  |  | SP00128 | 10 |
|  |  | SP00129 | 10 |
|  |  | SP00012 | 10 |
| **ALC** | **Most used primer pairs** | **ID** | **Number of Ids/sp.** |
| S | KP_F078-OP_R010^B+A^ | SP00013 | 11 |
|  |  | SP00017 | 11 |
|  |  | SP00124 | 11 |
|  |  | SP00126 | 11 |
|  |  | SP00128 | 11 |
|  |  | SP00129 | 11 |
|  |  | SP00012 | 11 |
|  |  | SP00120 | 10 |
|  |  | SP00121 | 10 |
|  |  | SP00122 | 10 |
|  |  | SP00125 | 10 |
| L | KP_F014_KP_R011^A^ | SP00036 | 13 |
|  |  | SP00041 | 13 |
|  |  | SP00067 | 13 |
|  |  | SP00091 | 13 |
|  |  | SP00109 | 13 |
|  |  | SP00113 | 13 |
|  |  | SP00118 | 13 |
|  |  | SP00042 | 12 |
|  |  | SP00074 | 11 |
|  |  | SP00080 | 11 |
|  |  | SP00006 | 11 |
|  |  | SP00089 | 10 |

A= archaea; ALC= amplicon length category; B= bacteria; ID= species identifier; Ids= identities; L= long mean amplicon length category, >600 base pairs; M= medium mean amplicon length category, 301-600 base pairs; S= short mean amplicon length category, 100-300 base pairs; sp.= species; WUOL= widely used in the oral literature.

Appendix Table 7. Number of species with amplicon similarity values ≥97%, the number of identities, and the SC and SC-NASI≥97% values obtained by each primer pair analyzed against both domains.

| **ALC** | **Bacterial and archaeal primer pairs** | **No. sp.**  **ASI≥ 97%** | **Total**  **Ids** | **Mean Ids/sp.**  **(±SD)** | **Max. Ids/sp.** | **SC**  **No. sp. (%)** | **SC-NASI97% No. sp. (%)** |
| --- | --- | --- | --- | --- | --- | --- | --- |
| S | OP_F114-KP_R002 | 144 | 244 | 3.39 (2.78) | 11 | 306 (95.33) | 162 (50.47) |
|  | KP_F020-KP_R032 | 186 | 392 | 4.22 (3.50) | 12 | 310 (96.57) | 124 (38.63) |
|  | OP_F066-OP_R073 | 219 | 525 | 4.80 (3.84) | 14 | 308 (95.95) | 89 (27.73) |
| M | OP_F114-KP_R031 | 147 | 269 | 3.66 (3.30) | 13 | 314 (97.82) | 167 (52.02) |
|  | OP_F114-OP_R073 | 153 | 286 | 3.74 (3.63) | 13 | 304 (94.70) | 151 (47.04) |
|  | KP_F020-OP_R073 | 175 | 361 | 4.13 (3.50) | 13 | 301 (93.77) | 126 (39.25) |
| L | OP_F114-OP_R121 | 154 | 312 | 4.05 (3.91) | 13 | 309 (96.26) | 155 (48.29) |
|  | KP_F020-OP_R121 | 163 | 341 | 4.18 (3.93) | 13 | 306 (95.33) | 143 (44.55) |
|  | OP_F066-OP_R121 | 169 | 366 | 4.33 (4.25) | 14 | 315 (98.13) | 146 (45.48) |
| **ALC** | **Most used primer pairs** | **No. sp. ASI**  **≥ 97%** | **Total Ids** | **Mean Ids/sp.**  **(±SD)** | **Max. Ids/sp.** | **SC**  **No. sp. (%)** | **SC-NASI97%**  **No. sp. (%)** |
| S | KP_F078-OP_R010^B+A^ | 155 | 314 | 4.05 (3.37) | 12 | 262 (81.62) | 107 (33.33) |

A= archaea; ALC= amplicon length category; ASI≥97%= species with amplicon similarity/identity values ≥97%; B= bacteria; Ids= identities; L= long mean amplicon length category, >600 base pairs; M= medium mean amplicon length category, 301-600 base pairs; Max.= maximum; S= short mean amplicon length category, 100-300 base pairs; SC= species coverage; SC-NASI97%= species coverage without amplicon similarity/identity ≥97%; SD= standard deviation; sp.= species.

Appendix Table 8. Taxonomy of all different bacterial taxa with a maximum number of identities/species ≥10.

| **ID** | **Phylum** | **Class** | **Order** | **Family** | **Genus** | **Species** |
| --- | --- | --- | --- | --- | --- | --- |
| SP00179 | Firmicutes | Bacilli | Bacillales | Bacillaceae | Bacillus | subtilis |
| SP00237 | Firmicutes | Bacilli | Lactobacillales | Lactobacillaceae | Lentilactobacillus | buchneri |
| SP00181 | Firmicutes | Bacilli | Lactobacillales | Lactobacillaceae | Levilactobacillus | brevis |
| SP00160 | Firmicutes | Bacilli | Bacillales | Listeriaceae | Listeria | monocytogenes |
| SP00163 | Firmicutes | Bacilli | Bacillales | Staphylococcaceae | Staphylococcus | aureus |
| SP00258 | Firmicutes | Bacilli | Bacillales | Staphylococcaceae | Staphylococcus | capitis |
| SP00297 | Firmicutes | Bacilli | Bacillales | Staphylococcaceae | Staphylococcus | cohnii |
| SP00154 | Firmicutes | Bacilli | Bacillales | Staphylococcaceae | Staphylococcus | epidermidis |
| SP00279 | Firmicutes | Bacilli | Bacillales | Staphylococcaceae | Staphylococcus | haemolyticus |
| SP00280 | Firmicutes | Bacilli | Bacillales | Staphylococcaceae | Staphylococcus | lugdunensis |
| SP00252 | Firmicutes | Bacilli | Bacillales | Staphylococcaceae | Staphylococcus | pasteuri |
| SP00302 | Firmicutes | Bacilli | Bacillales | Staphylococcaceae | Staphylococcus | pettenkoferi |
| SP00186 | Firmicutes | Bacilli | Bacillales | Staphylococcaceae | Staphylococcus | schleiferi |
| SP00248 | Firmicutes | Bacilli | Bacillales | Staphylococcaceae | Staphylococcus | warneri |
| SP00148 | Firmicutes | Bacilli | Lactobacillales | Streptococcaceae | Streptococcus | agalactiae |
| SP00184 | Firmicutes | Bacilli | Lactobacillales | Streptococcaceae | Streptococcus | anginosus |
| SP00251 | Firmicutes | Bacilli | Lactobacillales | Streptococcaceae | Streptococcus | constellatus |
| SP00253 | Firmicutes | Bacilli | Lactobacillales | Streptococcaceae | Streptococcus | cristatus |
| SP00199 | Firmicutes | Bacilli | Lactobacillales | Streptococcaceae | Streptococcus | gordonii |
| SP00177 | Firmicutes | Bacilli | Lactobacillales | Streptococcaceae | Streptococcus | intermedius |
| SP00315 | Firmicutes | Bacilli | Lactobacillales | Streptococcaceae | Streptococcus | mitis |
| SP00151 | Firmicutes | Bacilli | Lactobacillales | Streptococcaceae | Streptococcus | mutans |
| SP00319 | Firmicutes | Bacilli | Lactobacillales | Streptococcaceae | Streptococcus | oralis |
| SP00241 | Firmicutes | Bacilli | Lactobacillales | Streptococcaceae | Streptococcus | parasanguinis |
| SP00146 | Firmicutes | Bacilli | Lactobacillales | Streptococcaceae | Streptococcus | pneumoniae |
| SP00143 | Firmicutes | Bacilli | Lactobacillales | Streptococcaceae | Streptococcus | pyogenes |
| SP00242 | Firmicutes | Bacilli | Lactobacillales | Streptococcaceae | Streptococcus | salivarius |
| SP00194 | Firmicutes | Bacilli | Lactobacillales | Streptococcaceae | Streptococcus | sanguinis |
| SP00288 | Firmicutes | Bacilli | Lactobacillales | Streptococcaceae | Streptococcus | sp. oral taxon 431 |
| SP00320 | Firmicutes | Bacilli | Lactobacillales | Streptococcaceae | Streptococcus | thermophilus |

ID= species identifier.

Appendix Table 9. Taxonomy of all different archaeal taxa with a maximum number of identities/species ≥10.

| **ID** | **Phylum** | **Class** | **Order** | **Family** | **Genus** | **Species** |
| --- | --- | --- | --- | --- | --- | --- |
| SP00012 | Euryarchaeota | Methanomicrobia | Methanosarcinales | Methanosarcinaceae | Methanosarcina | acetivorans |
| SP00017 | Euryarchaeota | Methanomicrobia | Methanosarcinales | Methanosarcinaceae | Methanosarcina | barkeri |
| SP00126 | Euryarchaeota | Methanomicrobia | Methanosarcinales | Methanosarcinaceae | Methanosarcina | horonobensis |
| SP00125 | Euryarchaeota | Methanomicrobia | Methanosarcinales | Methanosarcinaceae | Methanosarcina | lacustris |
| SP00013 | Euryarchaeota | Methanomicrobia | Methanosarcinales | Methanosarcinaceae | Methanosarcina | mazei |
| SP00124 | Euryarchaeota | Methanomicrobia | Methanosarcinales | Methanosarcinaceae | Methanosarcina | siciliae |
| SP00129 | Euryarchaeota | Methanomicrobia | Methanosarcinales | Methanosarcinaceae | Methanosarcina | sp. Kolksee |
| SP00122 | Euryarchaeota | Methanomicrobia | Methanosarcinales | Methanosarcinaceae | Methanosarcina | sp. WH1 |
| SP00121 | Euryarchaeota | Methanomicrobia | Methanosarcinales | Methanosarcinaceae | Methanosarcina | sp. WWM596 |
| SP00120 | Euryarchaeota | Methanomicrobia | Methanosarcinales | Methanosarcinaceae | Methanosarcina | thermophila |
| SP00128 | Euryarchaeota | Methanomicrobia | Methanosarcinales | Methanosarcinaceae | Methanosarcina | vacuolata |
| SP00111 | Euryarchaeota | Thermococci | Thermococcales | Thermococcaceae | Palaeococcus | pacificus |
| SP00006 | Euryarchaeota | Thermococci | Thermococcales | Thermococcaceae | Pyrococcus | horikoshii |
| SP00074 | Euryarchaeota | Thermococci | Thermococcales | Thermococcaceae | Pyrococcus | sp. NA2 |
| SP00089 | Euryarchaeota | Thermococci | Thermococcales | Thermococcaceae | Pyrococcus | sp. ST04 |
| SP00080 | Euryarchaeota | Thermococci | Thermococcales | Thermococcaceae | Pyrococcus | yayanosii |
| SP00057 | Crenarchaeota | Thermoprotei | Desulfurococcales | Desulfurococcaceae | Staphylothermus | hellenicus |
| SP00026 | Crenarchaeota | Thermoprotei | Desulfurococcales | Desulfurococcaceae | Staphylothermus | marinus |
| SP00067 | Euryarchaeota | Thermococci | Thermococcales | Thermococcaceae | Thermococcus | barophilus |
| SP00091 | Euryarchaeota | Thermococci | Thermococcales | Thermococcaceae | Thermococcus | cleftensis |
| SP00118 | Euryarchaeota | Thermococci | Thermococcales | Thermococcaceae | Thermococcus | eurythermalis |
| SP00041 | Euryarchaeota | Thermococci | Thermococcales | Thermococcaceae | Thermococcus | gammatolerans |
| SP00109 | Euryarchaeota | Thermococci | Thermococcales | Thermococcaceae | Thermococcus | litoralis |
| SP00036 | Euryarchaeota | Thermococci | Thermococcales | Thermococcaceae | Thermococcus | onnurineus |
| SP00113 | Euryarchaeota | Thermococci | Thermococcales | Thermococcaceae | Thermococcus | paralvinellae |
| SP00042 | Euryarchaeota | Thermococci | Thermococcales | Thermococcaceae | Thermococcus | sibiricus |
| SP00081 | Euryarchaeota | Thermococci | Thermococcales | Thermococcaceae | Thermococcus | sp. 4557 |

ID= species identifier.

Appendix Table 10. Oral bacteria species with amplicon similarity values ≥97% with at least one different taxon.

| **ID** | **Phylum** | **Class** | **Order** | **Family** | **Genus** | **Species** |
| --- | --- | --- | --- | --- | --- | --- |
| SP00136 | Proteobacteria | Gammaproteobacteria | Enterobacterales | Enterobacteriaceae | Escherichia | coli |
| SP00138 | Actinobacteria | Actinomycetia | Corynebacteriales | Mycobacteriaceae | Mycobacterium | tuberculosis |
| SP00141 | Proteobacteria | Betaproteobacteria | Neisseriales | Neisseriaceae | Neisseria | meningitidis |
| SP00142 | Proteobacteria | Gammaproteobacteria | Pseudomonadales | Pseudomonadaceae | Pseudomonas | aeruginosa |
| SP00143 | Firmicutes | Bacilli | Lactobacillales | Streptococcaceae | Streptococcus | pyogenes |
| SP00144 | Proteobacteria | Betaproteobacteria | Neisseriales | Neisseriaceae | Neisseria | gonorrhoeae |
| SP00146 | Firmicutes | Bacilli | Lactobacillales | Streptococcaceae | Streptococcus | pneumoniae |
| SP00147 | Proteobacteria | Alphaproteobacteria | Hyphomicrobiales | Rhizobiaceae | Agrobacterium | fabrum |
| SP00148 | Firmicutes | Bacilli | Lactobacillales | Streptococcaceae | Streptococcus | agalactiae |
| SP00149 | Fusobacteria | Fusobacteriia | Fusobacteriales | Fusobacteriaceae | Fusobacterium | nucleatum |
| SP00150 | Proteobacteria | Gammaproteobacteria | Enterobacterales | Yersiniaceae | Yersinia | pestis |
| SP00151 | Firmicutes | Bacilli | Lactobacillales | Streptococcaceae | Streptococcus | mutans |
| SP00152 | Actinobacteria | Actinomycetia | Bifidobacteriales | Bifidobacteriaceae | Bifidobacterium | longum |
| SP00154 | Firmicutes | Bacilli | Bacillales | Staphylococcaceae | Staphylococcus | epidermidis |
| SP00155 | Firmicutes | Bacilli | Lactobacillales | Enterococcaceae | Enterococcus | faecalis |
| SP00156 | Firmicutes | Bacilli | Bacillales | Bacillaceae | Bacillus | anthracis |
| SP00157 | Proteobacteria | Gammaproteobacteria | Pasteurellales | Pasteurellaceae | Haemophilus | ducreyi |
| SP00158 | Firmicutes | Bacilli | Lactobacillales | Lactobacillaceae | Lactobacillus | johnsonii |
| SP00159 | Spirochaetes | Spirochaetia | Spirochaetales | Spirochaetaceae | Treponema | denticola |
| SP00160 | Firmicutes | Bacilli | Bacillales | Listeriaceae | Listeria | monocytogenes |
| SP00161 | Actinobacteria | Actinomycetia | Propionibacteriales | Propionibacteriaceae | Cutibacterium | acnes |
| SP00162 | Firmicutes | Bacilli | Lactobacillales | Lactobacillaceae | Ligilactobacillus | salivarius |
| SP00163 | Firmicutes | Bacilli | Bacillales | Staphylococcaceae | Staphylococcus | aureus |
| SP00164 | Actinobacteria | Actinomycetia | Corynebacteriales | Mycobacteriaceae | Mycobacterium | leprae |
| SP00165 | Firmicutes | Bacilli | Lactobacillales | Lactobacillaceae | Lactiplantibacillus | plantarum |
| SP00166 | Proteobacteria | Gammaproteobacteria | Pseudomonadales | Pseudomonadaceae | Pseudomonas | fluorescens |
| SP00168 | Actinobacteria | Actinomycetia | Corynebacteriales | Corynebacteriaceae | Corynebacterium | urealyticum |
| SP00169 | Proteobacteria | Gammaproteobacteria | Enterobacterales | Morganellaceae | Proteus | mirabilis |
| SP00170 | Proteobacteria | Alphaproteobacteria | Hyphomicrobiales | Phyllobacteriaceae | Mesorhizobium | japonicum |
| SP00172 | Proteobacteria | Gammaproteobacteria | Enterobacterales | Enterobacteriaceae | Klebsiella | pneumoniae |
| SP00173 | Firmicutes | Bacilli | Lactobacillales | Lactobacillaceae | Limosilactobacillus | reuteri |
| SP00174 | Firmicutes | Bacilli | Lactobacillales | Lactobacillaceae | Limosilactobacillus | fermentum |
| SP00177 | Firmicutes | Bacilli | Lactobacillales | Streptococcaceae | Streptococcus | intermedius |
| SP00178 | Actinobacteria | Actinomycetia | Micrococcales | Micrococcaceae | Rothia | mucilaginosa |
| SP00179 | Firmicutes | Bacilli | Bacillales | Bacillaceae | Bacillus | subtilis |
| SP00181 | Firmicutes | Bacilli | Lactobacillales | Lactobacillaceae | Levilactobacillus | brevis |
| SP00182 | Tenericutes | Mollicutes | Mycoplasmatales | Mycoplasmataceae | Mycoplasma | pneumoniae |
| SP00184 | Firmicutes | Bacilli | Lactobacillales | Streptococcaceae | Streptococcus | anginosus |
| SP00185 | Proteobacteria | Gammaproteobacteria | Pasteurellales | Pasteurellaceae | Aggregatibacter | actinomycetemcomitans |
| SP00186 | Firmicutes | Bacilli | Bacillales | Staphylococcaceae | Staphylococcus | schleiferi |
| SP00187 | Actinobacteria | Actinomycetia | Corynebacteriales | Corynebacteriaceae | Corynebacterium | diphtheriae |
| SP00188 | Proteobacteria | Betaproteobacteria | Burkholderiales | Alcaligenaceae | Bordetella | pertussis |
| SP00189 | Firmicutes | Bacilli | Lactobacillales | Lactobacillaceae | Lactobacillus | acidophilus |
| SP00190 | Proteobacteria | Gammaproteobacteria | Pasteurellales | Pasteurellaceae | Haemophilus | influenzae |
| SP00191 | Proteobacteria | Gammaproteobacteria | Pseudomonadales | Pseudomonadaceae | Pseudomonas | protegens |
| SP00192 | Actinobacteria | Actinomycetia | Bifidobacteriales | Bifidobacteriaceae | Bifidobacterium | breve |
| SP00193 | Proteobacteria | Gammaproteobacteria | Pseudomonadales | Pseudomonadaceae | Pseudomonas | stutzeri |
| SP00194 | Firmicutes | Bacilli | Lactobacillales | Streptococcaceae | Streptococcus | sanguinis |
| SP00195 | Firmicutes | Bacilli | Lactobacillales | Lactobacillaceae | Lactobacillus | gasseri |
| SP00196 | Firmicutes | Bacilli | Lactobacillales | Lactobacillaceae | Lacticaseibacillus | paracasei |
| SP00197 | Proteobacteria | Gammaproteobacteria | Pseudomonadales | Moraxellaceae | Acinetobacter | baumannii |
| SP00198 | Proteobacteria | Alphaproteobacteria | Hyphomicrobiales | Rhizobiaceae | Agrobacterium | radiobacter |
| SP00199 | Firmicutes | Bacilli | Lactobacillales | Streptococcaceae | Streptococcus | gordonii |
| SP00200 | Proteobacteria | Alphaproteobacteria | Hyphomicrobiales | Brucellaceae | Brucella | anthropi |
| SP00201 | Proteobacteria | Epsilonproteobacteria | Campylobacterales | Campylobacteraceae | Campylobacter | curvus |
| SP00202 | Proteobacteria | Gammaproteobacteria | Enterobacterales | Enterobacteriaceae | Cronobacter | sakazakii |
| SP00203 | Proteobacteria | Epsilonproteobacteria | Campylobacterales | Campylobacteraceae | Campylobacter | concisus |
| SP00204 | Proteobacteria | Betaproteobacteria | Burkholderiales | Comamonadaceae | Delftia | acidovorans |
| SP00205 | Proteobacteria | Gammaproteobacteria | Enterobacterales | Enterobacteriaceae | Klebsiella | variicola |
| SP00206 | Proteobacteria | Betaproteobacteria | Burkholderiales | Burkholderiaceae | Ralstonia | pickettii |
| SP00208 | Actinobacteria | Actinomycetia | Bifidobacteriales | Bifidobacteriaceae | Bifidobacterium | animalis |
| SP00209 | Proteobacteria | Betaproteobacteria | Burkholderiales | Comamonadaceae | Comamonas | thiooxydans |
| SP00210 | Proteobacteria | Alphaproteobacteria | Rhodobacterales | Rhodobacteraceae | Rhodobacter | capsulatus |
| SP00211 | Proteobacteria | Betaproteobacteria | Burkholderiales | Comamonadaceae | Acidovorax | ebreus |
| SP00212 | Proteobacteria | Gammaproteobacteria | Pasteurellales | Pasteurellaceae | Aggregatibacter | aphrophilus |
| SP00213 | Actinobacteria | Actinomycetia | Corynebacteriales | Corynebacteriaceae | Corynebacterium | kroppenstedtii |
| SP00214 | Actinobacteria | Actinomycetia | Micrococcales | Micrococcaceae | Micrococcus | luteus |
| SP00215 | Bacteroidetes | Flavobacteriia | Flavobacteriales | Flavobacteriaceae | Capnocytophaga | ochracea |
| SP00216 | Proteobacteria | Betaproteobacteria | Burkholderiales | Comamonadaceae | Variovorax | paradoxus |
| SP00218 | Fusobacteria | Fusobacteriia | Fusobacteriales | Leptotrichiaceae | Leptotrichia | buccalis |
| SP00219 | Actinobacteria | Actinomycetia | Micrococcales | Kytococcaceae | Kytococcus | sedentarius |
| SP00223 | Actinobacteria | Actinomycetia | Bifidobacteriales | Bifidobacteriaceae | Bifidobacterium | dentium |
| SP00224 | Actinobacteria | Actinomycetia | Micrococcales | Sanguibacteraceae | Sanguibacter | keddieii |
| SP00226 | Actinobacteria | Actinomycetia | Bifidobacteriales | Bifidobacteriaceae | Gardnerella | vaginalis |
| SP00227 | Proteobacteria | Gammaproteobacteria | Pseudomonadales | Moraxellaceae | Moraxella | catarrhalis |
| SP00229 | Actinobacteria | Coriobacteriia | Coriobacteriales | Atopobiaceae | Olsenella | uli |
| SP00230 | Bacteroidetes | Bacteroidia | Bacteroidales | Prevotellaceae | Prevotella | melaninogenica |
| SP00232 | Actinobacteria | Actinomycetia | Micrococcales | Micrococcaceae | Rothia | dentocariosa |
| SP00233 | Proteobacteria | Betaproteobacteria | Burkholderiales | Alcaligenaceae | Achromobacter | xylosoxidans |
| SP00235 | Firmicutes | Bacilli | Lactobacillales | Lactobacillaceae | Lactobacillus | amylovorus |
| SP00236 | Bacteroidetes | Bacteroidia | Bacteroidales | Prevotellaceae | Prevotella | denticola |
| SP00237 | Firmicutes | Bacilli | Lactobacillales | Lactobacillaceae | Lentilactobacillus | buchneri |
| SP00239 | Actinobacteria | Actinomycetia | Propionibacteriales | Propionibacteriaceae | Pseudopropionibacterium | propionicum |
| SP00240 | Proteobacteria | Gammaproteobacteria | Enterobacterales | Enterobacteriaceae | Klebsiella | aerogenes |
| SP00241 | Firmicutes | Bacilli | Lactobacillales | Streptococcaceae | Streptococcus | parasanguinis |
| SP00242 | Firmicutes | Bacilli | Lactobacillales | Streptococcaceae | Streptococcus | salivarius |
| SP00243 | Firmicutes | Bacilli | Lactobacillales | Lactobacillaceae | Lacticaseibacillus | rhamnosus |
| SP00244 | Bacteroidetes | Bacteroidia | Bacteroidales | Tannerellaceae | Tannerella | forsythia |
| SP00248 | Firmicutes | Bacilli | Bacillales | Staphylococcaceae | Staphylococcus | warneri |
| SP00249 | Tenericutes | Mollicutes | Mycoplasmatales | Mycoplasmataceae | Mycoplasma | genitalium |
| SP00250 | Proteobacteria | Betaproteobacteria | Burkholderiales | Burkholderiaceae | Burkholderia | cepacia |
| SP00251 | Firmicutes | Bacilli | Lactobacillales | Streptococcaceae | Streptococcus | constellatus |
| SP00252 | Firmicutes | Bacilli | Bacillales | Staphylococcaceae | Staphylococcus | pasteuri |
| SP00253 | Firmicutes | Bacilli | Lactobacillales | Streptococcaceae | Streptococcus | cristatus |
| SP00254 | Actinobacteria | Actinomycetia | Propionibacteriales | Propionibacteriaceae | Cutibacterium | avidum |
| SP00255 | Proteobacteria | Gammaproteobacteria | Enterobacterales | Enterobacteriaceae | Cronobacter | malonaticus |
| SP00256 | Actinobacteria | Actinomycetia | Corynebacteriales | Mycobacteriaceae | Mycolicibacterium | neoaurum |
| SP00258 | Firmicutes | Bacilli | Bacillales | Staphylococcaceae | Staphylococcus | capitis |
| SP00259 | Actinobacteria | Actinomycetia | Corynebacteriales | Corynebacteriaceae | Corynebacterium | sp. ATCC 6931 |
| SP00260 | Proteobacteria | Gammaproteobacteria | Pseudomonadales | Moraxellaceae | Acinetobacter | johnsonii |
| SP00261 | Proteobacteria | Betaproteobacteria | Burkholderiales | Burkholderiaceae | Cupriavidus | gilardii |
| SP00262 | Actinobacteria | Actinomycetia | Corynebacteriales | Corynebacteriaceae | Corynebacterium | singulare |
| SP00264 | Actinobacteria | Coriobacteriia | Coriobacteriales | Atopobiaceae | Olsenella | sp. oral taxon 807 |
| SP00265 | Actinobacteria | Actinomycetia | Micrococcales | Intrasporangiaceae | Arsenicicoccus | sp. oral taxon 190 |
| SP00266 | Firmicutes | Negativicutes | Selenomonadales | Selenomonadaceae | Selenomonas | sp. oral taxon 478 |
| SP00267 | Actinobacteria | Actinomycetia | Actinomycetales | Actinomycetaceae | Schaalia | meyeri |
| SP00268 | Proteobacteria | Betaproteobacteria | Burkholderiales | Comamonadaceae | Ottowia | sp. oral taxon 894 |
| SP00269 | Bacteroidetes | Bacteroidia | Bacteroidales | Prevotellaceae | Prevotella | fusca |
| SP00270 | Actinobacteria | Actinomycetia | Corynebacteriales | Lawsonellaceae | Lawsonella | clevelandensis |
| SP00271 | Fusobacteria | Fusobacteriia | Fusobacteriales | Leptotrichiaceae | Leptotrichia | sp. oral taxon 212 |
| SP00272 | Actinobacteria | Actinomycetia | Micrococcales | Micrococcaceae | Kocuria | palustris |
| SP00273 | Bacteroidetes | Flavobacteriia | Flavobacteriales | Flavobacteriaceae | Capnocytophaga | sp. oral taxon 323 |
| SP00275 | Proteobacteria | Gammaproteobacteria | Enterobacterales | Yersiniaceae | Serratia | marcescens |
| SP00276 | Bacteroidetes | Bacteroidia | Bacteroidales | Prevotellaceae | Prevotella | enoeca |
| SP00277 | Actinobacteria | Actinomycetia | Micrococcales | Intrasporangiaceae | Janibacter | indicus |
| SP00278 | Fusobacteria | Fusobacteriia | Fusobacteriales | Fusobacteriaceae | Fusobacterium | hwasookii |
| SP00279 | Firmicutes | Bacilli | Bacillales | Staphylococcaceae | Staphylococcus | haemolyticus |
| SP00280 | Firmicutes | Bacilli | Bacillales | Staphylococcaceae | Staphylococcus | lugdunensis |
| SP00282 | Actinobacteria | Actinomycetia | Actinomycetales | Actinomycetaceae | Actinomyces | radicidentis |
| SP00285 | Fusobacteria | Fusobacteriia | Fusobacteriales | Leptotrichiaceae | Leptotrichia | sp. oral taxon 847 |
| SP00286 | Actinobacteria | Actinomycetia | Actinomycetales | Actinomycetaceae | Actinomyces | oris |
| SP00287 | Proteobacteria | Gammaproteobacteria | Pseudomonadales | Moraxellaceae | Moraxella | osloensis |
| SP00288 | Firmicutes | Bacilli | Lactobacillales | Streptococcaceae | Streptococcus | sp. oral taxon 431 |
| SP00289 | Actinobacteria | Actinomycetia | Corynebacteriales | Corynebacteriaceae | Corynebacterium | simulans |
| SP00291 | Fusobacteria | Fusobacteriia | Fusobacteriales | Leptotrichiaceae | Leptotrichia | sp. oral taxon 498 |
| SP00293 | Bacteroidetes | Bacteroidia | Bacteroidales | Tannerellaceae | Tannerella | sp. oral taxon HOT-286 |
| SP00294 | Actinobacteria | Actinomycetia | Propionibacteriales | Propionibacteriaceae | Propionibacterium | sp. oral taxon 193 |
| SP00295 | Firmicutes | Negativicutes | Selenomonadales | Selenomonadaceae | Selenomonas | sp. oral taxon 920 |
| SP00296 | Proteobacteria | Gammaproteobacteria | Pseudomonadales | Moraxellaceae | Acinetobacter | junii |
| SP00297 | Firmicutes | Bacilli | Bacillales | Staphylococcaceae | Staphylococcus | cohnii |
| SP00298 | Proteobacteria | Alphaproteobacteria | Rhodobacterales | Rhodobacteraceae | Paracoccus | yeei |
| SP00300 | Actinobacteria | Actinomycetia | Corynebacteriales | Corynebacteriaceae | Corynebacterium | striatum |
| SP00301 | Actinobacteria | Actinomycetia | Micrococcales | Micrococcaceae | Kocuria | rhizophila |
| SP00302 | Firmicutes | Bacilli | Bacillales | Staphylococcaceae | Staphylococcus | pettenkoferi |
| SP00305 | Bacteroidetes | Flavobacteriia | Flavobacteriales | Flavobacteriaceae | Capnocytophaga | sp. oral taxon 878 |
| SP00306 | Bacteroidetes | Bacteroidia | Bacteroidales | Bacteroidaceae | Bacteroides | zoogleoformans |
| SP00307 | Bacteroidetes | Bacteroidia | Bacteroidales | Bacteroidaceae | Bacteroides | heparinolyticus |
| SP00308 | Proteobacteria | Gammaproteobacteria | Pasteurellales | Pasteurellaceae | Haemophilus | sp. oral taxon 036 |
| SP00309 | Actinobacteria | Actinomycetia | Actinomycetales | Actinomycetaceae | Actinomyces | sp. oral taxon 897 |
| SP00310 | Actinobacteria | Actinomycetia | Corynebacteriales | Dietziaceae | Dietzia | sp. oral taxon 368 |
| SP00312 | Actinobacteria | Actinomycetia | Actinomycetales | Actinomycetaceae | Actinomyces | sp. oral taxon 171 |
| SP00313 | Actinobacteria | Actinomycetia | Actinomycetales | Actinomycetaceae | Schaalia | odontolytica |
| SP00314 | Actinobacteria | Actinomycetia | Actinomycetales | Actinomycetaceae | Actinomyces | sp. oral taxon 169 |
| SP00315 | Firmicutes | Bacilli | Lactobacillales | Streptococcaceae | Streptococcus | mitis |
| SP00316 | Proteobacteria | Betaproteobacteria | Neisseriales | Neisseriaceae | Neisseria | lactamica |
| SP00318 | Proteobacteria | Gammaproteobacteria | Pasteurellales | Pasteurellaceae | Haemophilus | parainfluenzae |
| SP00319 | Firmicutes | Bacilli | Lactobacillales | Streptococcaceae | Streptococcus | oralis |
| SP00320 | Firmicutes | Bacilli | Lactobacillales | Streptococcaceae | Streptococcus | thermophilus |
| SP00321 | Spirochaetes | Spirochaetia | Spirochaetales | Spirochaetaceae | Treponema | putidum |

The Table illustrates the bacterial species which had amplicon similarity/identity values ≥97% with at least one different taxon using the bacterial-specific and the bacterial and archaeal primer pairs analyzed in the present study. ID= species identifier.

Appendix Table 11. Oral bacteria species with no amplicon similarity values ≥97% with different taxa.

| **ID** | **Phylum** | **Class** | **Order** | **Family** | **Genus** | **Species** |
| --- | --- | --- | --- | --- | --- | --- |
| SP00137 | Proteobacteria | Epsilonproteobacteria | Campylobacterales | Helicobacteraceae | Helicobacter | pylori |
| SP00139 | Spirochaetes | Spirochaetia | Spirochaetales | Spirochaetaceae | Treponema | pallidum |
| SP00140 | Chlamydiae | Chlamydiia | Chlamydiales | Chlamydiaceae | Chlamydia | pneumoniae |
| SP00145 | Firmicutes | Bacilli | Lactobacillales | Streptococcaceae | Lactococcus | lactis |
| SP00153 | Bacteroidetes | Bacteroidia | Bacteroidales | Porphyromonadaceae | Porphyromonas | gingivalis |
| SP00167 | Proteobacteria | Gammaproteobacteria | Xanthomonadales | Xanthomonadaceae | Stenotrophomonas | maltophilia |
| SP00171 | Firmicutes | Bacilli | Bacillales | Bacillaceae | Alkalihalobacillus | clausii |
| SP00175 | Firmicutes | Tissierellia | Tissierellales | Peptoniphilaceae | Finegoldia | magna |
| SP00176 | Tenericutes | Mollicutes | Mycoplasmatales | Mycoplasmataceae | Mycoplasmopsis | fermentans |
| SP00180 | Chloroflexi | Anaerolineae | Anaerolineales | Anaerolineaceae | Anaerolinea | thermophila |
| SP00183 | Chloroflexi | Caldilineae | Caldilineales | Caldilineaceae | Caldilinea | aerophila |
| SP00207 | Chlorobi | Chlorobia | Chlorobiales | Chlorobiaceae | Chlorobium | limicola |
| SP00217 | Actinobacteria | Coriobacteriia | Eggerthellales | Eggerthellaceae | Cryptobacterium | curtum |
| SP00220 | Firmicutes | Tissierellia | Tissierellales | Peptoniphilaceae | Anaerococcus | prevotii |
| SP00221 | Actinobacteria | Coriobacteriia | Coriobacteriales | Atopobiaceae | Lancefieldella | parvulum |
| SP00222 | Actinobacteria | Coriobacteriia | Eggerthellales | Eggerthellaceae | Eggerthella | lenta |
| SP00225 | Firmicutes | Negativicutes | Veillonellales | Veillonellaceae | Veillonella | parvula |
| SP00228 | Actinobacteria | Actinomycetia | Actinomycetales | Actinomycetaceae | Arcanobacterium | haemolyticum |
| SP00231 | Firmicutes | Clostridia | Eubacteriales | Eubacteriaceae | Eubacterium | callanderi |
| SP00234 | Firmicutes | Clostridia | Eubacteriales | Peptostreptococcaceae | Filifactor | alocis |
| SP00238 | Bacteroidetes | Bacteroidia | Bacteroidales | Porphyromonadaceae | Porphyromonas | asaccharolytica |
| SP00245 | Ignavibacteriae | Ignavibacteria | Ignavibacteriales | Ignavibacteriaceae | Ignavibacterium | album |
| SP00246 | Bacteroidetes | Bacteroidia | Bacteroidales | Prevotellaceae | Prevotella | intermedia |
| SP00247 | Ignavibacteriae | Ignavibacteria | Ignavibacteriales | Melioribacteraceae | Melioribacter | roseus |
| SP00257 | C.Saccharibacteria | C.Saccharimonia | C.Nanosynbacterales | C.Nanosynbacteraceae | C. Nanosynbacter | lyticus |
| SP00263 | Fusobacteria | Fusobacteriia | Fusobacteriales | Leptotrichiaceae | Sneathia | vaginalis |
| SP00274 | Actinobacteria | Actinomycetia | Actinomycetales | Actinomycetaceae | Actinomyces | sp. oral taxon 414 |
| SP00281 | Bacteroidetes | Flavobacteriia | Flavobacteriales | Flavobacteriaceae | Capnocytophaga | haemolytica |
| SP00283 | Proteobacteria | Deltaproteobacteria | Desulfovibrionales | Desulfovibrionaceae | Desulfovibrio | fairfieldensis |
| SP00284 | Proteobacteria | Deltaproteobacteria | Desulfovibrionales | Desulfomicrobiaceae | Desulfomicrobium | orale |
| SP00290 | Proteobacteria | Alphaproteobacteria | Rhodospirillales | Acetobacteraceae | Roseomonas | gilardii |
| SP00292 | Firmicutes | Negativicutes | Veillonellales | Veillonellaceae | Dialister | pneumosintes |
| SP00299 | Firmicutes | Bacilli | Bacillales | Bacillaceae | Anoxybacillus | flavithermus |
| SP00303 | Firmicutes | Clostridia | Eubacteriales | Oscillospiraceae | Fastidiosipila | sanguinis |
| SP00304 | Firmicutes | Clostridia | Eubacteriales | Clos. F. XIII. I. Sedis | Mogibacterium | diversum |
| SP00311 | Firmicutes | Bacilli | Bacillales | Paenibacillaceae | Paenibacillus | glucanolyticus |
| SP00317 | Tenericutes | Mollicutes | Mycoplasmatales | Mycoplasmataceae | Mycoplasma | hominis |

The Table details the bacterial species which had no amplicon similarity/identity values ≥97% with different taxa using the bacterial-specific and the bacterial and archaeal primer pairs analyzed in the present study. C.= candidatus; Clos. F. XIII. I. Sedis= clostridiales Family XIII. Incertae Sedis; ID= species identifier.

Appendix Table 12. Pairs of oral bacterial species with amplicon similarity ≥97% using the analyzed primer pairs.

| **ID** | **Genus** | **Species** | **ID** | **Genus** | **Species** | **Freq.** |
| --- | --- | --- | --- | --- | --- | --- |
| SP00141 | Neisseria | meningitidis | SP00144 | Neisseria | Gonorrhoeae | 29 |
| SP00146 | Streptococcus | pneumoniae | SP00288 | Streptococcus | sp. oral taxon 431 | 29 |
| SP00146 | Streptococcus | pneumoniae | SP00315 | Streptococcus | Mitis | 29 |
| SP00146 | Streptococcus | pneumoniae | SP00319 | Streptococcus | Oralis | 29 |
| SP00154 | Staphylococcus | epidermidis | SP00248 | Staphylococcus | Warneri | 29 |
| SP00154 | Staphylococcus | epidermidis | SP00252 | Staphylococcus | Pasteuri | 29 |
| SP00154 | Staphylococcus | epidermidis | SP00258 | Staphylococcus | Capitis | 29 |
| SP00158 | Lactobacillus | johnsonii | SP00195 | Lactobacillus | Gasseri | 29 |
| SP00242 | Streptococcus | salivarius | SP00320 | Streptococcus | Thermophilus | 29 |
| SP00248 | Staphylococcus | warneri | SP00252 | Staphylococcus | Pasteuri | 29 |
| SP00248 | Staphylococcus | warneri | SP00258 | Staphylococcus | Capitis | 29 |
| SP00252 | Staphylococcus | pasteuri | SP00258 | Staphylococcus | Capitis | 29 |
| SP00279 | Staphylococcus | haemolyticus | SP00280 | Staphylococcus | Lugdunensis | 29 |
| SP00279 | Staphylococcus | haemolyticus | SP00297 | Staphylococcus | Cohnii | 29 |
| SP00286 | Actinomyces | oris | SP00314 | Actinomyces | sp. oral taxon 169 | 29 |
| SP00288 | Streptococcus | sp. oral taxon 431 | SP00315 | Streptococcus | Mitis | 29 |
| SP00288 | Streptococcus | sp. oral taxon 431 | SP00319 | Streptococcus | Oralis | 29 |
| SP00315 | Streptococcus | mitis | SP00319 | Streptococcus | Oralis | 29 |
| SP00149 | Fusobacterium | nucleatum | SP00278 | Fusobacterium | Hwasookii | 28 |
| SP00154 | Staphylococcus | epidermidis | SP00163 | Staphylococcus | Aureus | 28 |
| SP00154 | Staphylococcus | epidermidis | SP00280 | Staphylococcus | Lugdunensis | 28 |
| SP00163 | Staphylococcus | aureus | SP00258 | Staphylococcus | Capitis | 28 |
| SP00205 | Klebsiella | variicola | SP00172 | Klebsiella | Pneumoniae | 28 |
| SP00240 | Klebsiella | aerogenes | SP00172 | Klebsiella | Pneumoniae | 28 |
| SP00189 | Lactobacillus | acidophilus | SP00235 | Lactobacillus | Amylovorus | 28 |
| SP00196 | Lacticaseibacillus | paracasei | SP00243 | Lacticaseibacillus | Rhamnosus | 28 |
| SP00202 | Cronobacter | sakazakii | SP00255 | Cronobacter | Malonaticus | 28 |
| SP00215 | Capnocytophaga | ochracea | SP00273 | Capnocytophaga | sp. oral taxon 323 | 28 |
| SP00280 | Staphylococcus | lugdunensis | SP00297 | Staphylococcus | Cohnii | 28 |
| SP00138 | Mycobacterium | tuberculosis | SP00164 | Mycobacterium | Leprae | 27 |
| SP00154 | Staphylococcus | epidermidis | SP00279 | Staphylococcus | Haemolyticus | 27 |
| SP00154 | Staphylococcus | epidermidis | SP00302 | Staphylococcus | Pettenkoferi | 27 |
| SP00159 | Treponema | denticola | SP00321 | Treponema | Putidum | 27 |
| SP00163 | Staphylococcus | aureus | SP00248 | Staphylococcus | Warneri | 27 |
| SP00163 | Staphylococcus | aureus | SP00252 | Staphylococcus | Pasteuri | 27 |
| SP00163 | Staphylococcus | aureus | SP00279 | Staphylococcus | Haemolyticus | 27 |
| SP00163 | Staphylococcus | aureus | SP00280 | Staphylococcus | Lugdunensis | 27 |
| SP00163 | Staphylococcus | aureus | SP00297 | Staphylococcus | Cohnii | 27 |
| SP00163 | Staphylococcus | aureus | SP00302 | Staphylococcus | Pettenkoferi | 27 |
| SP00166 | Pseudomonas | fluorescens | SP00191 | Pseudomonas | Protegens | 27 |
| SP00186 | Staphylococcus | schleiferi | SP00280 | Staphylococcus | lugdunensis | 27 |
| SP00194 | Streptococcus | sanguinis | SP00288 | Streptococcus | sp. oral taxon 431 | 27 |
| SP00194 | Streptococcus | sanguinis | SP00315 | Streptococcus | mitis | 27 |
| SP00194 | Streptococcus | sanguinis | SP00319 | Streptococcus | oralis | 27 |
| SP00248 | Staphylococcus | warneri | SP00280 | Staphylococcus | lugdunensis | 27 |
| SP00248 | Staphylococcus | warneri | SP00297 | Staphylococcus | cohnii | 27 |
| SP00252 | Staphylococcus | pasteuri | SP00280 | Staphylococcus | lugdunensis | 27 |
| SP00252 | Staphylococcus | pasteuri | SP00297 | Staphylococcus | cohnii | 27 |
| SP00258 | Staphylococcus | capitis | SP00280 | Staphylococcus | lugdunensis | 27 |
| SP00258 | Staphylococcus | capitis | SP00302 | Staphylococcus | pettenkoferi | 27 |
| SP00266 | Selenomonas | sp. oral taxon 478 | SP00295 | Selenomonas | sp. oral taxon 920 | 27 |
| SP00280 | Staphylococcus | lugdunensis | SP00302 | Staphylococcus | pettenkoferi | 27 |
| SP00286 | Actinomyces | oris | SP00312 | Actinomyces | sp. oral taxon 171 | 27 |
| SP00312 | Actinomyces | sp. oral taxon 171 | SP00314 | Actinomyces | sp. oral taxon 169 | 27 |
| SP00194 | Streptococcus | sanguinis | SP00146 | Streptococcus | pneumoniae | 26 |
| SP00251 | Streptococcus | constellatus | SP00177 | Streptococcus | intermedius | 26 |
| SP00205 | Klebsiella | variicola | SP00240 | Klebsiella | aerogenes | 26 |
| SP00258 | Staphylococcus | capitis | SP00279 | Staphylococcus | haemolyticus | 26 |
| SP00279 | Staphylococcus | haemolyticus | SP00302 | Staphylococcus | pettenkoferi | 26 |
| SP00297 | Staphylococcus | cohnii | SP00302 | Staphylococcus | pettenkoferi | 26 |
| SP00154 | Staphylococcus | epidermidis | SP00186 | Staphylococcus | schleiferi | 25 |
| SP00154 | Staphylococcus | epidermidis | SP00297 | Staphylococcus | cohnii | 25 |
| SP00186 | Staphylococcus | schleiferi | SP00163 | Staphylococcus | aureus | 25 |
| SP00166 | Pseudomonas | fluorescens | SP00193 | Pseudomonas | stutzeri | 25 |
| SP00186 | Staphylococcus | schleiferi | SP00248 | Staphylococcus | warneri | 25 |
| SP00186 | Staphylococcus | schleiferi | SP00252 | Staphylococcus | pasteuri | 25 |
| SP00186 | Staphylococcus | schleiferi | SP00258 | Staphylococcus | capitis | 25 |
| SP00188 | Bordetella | pertussis | SP00233 | Achromobacter | xylosoxidans | 25 |
| SP00194 | Streptococcus | sanguinis | SP00199 | Streptococcus | gordonii | 25 |
| SP00197 | Acinetobacter | baumannii | SP00296 | Acinetobacter | junii | 25 |
| SP00229 | Olsenella | uli | SP00264 | Olsenella | sp. oral taxon 807 | 25 |
| SP00241 | Streptococcus | parasanguinis | SP00315 | Streptococcus | mitis | 25 |
| SP00241 | Streptococcus | parasanguinis | SP00319 | Streptococcus | oralis | 25 |
| SP00248 | Staphylococcus | warneri | SP00279 | Staphylococcus | haemolyticus | 25 |
| SP00248 | Staphylococcus | warneri | SP00302 | Staphylococcus | pettenkoferi | 25 |
| SP00252 | Staphylococcus | pasteuri | SP00279 | Staphylococcus | haemolyticus | 25 |
| SP00252 | Staphylococcus | pasteuri | SP00302 | Staphylococcus | pettenkoferi | 25 |
| SP00258 | Staphylococcus | capitis | SP00297 | Staphylococcus | cohnii | 25 |
| SP00194 | Streptococcus | sanguinis | SP00241 | Streptococcus | parasanguinis | 24 |
| SP00194 | Streptococcus | sanguinis | SP00251 | Streptococcus | constellatus | 24 |
| SP00241 | Streptococcus | parasanguinis | SP00288 | Streptococcus | sp. oral taxon 431 | 24 |
| SP00262 | Corynebacterium | singulare | SP00289 | Corynebacterium | simulans | 24 |
| SP00273 | Capnocytophaga | sp. oral taxon 323 | SP00305 | Capnocytophaga | sp. oral taxon 878 | 24 |
| SP00289 | Corynebacterium | simulans | SP00300 | Corynebacterium | striatum | 24 |
| SP00316 | Neisseria | lactamica | SP00141 | Neisseria | meningitidis | 23 |
| SP00172 | Klebsiella | pneumoniae | SP00275 | Serratia | marcescens | 23 |
| SP00184 | Streptococcus | anginosus | SP00177 | Streptococcus | intermedius | 23 |
| SP00186 | Staphylococcus | schleiferi | SP00279 | Staphylococcus | haemolyticus | 23 |
| SP00186 | Staphylococcus | schleiferi | SP00297 | Staphylococcus | cohnii | 23 |
| SP00186 | Staphylococcus | schleiferi | SP00302 | Staphylococcus | pettenkoferi | 23 |
| SP00199 | Streptococcus | gordonii | SP00288 | Streptococcus | sp. oral taxon 431 | 23 |
| SP00199 | Streptococcus | gordonii | SP00315 | Streptococcus | mitis | 23 |
| SP00199 | Streptococcus | gordonii | SP00319 | Streptococcus | oralis | 23 |
| SP00205 | Klebsiella | variicola | SP00275 | Serratia | marcescens | 23 |
| SP00206 | Ralstonia | pickettii | SP00261 | Cupriavidus | gilardii | 23 |
| SP00215 | Capnocytophaga | ochracea | SP00305 | Capnocytophaga | sp. oral taxon 878 | 23 |
| SP00142 | Pseudomonas | aeruginosa | SP00193 | Pseudomonas | stutzeri | 22 |
| SP00146 | Streptococcus | pneumoniae | SP00241 | Streptococcus | parasanguinis | 22 |
| SP00184 | Streptococcus | anginosus | SP00251 | Streptococcus | constellatus | 22 |
| SP00241 | Streptococcus | parasanguinis | SP00253 | Streptococcus | cristatus | 22 |
| SP00260 | Acinetobacter | johnsonii | SP00296 | Acinetobacter | junii | 22 |
| SP00267 | Schaalia | meyeri | SP00313 | Schaalia | odontolytica | 22 |
| SP00199 | Streptococcus | gordonii | SP00146 | Streptococcus | pneumoniae | 21 |
| SP00194 | Streptococcus | sanguinis | SP00177 | Streptococcus | intermedius | 21 |
| SP00199 | Streptococcus | gordonii | SP00241 | Streptococcus | parasanguinis | 21 |
| SP00240 | Klebsiella | aerogenes | SP00275 | Serratia | marcescens | 21 |
| SP00136 | Escherichia | coli | SP00275 | Serratia | marcescens | 20 |
| SP00199 | Streptococcus | gordonii | SP00177 | Streptococcus | intermedius | 20 |
| SP00199 | Streptococcus | gordonii | SP00253 | Streptococcus | cristatus | 20 |
| SP00202 | Cronobacter | sakazakii | SP00240 | Klebsiella | aerogenes | 20 |
| SP00262 | Corynebacterium | singulare | SP00300 | Corynebacterium | striatum | 20 |
| SP00308 | Haemophilus | sp. oral taxon 036 | SP00190 | Haemophilus | influenzae | 19 |
| SP00201 | Campylobacter | curvus | SP00203 | Campylobacter | concisus | 19 |
| SP00202 | Cronobacter | sakazakii | SP00136 | Escherichia | coli | 18 |
| SP00136 | Escherichia | coli | SP00255 | Cronobacter | malonaticus | 18 |
| SP00144 | Neisseria | gonorrhoeae | SP00316 | Neisseria | lactamica | 18 |
| SP00202 | Cronobacter | sakazakii | SP00172 | Klebsiella | pneumoniae | 18 |
| SP00172 | Klebsiella | pneumoniae | SP00255 | Cronobacter | malonaticus | 18 |
| SP00178 | Rothia | mucilaginosa | SP00232 | Rothia | dentocariosa | 18 |
| SP00197 | Acinetobacter | baumannii | SP00260 | Acinetobacter | johnsonii | 18 |
| SP00209 | Comamonas | thiooxydans | SP00211 | Acidovorax | ebreus | 18 |
| SP00212 | Aggregatibacter | aphrophilus | SP00308 | Haemophilus | sp. oral taxon 036 | 18 |
| SP00241 | Streptococcus | parasanguinis | SP00251 | Streptococcus | constellatus | 18 |
| SP00253 | Streptococcus | cristatus | SP00288 | Streptococcus | sp. oral taxon 431 | 18 |
| SP00272 | Kocuria | palustris | SP00301 | Kocuria | rhizophila | 18 |
| SP00152 | Bifidobacterium | longum | SP00192 | Bifidobacterium | breve | 17 |
| SP00161 | Cutibacterium | acnes | SP00294 | Propionibacterium | sp. oral taxon 193 | 17 |
| SP00191 | Pseudomonas | protegens | SP00193 | Pseudomonas | stutzeri | 17 |
| SP00253 | Streptococcus | cristatus | SP00315 | Streptococcus | mitis | 17 |
| SP00253 | Streptococcus | cristatus | SP00319 | Streptococcus | oralis | 17 |
| SP00172 | Klebsiella | pneumoniae | SP00136 | Escherichia | coli | 16 |
| SP00146 | Streptococcus | pneumoniae | SP00253 | Streptococcus | cristatus | 16 |
| SP00150 | Yersinia | pestis | SP00275 | Serratia | marcescens | 16 |
| SP00240 | Klebsiella | aerogenes | SP00255 | Cronobacter | malonaticus | 16 |
| SP00148 | Streptococcus | agalactiae | SP00143 | Streptococcus | pyogenes | 15 |
| SP00241 | Streptococcus | parasanguinis | SP00177 | Streptococcus | intermedius | 15 |
| SP00194 | Streptococcus | sanguinis | SP00253 | Streptococcus | cristatus | 15 |
| SP00205 | Klebsiella | variicola | SP00136 | Escherichia | coli | 14 |
| SP00177 | Streptococcus | intermedius | SP00253 | Streptococcus | cristatus | 14 |
| SP00181 | Levilactobacillus | brevis | SP00237 | Lentilactobacillus | buchneri | 14 |
| SP00184 | Streptococcus | anginosus | SP00194 | Streptococcus | sanguinis | 14 |
| SP00199 | Streptococcus | gordonii | SP00251 | Streptococcus | constellatus | 14 |
| SP00202 | Cronobacter | sakazakii | SP00205 | Klebsiella | variicola | 14 |
| SP00184 | Streptococcus | anginosus | SP00199 | Streptococcus | gordonii | 13 |
| SP00184 | Streptococcus | anginosus | SP00253 | Streptococcus | cristatus | 13 |
| SP00205 | Klebsiella | variicola | SP00255 | Cronobacter | malonaticus | 13 |
| SP00210 | Rhodobacter | capsulatus | SP00298 | Paracoccus | yeei | 13 |
| SP00219 | Kytococcus | sedentarius | SP00277 | Janibacter | indicus | 13 |
| SP00138 | Mycobacterium | tuberculosis | SP00256 | Mycolicibacterium | neoaurum | 12 |
| SP00177 | Streptococcus | intermedius | SP00143 | Streptococcus | pyogenes | 11 |
| SP00199 | Streptococcus | gordonii | SP00143 | Streptococcus | pyogenes | 11 |
| SP00242 | Streptococcus | salivarius | SP00177 | Streptococcus | intermedius | 11 |
| SP00182 | Mycoplasma | pneumoniae | SP00249 | Mycoplasma | genitalium | 11 |
| SP00199 | Streptococcus | gordonii | SP00320 | Streptococcus | thermophilus | 11 |
| SP00202 | Cronobacter | sakazakii | SP00275 | Serratia | marcescens | 11 |
| SP00251 | Streptococcus | constellatus | SP00315 | Streptococcus | mitis | 11 |
| SP00251 | Streptococcus | constellatus | SP00319 | Streptococcus | oralis | 11 |
| SP00255 | Cronobacter | malonaticus | SP00275 | Serratia | marcescens | 11 |
| SP00282 | Actinomyces | radicidentis | SP00312 | Actinomyces | sp. oral taxon 171 | 11 |
| SP00282 | Actinomyces | radicidentis | SP00314 | Actinomyces | sp. oral taxon 169 | 11 |
| SP00152 | Bifidobacterium | longum | SP00223 | Bifidobacterium | dentium | 10 |
| SP00242 | Streptococcus | salivarius | SP00184 | Streptococcus | anginosus | 10 |
| SP00184 | Streptococcus | anginosus | SP00320 | Streptococcus | thermophilus | 10 |
| SP00230 | Prevotella | melaninogenica | SP00269 | Prevotella | fusca | 10 |
| SP00242 | Streptococcus | salivarius | SP00253 | Streptococcus | cristatus | 10 |
| SP00251 | Streptococcus | constellatus | SP00253 | Streptococcus | cristatus | 10 |
| SP00251 | Streptococcus | constellatus | SP00288 | Streptococcus | sp. oral taxon 431 | 10 |
| SP00136 | Escherichia | coli | SP00240 | Klebsiella | aerogenes | 9 |
| SP00251 | Streptococcus | constellatus | SP00146 | Streptococcus | pneumoniae | 9 |
| SP00148 | Streptococcus | agalactiae | SP00194 | Streptococcus | sanguinis | 9 |
| SP00150 | Yersinia | pestis | SP00240 | Klebsiella | aerogenes | 9 |
| SP00164 | Mycobacterium | leprae | SP00256 | Mycolicibacterium | neoaurum | 9 |
| SP00177 | Streptococcus | intermedius | SP00320 | Streptococcus | thermophilus | 9 |
| SP00192 | Bifidobacterium | breve | SP00223 | Bifidobacterium | dentium | 9 |
| SP00194 | Streptococcus | sanguinis | SP00242 | Streptococcus | salivarius | 9 |
| SP00194 | Streptococcus | sanguinis | SP00320 | Streptococcus | thermophilus | 9 |
| SP00199 | Streptococcus | gordonii | SP00242 | Streptococcus | salivarius | 9 |
| SP00253 | Streptococcus | cristatus | SP00320 | Streptococcus | thermophilus | 9 |
| SP00282 | Actinomyces | radicidentis | SP00286 | Actinomyces | oris | 9 |
| SP00194 | Streptococcus | sanguinis | SP00143 | Streptococcus | pyogenes | 8 |
| SP00184 | Streptococcus | anginosus | SP00241 | Streptococcus | parasanguinis | 8 |
| SP00187 | Corynebacterium | diphtheriae | SP00262 | Corynebacterium | singulare | 8 |
| SP00187 | Corynebacterium | diphtheriae | SP00289 | Corynebacterium | simulans | 8 |
| SP00212 | Aggregatibacter | aphrophilus | SP00190 | Haemophilus | influenzae | 8 |
| SP00211 | Acidovorax | ebreus | SP00216 | Variovorax | paradoxus | 8 |
| SP00241 | Streptococcus | parasanguinis | SP00242 | Streptococcus | salivarius | 8 |
| SP00241 | Streptococcus | parasanguinis | SP00320 | Streptococcus | thermophilus | 8 |
| SP00244 | Tannerella | forsythia | SP00293 | Tannerella | sp. oral taxon HOT-286 | 8 |
| SP00136 | Escherichia | coli | SP00150 | Yersinia | pestis | 7 |
| SP00146 | Streptococcus | pneumoniae | SP00177 | Streptococcus | intermedius | 7 |
| SP00150 | Yersinia | pestis | SP00172 | Klebsiella | pneumoniae | 7 |
| SP00150 | Yersinia | pestis | SP00202 | Cronobacter | sakazakii | 7 |
| SP00150 | Yersinia | pestis | SP00205 | Klebsiella | variicola | 7 |
| SP00150 | Yersinia | pestis | SP00255 | Cronobacter | malonaticus | 7 |
| SP00161 | Cutibacterium | acnes | SP00254 | Cutibacterium | avidum | 7 |
| SP00165 | Lactiplantibacillus | plantarum | SP00181 | Levilactobacillus | brevis | 7 |
| SP00168 | Corynebacterium | urealyticum | SP00262 | Corynebacterium | singulare | 7 |
| SP00173 | Limosilactobacillus | reuteri | SP00174 | Limosilactobacillus | fermentum | 7 |
| SP00173 | Limosilactobacillus | reuteri | SP00237 | Lentilactobacillus | buchneri | 7 |
| SP00209 | Comamonas | thiooxydans | SP00216 | Variovorax | paradoxus | 7 |
| SP00254 | Cutibacterium | avidum | SP00294 | Propionibacterium | sp. oral taxon 193 | 7 |
| SP00152 | Bifidobacterium | longum | SP00208 | Bifidobacterium | animalis | 6 |
| SP00165 | Lactiplantibacillus | plantarum | SP00196 | Lacticaseibacillus | paracasei | 6 |
| SP00165 | Lactiplantibacillus | plantarum | SP00237 | Lentilactobacillus | buchneri | 6 |
| SP00165 | Lactiplantibacillus | plantarum | SP00243 | Lacticaseibacillus | rhamnosus | 6 |
| SP00181 | Levilactobacillus | brevis | SP00243 | Lacticaseibacillus | rhamnosus | 6 |
| SP00318 | Haemophilus | parainfluenzae | SP00190 | Haemophilus | influenzae | 6 |
| SP00204 | Delftia | acidovorans | SP00216 | Variovorax | paradoxus | 6 |
| SP00223 | Bifidobacterium | dentium | SP00208 | Bifidobacterium | animalis | 6 |
| SP00237 | Lentilactobacillus | buchneri | SP00243 | Lacticaseibacillus | rhamnosus | 6 |
| SP00242 | Streptococcus | salivarius | SP00251 | Streptococcus | constellatus | 6 |
| SP00251 | Streptococcus | constellatus | SP00320 | Streptococcus | thermophilus | 6 |
| SP00306 | Bacteroides | zoogleoformans | SP00307 | Bacteroides | heparinolyticus | 6 |
| SP00143 | Streptococcus | pyogenes | SP00288 | Streptococcus | sp. oral taxon 431 | 5 |
| SP00143 | Streptococcus | pyogenes | SP00315 | Streptococcus | mitis | 5 |
| SP00143 | Streptococcus | pyogenes | SP00319 | Streptococcus | oralis | 5 |
| SP00148 | Streptococcus | agalactiae | SP00199 | Streptococcus | gordonii | 5 |
| SP00148 | Streptococcus | agalactiae | SP00315 | Streptococcus | mitis | 5 |
| SP00148 | Streptococcus | agalactiae | SP00319 | Streptococcus | oralis | 5 |
| SP00151 | Streptococcus | mutans | SP00184 | Streptococcus | anginosus | 5 |
| SP00177 | Streptococcus | intermedius | SP00315 | Streptococcus | mitis | 5 |
| SP00177 | Streptococcus | intermedius | SP00319 | Streptococcus | oralis | 5 |
| SP00181 | Levilactobacillus | brevis | SP00196 | Lacticaseibacillus | paracasei | 5 |
| SP00185 | Aggregatibacter | actinomycetemcomitans | SP00212 | Aggregatibacter | aphrophilus | 5 |
| SP00187 | Corynebacterium | diphtheriae | SP00300 | Corynebacterium | striatum | 5 |
| SP00192 | Bifidobacterium | breve | SP00208 | Bifidobacterium | animalis | 5 |
| SP00196 | Lacticaseibacillus | paracasei | SP00237 | Lentilactobacillus | buchneri | 5 |
| SP00308 | Haemophilus | sp. oral taxon 036 | SP00318 | Haemophilus | parainfluenzae | 5 |
| SP00143 | Streptococcus | pyogenes | SP00146 | Streptococcus | pneumoniae | 4 |
| SP00143 | Streptococcus | pyogenes | SP00151 | Streptococcus | mutans | 4 |
| SP00143 | Streptococcus | pyogenes | SP00184 | Streptococcus | anginosus | 4 |
| SP00143 | Streptococcus | pyogenes | SP00251 | Streptococcus | constellatus | 4 |
| SP00147 | Agrobacterium | fabrum | SP00198 | Agrobacterium | radiobacter | 4 |
| SP00147 | Agrobacterium | fabrum | SP00200 | Brucella | anthropi | 4 |
| SP00148 | Streptococcus | agalactiae | SP00177 | Streptococcus | intermedius | 4 |
| SP00148 | Streptococcus | agalactiae | SP00251 | Streptococcus | constellatus | 4 |
| SP00148 | Streptococcus | agalactiae | SP00288 | Streptococcus | sp. oral taxon 431 | 4 |
| SP00151 | Streptococcus | mutans | SP00199 | Streptococcus | gordonii | 4 |
| SP00155 | Enterococcus | faecalis | SP00160 | Listeria | monocytogenes | 4 |
| SP00157 | Haemophilus | ducreyi | SP00318 | Haemophilus | parainfluenzae | 4 |
| SP00187 | Corynebacterium | diphtheriae | SP00168 | Corynebacterium | urealyticum | 4 |
| SP00198 | Agrobacterium | radiobacter | SP00200 | Brucella | anthropi | 4 |
| SP00214 | Micrococcus | luteus | SP00272 | Kocuria | palustris | 4 |
| SP00218 | Leptotrichia | buccalis | SP00291 | Leptotrichia | sp. oral taxon 498 | 4 |
| SP00224 | Sanguibacter | keddieii | SP00277 | Janibacter | indicus | 4 |
| SP00146 | Streptococcus | pneumoniae | SP00148 | Streptococcus | agalactiae | 3 |
| SP00151 | Streptococcus | mutans | SP00148 | Streptococcus | agalactiae | 3 |
| SP00148 | Streptococcus | agalactiae | SP00184 | Streptococcus | anginosus | 3 |
| SP00148 | Streptococcus | agalactiae | SP00241 | Streptococcus | parasanguinis | 3 |
| SP00151 | Streptococcus | mutans | SP00242 | Streptococcus | salivarius | 3 |
| SP00151 | Streptococcus | mutans | SP00320 | Streptococcus | thermophilus | 3 |
| SP00156 | Bacillus | anthracis | SP00179 | Bacillus | subtilis | 3 |
| SP00156 | Bacillus | anthracis | SP00297 | Staphylococcus | cohnii | 3 |
| SP00157 | Haemophilus | ducreyi | SP00190 | Haemophilus | influenzae | 3 |
| SP00157 | Haemophilus | ducreyi | SP00308 | Haemophilus | sp. oral taxon 036 | 3 |
| SP00158 | Lactobacillus | johnsonii | SP00189 | Lactobacillus | acidophilus | 3 |
| SP00158 | Lactobacillus | johnsonii | SP00235 | Lactobacillus | amylovorus | 3 |
| SP00160 | Listeria | monocytogenes | SP00163 | Staphylococcus | aureus | 3 |
| SP00160 | Listeria | monocytogenes | SP00280 | Staphylococcus | lugdunensis | 3 |
| SP00160 | Listeria | monocytogenes | SP00297 | Staphylococcus | cohnii | 3 |
| SP00168 | Corynebacterium | urealyticum | SP00289 | Corynebacterium | simulans | 3 |
| SP00168 | Corynebacterium | urealyticum | SP00300 | Corynebacterium | striatum | 3 |
| SP00170 | Mesorhizobium | japonicum | SP00198 | Agrobacterium | radiobacter | 3 |
| SP00170 | Mesorhizobium | japonicum | SP00200 | Brucella | anthropi | 3 |
| SP00181 | Levilactobacillus | brevis | SP00173 | Limosilactobacillus | reuteri | 3 |
| SP00173 | Limosilactobacillus | reuteri | SP00243 | Lacticaseibacillus | rhamnosus | 3 |
| SP00177 | Streptococcus | intermedius | SP00288 | Streptococcus | sp. oral taxon 431 | 3 |
| SP00178 | Rothia | mucilaginosa | SP00272 | Kocuria | palustris | 3 |
| SP00187 | Corynebacterium | diphtheriae | SP00259 | Corynebacterium | sp. ATCC 6931 | 3 |
| SP00189 | Lactobacillus | acidophilus | SP00195 | Lactobacillus | gasseri | 3 |
| SP00195 | Lactobacillus | gasseri | SP00235 | Lactobacillus | amylovorus | 3 |
| SP00204 | Delftia | acidovorans | SP00209 | Comamonas | thiooxydans | 3 |
| SP00213 | Corynebacterium | kroppenstedtii | SP00259 | Corynebacterium | sp. ATCC 6931 | 3 |
| SP00214 | Micrococcus | luteus | SP00224 | Sanguibacter | keddieii | 3 |
| SP00224 | Sanguibacter | keddieii | SP00265 | Arsenicicoccus | sp. oral taxon 190 | 3 |
| SP00259 | Corynebacterium | sp. ATCC 6931 | SP00262 | Corynebacterium | singulare | 3 |
| SP00259 | Corynebacterium | sp. ATCC 6931 | SP00289 | Corynebacterium | simulans | 3 |
| SP00259 | Corynebacterium | sp. ATCC 6931 | SP00300 | Corynebacterium | striatum | 3 |
| SP00265 | Arsenicicoccus | sp. oral taxon 190 | SP00277 | Janibacter | indicus | 3 |
| SP00267 | Schaalia | meyeri | SP00282 | Actinomyces | radicidentis | 3 |
| SP00267 | Schaalia | meyeri | SP00286 | Actinomyces | oris | 3 |
| SP00267 | Schaalia | meyeri | SP00312 | Actinomyces | sp. oral taxon 171 | 3 |
| SP00267 | Schaalia | meyeri | SP00314 | Actinomyces | sp. oral taxon 169 | 3 |
| SP00286 | Actinomyces | oris | SP00313 | Schaalia | odontolytica | 3 |
| SP00312 | Actinomyces | sp. oral taxon 171 | SP00313 | Schaalia | odontolytica | 3 |
| SP00313 | Schaalia | odontolytica | SP00314 | Actinomyces | sp. oral taxon 169 | 3 |
| SP00136 | Escherichia | coli | SP00318 | Haemophilus | parainfluenzae | 2 |
| SP00143 | Streptococcus | pyogenes | SP00241 | Streptococcus | parasanguinis | 2 |
| SP00143 | Streptococcus | pyogenes | SP00242 | Streptococcus | salivarius | 2 |
| SP00143 | Streptococcus | pyogenes | SP00253 | Streptococcus | cristatus | 2 |
| SP00143 | Streptococcus | pyogenes | SP00320 | Streptococcus | thermophilus | 2 |
| SP00151 | Streptococcus | mutans | SP00177 | Streptococcus | intermedius | 2 |
| SP00151 | Streptococcus | mutans | SP00194 | Streptococcus | sanguinis | 2 |
| SP00151 | Streptococcus | mutans | SP00251 | Streptococcus | constellatus | 2 |
| SP00154 | Staphylococcus | epidermidis | SP00160 | Listeria | monocytogenes | 2 |
| SP00154 | Staphylococcus | epidermidis | SP00179 | Bacillus | subtilis | 2 |
| SP00181 | Levilactobacillus | brevis | SP00155 | Enterococcus | faecalis | 2 |
| SP00155 | Enterococcus | faecalis | SP00237 | Lentilactobacillus | buchneri | 2 |
| SP00155 | Enterococcus | faecalis | SP00297 | Staphylococcus | cohnii | 2 |
| SP00160 | Listeria | monocytogenes | SP00179 | Bacillus | subtilis | 2 |
| SP00160 | Listeria | monocytogenes | SP00186 | Staphylococcus | schleiferi | 2 |
| SP00160 | Listeria | monocytogenes | SP00248 | Staphylococcus | warneri | 2 |
| SP00160 | Listeria | monocytogenes | SP00252 | Staphylococcus | pasteuri | 2 |
| SP00160 | Listeria | monocytogenes | SP00258 | Staphylococcus | capitis | 2 |
| SP00160 | Listeria | monocytogenes | SP00279 | Staphylococcus | haemolyticus | 2 |
| SP00160 | Listeria | monocytogenes | SP00302 | Staphylococcus | pettenkoferi | 2 |
| SP00196 | Lacticaseibacillus | paracasei | SP00162 | Ligilactobacillus | salivarius | 2 |
| SP00162 | Ligilactobacillus | salivarius | SP00243 | Lacticaseibacillus | rhamnosus | 2 |
| SP00163 | Staphylococcus | aureus | SP00179 | Bacillus | subtilis | 2 |
| SP00172 | Klebsiella | pneumoniae | SP00318 | Haemophilus | parainfluenzae | 2 |
| SP00173 | Limosilactobacillus | reuteri | SP00196 | Lacticaseibacillus | paracasei | 2 |
| SP00174 | Limosilactobacillus | fermentum | SP00181 | Levilactobacillus | brevis | 2 |
| SP00174 | Limosilactobacillus | fermentum | SP00196 | Lacticaseibacillus | paracasei | 2 |
| SP00174 | Limosilactobacillus | fermentum | SP00237 | Lentilactobacillus | buchneri | 2 |
| SP00174 | Limosilactobacillus | fermentum | SP00243 | Lacticaseibacillus | rhamnosus | 2 |
| SP00186 | Staphylococcus | schleiferi | SP00179 | Bacillus | subtilis | 2 |
| SP00179 | Bacillus | subtilis | SP00248 | Staphylococcus | warneri | 2 |
| SP00179 | Bacillus | subtilis | SP00252 | Staphylococcus | pasteuri | 2 |
| SP00179 | Bacillus | subtilis | SP00258 | Staphylococcus | capitis | 2 |
| SP00179 | Bacillus | subtilis | SP00279 | Staphylococcus | haemolyticus | 2 |
| SP00179 | Bacillus | subtilis | SP00280 | Staphylococcus | lugdunensis | 2 |
| SP00179 | Bacillus | subtilis | SP00302 | Staphylococcus | pettenkoferi | 2 |
| SP00184 | Streptococcus | anginosus | SP00288 | Streptococcus | sp. oral taxon 431 | 2 |
| SP00184 | Streptococcus | anginosus | SP00315 | Streptococcus | mitis | 2 |
| SP00184 | Streptococcus | anginosus | SP00319 | Streptococcus | oralis | 2 |
| SP00190 | Haemophilus | influenzae | SP00185 | Aggregatibacter | actinomycetemcomitans | 2 |
| SP00204 | Delftia | acidovorans | SP00211 | Acidovorax | ebreus | 2 |
| SP00204 | Delftia | acidovorans | SP00268 | Ottowia | sp. oral taxon 894 | 2 |
| SP00205 | Klebsiella | variicola | SP00318 | Haemophilus | parainfluenzae | 2 |
| SP00250 | Burkholderia | cepacia | SP00206 | Ralstonia | pickettii | 2 |
| SP00214 | Micrococcus | luteus | SP00219 | Kytococcus | sedentarius | 2 |
| SP00214 | Micrococcus | luteus | SP00270 | Lawsonella | clevelandensis | 2 |
| SP00214 | Micrococcus | luteus | SP00277 | Janibacter | indicus | 2 |
| SP00214 | Micrococcus | luteus | SP00301 | Kocuria | rhizophila | 2 |
| SP00218 | Leptotrichia | buccalis | SP00271 | Leptotrichia | sp. oral taxon 212 | 2 |
| SP00218 | Leptotrichia | buccalis | SP00285 | Leptotrichia | sp. oral taxon 847 | 2 |
| SP00219 | Kytococcus | sedentarius | SP00265 | Arsenicicoccus | sp. oral taxon 190 | 2 |
| SP00224 | Sanguibacter | keddieii | SP00270 | Lawsonella | clevelandensis | 2 |
| SP00224 | Sanguibacter | keddieii | SP00272 | Kocuria | palustris | 2 |
| SP00227 | Moraxella | catarrhalis | SP00287 | Moraxella | osloensis | 2 |
| SP00232 | Rothia | dentocariosa | SP00272 | Kocuria | palustris | 2 |
| SP00236 | Prevotella | denticola | SP00269 | Prevotella | fusca | 2 |
| SP00240 | Klebsiella | aerogenes | SP00318 | Haemophilus | parainfluenzae | 2 |
| SP00242 | Streptococcus | salivarius | SP00315 | Streptococcus | mitis | 2 |
| SP00242 | Streptococcus | salivarius | SP00319 | Streptococcus | oralis | 2 |
| SP00250 | Burkholderia | cepacia | SP00261 | Cupriavidus | gilardii | 2 |
| SP00259 | Corynebacterium | sp. ATCC 6931 | SP00310 | Dietzia | sp. oral taxon 368 | 2 |
| SP00270 | Lawsonella | clevelandensis | SP00277 | Janibacter | indicus | 2 |
| SP00271 | Leptotrichia | sp. oral taxon 212 | SP00291 | Leptotrichia | sp. oral taxon 498 | 2 |
| SP00272 | Kocuria | palustris | SP00277 | Janibacter | indicus | 2 |
| SP00275 | Serratia | marcescens | SP00318 | Haemophilus | parainfluenzae | 2 |
| SP00277 | Janibacter | indicus | SP00301 | Kocuria | rhizophila | 2 |
| SP00285 | Leptotrichia | sp. oral taxon 847 | SP00291 | Leptotrichia | sp. oral taxon 498 | 2 |
| SP00286 | Actinomyces | oris | SP00309 | Actinomyces | sp. oral taxon 897 | 2 |
| SP00309 | Actinomyces | sp. oral taxon 897 | SP00312 | Actinomyces | sp. oral taxon 171 | 2 |
| SP00309 | Actinomyces | sp. oral taxon 897 | SP00314 | Actinomyces | sp. oral taxon 169 | 2 |
| SP00315 | Streptococcus | mitis | SP00320 | Streptococcus | thermophilus | 2 |
| SP00319 | Streptococcus | oralis | SP00320 | Streptococcus | thermophilus | 2 |
| SP00142 | Pseudomonas | aeruginosa | SP00166 | Pseudomonas | fluorescens | 1 |
| SP00142 | Pseudomonas | aeruginosa | SP00191 | Pseudomonas | protegens | 1 |
| SP00184 | Streptococcus | anginosus | SP00146 | Streptococcus | pneumoniae | 1 |
| SP00146 | Streptococcus | pneumoniae | SP00242 | Streptococcus | salivarius | 1 |
| SP00146 | Streptococcus | pneumoniae | SP00320 | Streptococcus | thermophilus | 1 |
| SP00147 | Agrobacterium | fabrum | SP00170 | Mesorhizobium | japonicum | 1 |
| SP00148 | Streptococcus | agalactiae | SP00242 | Streptococcus | salivarius | 1 |
| SP00148 | Streptococcus | agalactiae | SP00253 | Streptococcus | cristatus | 1 |
| SP00148 | Streptococcus | agalactiae | SP00320 | Streptococcus | thermophilus | 1 |
| SP00169 | Proteus | mirabilis | SP00150 | Yersinia | pestis | 1 |
| SP00151 | Streptococcus | mutans | SP00288 | Streptococcus | sp. oral taxon 431 | 1 |
| SP00151 | Streptococcus | mutans | SP00315 | Streptococcus | mitis | 1 |
| SP00151 | Streptococcus | mutans | SP00319 | Streptococcus | oralis | 1 |
| SP00152 | Bifidobacterium | longum | SP00226 | Gardnerella | vaginalis | 1 |
| SP00155 | Enterococcus | faecalis | SP00162 | Ligilactobacillus | salivarius | 1 |
| SP00155 | Enterococcus | faecalis | SP00163 | Staphylococcus | aureus | 1 |
| SP00155 | Enterococcus | faecalis | SP00251 | Streptococcus | constellatus | 1 |
| SP00155 | Enterococcus | faecalis | SP00280 | Staphylococcus | lugdunensis | 1 |
| SP00158 | Lactobacillus | johnsonii | SP00181 | Levilactobacillus | brevis | 1 |
| SP00158 | Lactobacillus | johnsonii | SP00237 | Lentilactobacillus | buchneri | 1 |
| SP00178 | Rothia | mucilaginosa | SP00214 | Micrococcus | luteus | 1 |
| SP00178 | Rothia | mucilaginosa | SP00219 | Kytococcus | sedentarius | 1 |
| SP00178 | Rothia | mucilaginosa | SP00224 | Sanguibacter | keddieii | 1 |
| SP00178 | Rothia | mucilaginosa | SP00301 | Kocuria | rhizophila | 1 |
| SP00179 | Bacillus | subtilis | SP00297 | Staphylococcus | cohnii | 1 |
| SP00181 | Levilactobacillus | brevis | SP00189 | Lactobacillus | acidophilus | 1 |
| SP00181 | Levilactobacillus | brevis | SP00195 | Lactobacillus | gasseri | 1 |
| SP00181 | Levilactobacillus | brevis | SP00235 | Lactobacillus | amylovorus | 1 |
| SP00181 | Levilactobacillus | brevis | SP00297 | Staphylococcus | cohnii | 1 |
| SP00189 | Lactobacillus | acidophilus | SP00237 | Lentilactobacillus | buchneri | 1 |
| SP00192 | Bifidobacterium | breve | SP00226 | Gardnerella | vaginalis | 1 |
| SP00195 | Lactobacillus | gasseri | SP00237 | Lentilactobacillus | buchneri | 1 |
| SP00211 | Acidovorax | ebreus | SP00268 | Ottowia | sp. oral taxon 894 | 1 |
| SP00213 | Corynebacterium | kroppenstedtii | SP00310 | Dietzia | sp. oral taxon 368 | 1 |
| SP00214 | Micrococcus | luteus | SP00232 | Rothia | dentocariosa | 1 |
| SP00216 | Variovorax | paradoxus | SP00268 | Ottowia | sp. oral taxon 894 | 1 |
| SP00219 | Kytococcus | sedentarius | SP00224 | Sanguibacter | keddieii | 1 |
| SP00219 | Kytococcus | sedentarius | SP00232 | Rothia | dentocariosa | 1 |
| SP00223 | Bifidobacterium | dentium | SP00226 | Gardnerella | vaginalis | 1 |
| SP00224 | Sanguibacter | keddieii | SP00232 | Rothia | dentocariosa | 1 |
| SP00230 | Prevotella | melaninogenica | SP00276 | Prevotella | enoeca | 1 |
| SP00232 | Rothia | dentocariosa | SP00265 | Arsenicicoccus | sp. oral taxon 190 | 1 |
| SP00232 | Rothia | dentocariosa | SP00301 | Kocuria | rhizophila | 1 |
| SP00235 | Lactobacillus | amylovorus | SP00237 | Lentilactobacillus | buchneri | 1 |
| SP00237 | Lentilactobacillus | buchneri | SP00297 | Staphylococcus | cohnii | 1 |
| SP00239 | Pseudopropionibacterium | propionicum | SP00254 | Cutibacterium | avidum | 1 |
| SP00242 | Streptococcus | salivarius | SP00288 | Streptococcus | sp. oral taxon 431 | 1 |
| SP00260 | Acinetobacter | johnsonii | SP00287 | Moraxella | osloensis | 1 |
| SP00270 | Lawsonella | clevelandensis | SP00310 | Dietzia | sp. oral taxon 368 | 1 |
| SP00288 | Streptococcus | sp. oral taxon 431 | SP00320 | Streptococcus | thermophilus | 1 |
| ***TOTAL*** | | | | | | 4450 |

The Table details all the pairs of bacterial species which had amplicon similarity/identity ≥97% using the bacterial-specific and the bacterial and archaeal primer pairs analyzed in the present study. Freq.= frequency, number times that a pair of species had amplicon similarity/identity ≥97% in the different primer pairs; ID= species identifier.

Appendix Table 13. Genus of the pairs of bacterial species with amplicon similarity ≥97%.

| **Pair of genus** | **Frequency** |
| --- | --- |
| Streptococcus\|Streptococcus | 1310 |
| Staphylococcus\|Staphylococcus | 1198 |
| Corynebacterium\|Corynebacterium | 121 |
| Actinomyces\|Actinomyces | 120 |
| Pseudomonas\|Pseudomonas | 93 |
| Klebsiella\|Klebsiella | 82 |
| Capnocytophaga\|Capnocytophaga | 75 |
| Neisseria\|Neisseria | 70 |
| Lactobacillus\|Lactobacillus | 69 |
| Acinetobacter\|Acinetobacter | 65 |
| Bifidobacterium\|Bifidobacterium | 53 |
| Haemophilus\|Haemophilus | 40 |
| Cronobacter\|Cronobacter | 28 |
| Fusobacterium\|Fusobacterium | 28 |
| Lacticaseibacillus\|Lacticaseibacillus | 28 |
| Mycobacterium\|Mycobacterium | 27 |
| Selenomonas\|Selenomonas | 27 |
| Treponema\|Treponema | 27 |
| Olsenella\|Olsenella | 25 |
| Schaalia\|Schaalia | 22 |
| Campylobacter\|Campylobacter | 19 |
| Kocuria\|Kocuria | 18 |
| Rothia\|Rothia | 18 |
| Prevotella\|Prevotella | 13 |
| Leptotrichia\|Leptotrichia | 12 |
| Mycoplasma\|Mycoplasma | 11 |
| Tannerella\|Tannerella | 8 |
| Cutibacterium\|Cutibacterium | 7 |
| Limosilactobacillus\|Limosilactobacillus | 7 |
| Bacteroides\|Bacteroides | 6 |
| Aggregatibacter\|Aggregatibacter | 5 |
| Agrobacterium\|Agrobacterium | 4 |
| Bacillus\|Bacillus | 3 |
| Moraxella\|Moraxella | 2 |
| ***TOTAL*** | 3641 |
| **Pair of different genus** | **Frequency** |
| Cronobacter\|Klebsiella | 99 |
| Klebsiella\|Serratia | 67 |
| Escherichia\|Klebsiella | 39 |
| Cronobacter\|Escherichia | 36 |
| Aggregatibacter\|Haemophilus | 28 |
| Achromobacter\|Bordetella | 25 |
| Cutibacterium\|Propionibacterium | 24 |
| Cupriavidus\|Ralstonia | 23 |
| Klebsiella\|Yersinia | 23 |
| Listeria\|Staphylococcus | 23 |
| Bacillus\|Staphylococcus | 22 |
| Cronobacter\|Serratia | 22 |
| Actinomyces\|Schaalia | 21 |
| Mycobacterium\|Mycolicibacterium | 21 |
| Escherichia\|Serratia | 20 |
| Acidovorax\|Comamonas | 18 |
| Serratia\|Yersinia | 16 |
| Cronobacter\|Yersinia | 14 |
| Lentilactobacillus\|Levilactobacillus | 14 |
| Janibacter\|Kytococcus | 13 |
| Paracoccus\|Rhodobacter | 13 |
| Lacticaseibacillus\|Lactiplantibacillus | 12 |
| Lacticaseibacillus\|Lentilactobacillus | 11 |
| Lacticaseibacillus\|Levilactobacillus | 11 |
| Lacticaseibacillus\|Limosilactobacillus | 9 |
| Lentilactobacillus\|Limosilactobacillus | 9 |
| Acidovorax\|Variovorax | 8 |
| Agrobacterium\|Brucella | 8 |
| Comamonas\|Variovorax | 7 |
| Escherichia\|Yersinia | 7 |
| Kocuria\|Rothia | 7 |
| Lactiplantibacillus\|Levilactobacillus | 7 |
| Delftia\|Variovorax | 6 |
| Haemophilus\|Klebsiella | 6 |
| Kocuria\|Micrococcus | 6 |
| Lactiplantibacillus\|Lentilactobacillus | 6 |
| Levilactobacillus\|Limosilactobacillus | 5 |
| Agrobacterium\|Mesorhizobium | 4 |
| Enterococcus\|Listeria | 4 |
| Enterococcus\|Staphylococcus | 4 |
| Janibacter\|Kocuria | 4 |
| Janibacter\|Sanguibacter | 4 |
| Lacticaseibacillus\|Ligilactobacillus | 4 |
| Lactobacillus\|Lentilactobacillus | 4 |
| Lactobacillus\|Levilactobacillus | 4 |
| Arsenicicoccus\|Janibacter | 3 |
| Arsenicicoccus\|Sanguibacter | 3 |
| Bifidobacterium\|Gardnerella | 3 |
| Brucella\|Mesorhizobium | 3 |
| Comamonas\|Delftia | 3 |
| Corynebacterium\|Dietzia | 3 |
| Micrococcus\|Sanguibacter | 3 |
| Acidovorax\|Delftia | 2 |
| Arsenicicoccus\|Kytococcus | 2 |
| Bacillus\|Listeria | 2 |
| Burkholderia\|Cupriavidus | 2 |
| Burkholderia\|Ralstonia | 2 |
| Delftia\|Ottowia | 2 |
| Enterococcus\|Lentilactobacillus | 2 |
| Enterococcus\|Levilactobacillus | 2 |
| Escherichia\|Haemophilus | 2 |
| Haemophilus\|Serratia | 2 |
| Janibacter\|Lawsonella | 2 |
| Janibacter\|Micrococcus | 2 |
| Kocuria\|Sanguibacter | 2 |
| Kytococcus\|Micrococcus | 2 |
| Kytococcus\|Rothia | 2 |
| Lawsonella\|Micrococcus | 2 |
| Lawsonella\|Sanguibacter | 2 |
| Micrococcus\|Rothia | 2 |
| Rothia\|Sanguibacter | 2 |
| Acidovorax\|Ottowia | 1 |
| Acinetobacter\|Moraxella | 1 |
| Arsenicicoccus\|Rothia | 1 |
| Cutibacterium\|Pseudopropionibacterium | 1 |
| Dietzia\|Lawsonella | 1 |
| Enterococcus\|Ligilactobacillus | 1 |
| Enterococcus\|Streptococcus | 1 |
| Kytococcus\|Sanguibacter | 1 |
| Lentilactobacillus\|Staphylococcus | 1 |
| Levilactobacillus\|Staphylococcus | 1 |
| Ottowia\|Variovorax | 1 |
| Proteus\|Yersinia | 1 |
| ***TOTAL*** | 809 |

Appendix Table 14. Families of the pairs of bacterial species with amplicon similarity ≥97%.

| **Pair of families** | **Frequency** |
| --- | --- |
| Streptococcaceae\|Streptococcaceae | 1310 |
| Staphylococcaceae\|Staphylococcaceae | 1198 |
| Enterobacteriaceae\|Enterobacteriaceae | 284 |
| Lactobacillaceae\|Lactobacillaceae | 200 |
| Actinomycetaceae\|Actinomycetaceae | 163 |
| Corynebacteriaceae\|Corynebacteriaceae | 121 |
| Pseudomonadaceae\|Pseudomonadaceae | 93 |
| Flavobacteriaceae\|Flavobacteriaceae | 75 |
| Pasteurellaceae\|Pasteurellaceae | 73 |
| Neisseriaceae\|Neisseriaceae | 70 |
| Moraxellaceae\|Moraxellaceae | 68 |
| Bifidobacteriaceae\|Bifidobacteriaceae | 56 |
| Micrococcaceae\|Micrococcaceae | 51 |
| Comamonadaceae\|Comamonadaceae | 48 |
| Mycobacteriaceae\|Mycobacteriaceae | 48 |
| Propionibacteriaceae\|Propionibacteriaceae | 32 |
| Fusobacteriaceae\|Fusobacteriaceae | 28 |
| Burkholderiaceae\|Burkholderiaceae | 27 |
| Selenomonadaceae\|Selenomonadaceae | 27 |
| Spirochaetaceae\|Spirochaetaceae | 27 |
| Alcaligenaceae\|Alcaligenaceae | 25 |
| Atopobiaceae\|Atopobiaceae | 25 |
| Campylobacteraceae\|Campylobacteraceae | 19 |
| Yersiniaceae\|Yersiniaceae | 16 |
| Prevotellaceae\|Prevotellaceae | 13 |
| Rhodobacteraceae\|Rhodobacteraceae | 13 |
| Leptotrichiaceae\|Leptotrichiaceae | 12 |
| Mycoplasmataceae\|Mycoplasmataceae | 11 |
| Tannerellaceae\|Tannerellaceae | 8 |
| Bacteroidaceae\|Bacteroidaceae | 6 |
| Rhizobiaceae\|Rhizobiaceae | 4 |
| Bacillaceae\|Bacillaceae | 3 |
| Intrasporangiaceae\|Intrasporangiaceae | 3 |
| ***TOTAL*** | 4157 |
| **Pair of different families** | **Frequency** |
| Enterobacteriaceae\|Yersiniaceae | 153 |
| Listeriaceae\|Staphylococcaceae | 23 |
| Bacillaceae\|Staphylococcaceae | 22 |
| Intrasporangiaceae\|Kytococcaceae | 15 |
| Brucellaceae\|Rhizobiaceae | 8 |
| Enterobacteriaceae\|Pasteurellaceae | 8 |
| Intrasporangiaceae\|Micrococcaceae | 7 |
| Intrasporangiaceae\|Sanguibacteraceae | 7 |
| Micrococcaceae\|Sanguibacteraceae | 7 |
| Enterococcaceae\|Lactobacillaceae | 5 |
| Enterococcaceae\|Listeriaceae | 4 |
| Enterococcaceae\|Staphylococcaceae | 4 |
| Kytococcaceae\|Micrococcaceae | 4 |
| Phyllobacteriaceae\|Rhizobiaceae | 4 |
| Brucellaceae\|Phyllobacteriaceae | 3 |
| Corynebacteriaceae\|Dietziaceae | 3 |
| Bacillaceae\|Listeriaceae | 2 |
| Intrasporangiaceae\|Lawsonellaceae | 2 |
| Lactobacillaceae\|Staphylococcaceae | 2 |
| Lawsonellaceae\|Micrococcaceae | 2 |
| Lawsonellaceae\|Sanguibacteraceae | 2 |
| Pasteurellaceae\|Yersiniaceae | 2 |
| Dietziaceae\|Lawsonellaceae | 1 |
| Enterococcaceae\|Streptococcaceae | 1 |
| Kytococcaceae\|Sanguibacteraceae | 1 |
| Morganellaceae\|Yersiniaceae | 1 |
| ***TOTAL*** | 293 |

Appendix Table 15. Orders of the pairs of bacterial species with amplicon similarity ≥97%.

| **Pair of orders** | **Frequency** |
| --- | --- |
| Lactobacillales\|Lactobacillales | 1516 |
| Bacillales\|Bacillales | 1248 |
| Enterobacterales\|Enterobacterales | 454 |
| Corynebacteriales\|Corynebacteriales | 173 |
| Actinomycetales\|Actinomycetales | 163 |
| Pseudomonadales\|Pseudomonadales | 161 |
| Burkholderiales\|Burkholderiales | 100 |
| Micrococcales\|Micrococcales | 95 |
| Flavobacteriales\|Flavobacteriales | 75 |
| Pasteurellales\|Pasteurellales | 73 |
| Neisseriales\|Neisseriales | 70 |
| Bifidobacteriales\|Bifidobacteriales | 56 |
| Fusobacteriales\|Fusobacteriales | 40 |
| Propionibacteriales\|Propionibacteriales | 32 |
| Bacteroidales\|Bacteroidales | 27 |
| Selenomonadales\|Selenomonadales | 27 |
| Spirochaetales\|Spirochaetales | 27 |
| Coriobacteriales\|Coriobacteriales | 25 |
| Campylobacterales\|Campylobacterales | 19 |
| Hyphomicrobiales\|Hyphomicrobiales | 19 |
| Rhodobacterales\|Rhodobacterales | 13 |
| Mycoplasmatales\|Mycoplasmatales | 11 |
| ***TOTAL*** | 4424 |
| **Pair of different orders** | **Frequency** |
| Bacillales\|Lactobacillales | 10 |
| Enterobacterales\|Pasteurellales | 10 |
| Corynebacteriales\|Micrococcales | 6 |
| ***TOTAL*** | 26 |

Appendix Table 16. Classes of the pairs of bacterial species with amplicon similarity ≥97%.

| **Pair of classes** | **Frequency** |
| --- | --- |
| Bacilli\|Bacilli | 2774 |
| Gammaproteobacteria\|Gammaproteobacteria | 698 |
| Actinomycetia\|Actinomycetia | 525 |
| Betaproteobacteria\|Betaproteobacteria | 170 |
| Flavobacteriia\|Flavobacteriia | 75 |
| Fusobacteriia\|Fusobacteriia | 40 |
| Alphaproteobacteria\|Alphaproteobacteria | 32 |
| Bacteroidia\|Bacteroidia | 27 |
| Negativicutes\|Negativicutes | 27 |
| Spirochaetia\|Spirochaetia | 27 |
| Coriobacteriia\|Coriobacteriia | 25 |
| Epsilonproteobacteria\|Epsilonproteobacteria | 19 |
| Mollicutes\|Mollicutes | 11 |
| ***TOTAL*** | 4450 |

Appendix Table 17. Oral archaea species with amplicon similarity values ≥97% with at least one different taxon.

| **ID** | **Phylum** | **Class** | **Order** | **Family** | **Genus** | **Species** |
| --- | --- | --- | --- | --- | --- | --- |
| SP00001 | Crenarchaeota | Thermoprotei | Desulfurococcales | Desulfurococcaceae | Ignisphaera | aggregans |
| SP00002 | Euryarchaeota | Methanomicrobia | Methanosarcinales | Methanosarcinaceae | Methanolobus | psychrophilus |
| SP00003 | Crenarchaeota | Thermoprotei | Thermoproteales | Thermoproteaceae | Pyrobaculum | oguniense |
| SP00004 | Crenarchaeota | Thermoprotei | Desulfurococcales | Desulfurococcaceae | Aeropyrum | pernix |
| SP00005 | Euryarchaeota | Methanococci | Methanococcales | Methanocaldococcaceae | Methanocaldococcus | jannaschii |
| SP00006 | Euryarchaeota | Thermococci | Thermococcales | Thermococcaceae | Pyrococcus | horikoshii |
| SP00007 | Euryarchaeota | Methanococci | Methanococcales | Methanococcaceae | Methanococcus | maripaludis |
| SP00008 | Euryarchaeota | Halobacteria | Halobacteriales | Halobacteriaceae | Halobacterium | salinarum |
| SP00010 | Crenarchaeota | Thermoprotei | Thermoproteales | Thermoproteaceae | Pyrobaculum | aerophilum |
| SP00012 | Euryarchaeota | Methanomicrobia | Methanosarcinales | Methanosarcinaceae | Methanosarcina | acetivorans |
| SP00013 | Euryarchaeota | Methanomicrobia | Methanosarcinales | Methanosarcinaceae | Methanosarcina | mazei |
| SP00015 | Euryarchaeota | Halobacteria | Halobacteriales | Haloarculaceae | Haloarcula | marismortui |
| SP00017 | Euryarchaeota | Methanomicrobia | Methanosarcinales | Methanosarcinaceae | Methanosarcina | barkeri |
| SP00018 | Euryarchaeota | Halobacteria | Halobacteriales | Haloarculaceae | Natronomonas | pharaonis |
| SP00020 | Euryarchaeota | Methanomicrobia | Methanosarcinales | Methanosarcinaceae | Methanococcoides | burtonii |
| SP00022 | Euryarchaeota | Methanomicrobia | Methanosarcinales | Methanotrichaceae | Methanothrix | thermoacetophila |
| SP00024 | Crenarchaeota | Thermoprotei | Desulfurococcales | Pyrodictiaceae | Hyperthermus | butylicus |
| SP00026 | Crenarchaeota | Thermoprotei | Desulfurococcales | Desulfurococcaceae | Staphylothermus | marinus |
| SP00027 | Euryarchaeota | Methanomicrobia | Methanomicrobiales | Methanomicrobiaceae | Methanoculleus | marisnigri |
| SP00028 | Crenarchaeota | Thermoprotei | Thermoproteales | Thermoproteaceae | Pyrobaculum | arsenaticum |
| SP00030 | Euryarchaeota | Methanobacteria | Methanobacteriales | Methanobacteriaceae | Methanobrevibacter | smithii |
| SP00031 | Euryarchaeota | Methanococci | Methanococcales | Methanococcaceae | Methanococcus | vannielii |
| SP00032 | Euryarchaeota | Methanococci | Methanococcales | Methanococcaceae | Methanococcus | aeolicus |
| SP00033 | Euryarchaeota | Methanomicrobia | Methanomicrobiales | Methanoregulaceae | Methanoregula | boonei |
| SP00034 | Crenarchaeota | Thermoprotei | Desulfurococcales | Desulfurococcaceae | Ignicoccus | hospitalis |
| SP00036 | Euryarchaeota | Thermococci | Thermococcales | Thermococcaceae | Thermococcus | onnurineus |
| SP00037 | Crenarchaeota | Thermoprotei | Desulfurococcales | Desulfurococcaceae | Desulfurococcus | amylolyticus |
| SP00038 | Euryarchaeota | Methanomicrobia | Methanomicrobiales | Methanoregulaceae | Methanosphaerula | palustris |
| SP00041 | Euryarchaeota | Thermococci | Thermococcales | Thermococcaceae | Thermococcus | gammatolerans |
| SP00042 | Euryarchaeota | Thermococci | Thermococcales | Thermococcaceae | Thermococcus | sibiricus |
| SP00043 | Euryarchaeota | Methanococci | Methanococcales | Methanocaldococcaceae | Methanocaldococcus | fervens |
| SP00044 | Euryarchaeota | Halobacteria | Halobacteriales | Haloarculaceae | Halorhabdus | utahensis |
| SP00045 | Euryarchaeota | Halobacteria | Halobacteriales | Haloarculaceae | Halomicrobium | mukohataei |
| SP00046 | Euryarchaeota | Methanococci | Methanococcales | Methanocaldococcaceae | Methanocaldococcus | vulcanius |
| SP00048 | Euryarchaeota | Archaeoglobi | Archaeoglobales | Archaeoglobaceae | Archaeoglobus | profundus |
| SP00049 | Euryarchaeota | Halobacteria | Natrialbales | Natrialbaceae | Haloterrigena | turkmenica |
| SP00050 | Euryarchaeota | Methanobacteria | Methanobacteriales | Methanobacteriaceae | Methanobrevibacter | ruminantium |
| SP00051 | Euryarchaeota | Archaeoglobi | Archaeoglobales | Archaeoglobaceae | Ferroglobus | placidus |
| SP00052 | Euryarchaeota | Methanococci | Methanococcales | Methanocaldococcaceae | Methanocaldococcus | sp. FS406-22 |
| SP00053 | Euryarchaeota | Halobacteria | Natrialbales | Natrialbaceae | Natrialba | magadii |
| SP00055 | Euryarchaeota | Methanomicrobia | Methanosarcinales | Methanosarcinaceae | Methanohalophilus | mahii |
| SP00056 | Euryarchaeota | Methanococci | Methanococcales | Methanocaldococcaceae | Methanocaldococcus | infernus |
| SP00057 | Crenarchaeota | Thermoprotei | Desulfurococcales | Desulfurococcaceae | Staphylothermus | hellenicus |
| SP00058 | Euryarchaeota | Methanococci | Methanococcales | Methanococcaceae | Methanococcus | voltae |
| SP00059 | Euryarchaeota | Methanomicrobia | Methanosarcinales | Methanosarcinaceae | Methanohalobium | evestigatum |
| SP00061 | Crenarchaeota | Thermoprotei | Acidilobales | Acidilobaceae | Acidilobus | saccharovorans |
| SP00062 | Euryarchaeota | Methanobacteria | Methanobacteriales | Methanobacteriaceae | Methanothermobacter | marburgensis |
| SP00064 | Crenarchaeota | Thermoprotei | Thermoproteales | Thermoproteaceae | Vulcanisaeta | distributa |
| SP00067 | Euryarchaeota | Thermococci | Thermococcales | Thermococcaceae | Thermococcus | barophilus |
| SP00068 | Crenarchaeota | Thermoprotei | Desulfurococcales | Desulfurococcaceae | Desulfurococcus | mucosus |
| SP00069 | Euryarchaeota | Methanobacteria | Methanobacteriales | Methanobacteriaceae | Methanobacterium | lacus |
| SP00070 | Crenarchaeota | Thermoprotei | Thermoproteales | Thermoproteaceae | Thermoproteus | uzoniensis |
| SP00071 | Euryarchaeota | Archaeoglobi | Archaeoglobales | Archaeoglobaceae | Archaeoglobus | veneficus |
| SP00072 | Euryarchaeota | Methanomicrobia | Methanosarcinales | Methanotrichaceae | Methanothrix | soehngenii |
| SP00074 | Euryarchaeota | Thermococci | Thermococcales | Thermococcaceae | Pyrococcus | sp. NA2 |
| SP00075 | Euryarchaeota | Methanococci | Methanococcales | Methanocaldococcaceae | Methanotorris | igneus |
| SP00076 | Euryarchaeota | Methanobacteria | Methanobacteriales | Methanobacteriaceae | Methanobacterium | paludis |
| SP00077 | Euryarchaeota | Methanococci | Methanococcales | Methanococcaceae | Methanothermococcus | okinawensis |
| SP00078 | Euryarchaeota | Halobacteria | Natrialbales | Natrialbaceae | Halopiger | xanaduensis |
| SP00079 | Euryarchaeota | Methanomicrobia | Methanosarcinales | Methanosarcinaceae | Methanosalsum | zhilinae |
| SP00080 | Euryarchaeota | Thermococci | Thermococcales | Thermococcaceae | Pyrococcus | yayanosii |
| SP00081 | Euryarchaeota | Thermococci | Thermococcales | Thermococcaceae | Thermococcus | sp. 4557 |
| SP00082 | Crenarchaeota | Thermoprotei | Desulfurococcales | Pyrodictiaceae | Pyrolobus | fumarii |
| SP00083 | Euryarchaeota | Halobacteria | Halobacteriales | Haloarculaceae | Haloarcula | hispanica |
| SP00084 | Crenarchaeota | Thermoprotei | Thermoproteales | Thermoproteaceae | Thermoproteus | tenax |
| SP00086 | Crenarchaeota | Thermoprotei | Thermoproteales | Thermoproteaceae | Pyrobaculum | ferrireducens |
| SP00089 | Euryarchaeota | Thermococci | Thermococcales | Thermococcaceae | Pyrococcus | sp. ST04 |
| SP00090 | Crenarchaeota | Thermoprotei | Desulfurococcales | Desulfurococcaceae | Thermogladius | calderae |
| SP00091 | Euryarchaeota | Thermococci | Thermococcales | Thermococcaceae | Thermococcus | cleftensis |
| SP00092 | Euryarchaeota | Halobacteria | Natrialbales | Natrialbaceae | Natrinema | sp. J7-2 |
| SP00093 | Euryarchaeota | Methanomicrobia | Methanomicrobiales | Methanomicrobiaceae | Methanoculleus | bourgensis |
| SP00094 | Thaumarchaeota | Nitrososphaeria | Nitrososphaerales | Nitrososphaeraceae | Nitrososphaera | gargensis (C.) |
| SP00096 | Euryarchaeota | Halobacteria | Natrialbales | Natrialbaceae | Natronobacterium | gregoryi |
| SP00097 | Euryarchaeota | Methanomicrobia | Methanomicrobiales | Methanoregulaceae | Methanoregula | formicica |
| SP00098 | Euryarchaeota | Halobacteria | Natrialbales | Natrialbaceae | Natrinema | pellirubrum |
| SP00099 | Euryarchaeota | Halobacteria | Natrialbales | Natrialbaceae | Halovivax | ruber |
| SP00100 | Euryarchaeota | Methanomicrobia | Methanosarcinales | Methanosarcinaceae | Methanomethylovorans | hollandica |
| SP00101 | Euryarchaeota | Halobacteria | Natrialbales | Natrialbaceae | Natronococcus | occultus |
| SP00102 | Euryarchaeota | Halobacteria | Halobacteriales | Haloarculaceae | Natronomonas | moolapensis |
| SP00104 | Euryarchaeota | Archaeoglobi | Archaeoglobales | Archaeoglobaceae | Archaeoglobus | sulfaticallidus |
| SP00106 | Euryarchaeota | Methanobacteria | Methanobacteriales | Methanobacteriaceae | Methanobrevibacter | sp. AbM4 |
| SP00108 | Euryarchaeota | Halobacteria | Halobacteriales | Haloarculaceae | Halorhabdus | tiamatea |
| SP00109 | Euryarchaeota | Thermococci | Thermococcales | Thermococcaceae | Thermococcus | litoralis |
| SP00110 | Crenarchaeota | Thermoprotei | Desulfurococcales | Desulfurococcaceae | Aeropyrum | camini |
| SP00111 | Euryarchaeota | Thermococci | Thermococcales | Thermococcaceae | Palaeococcus | pacificus |
| SP00112 | Euryarchaeota | Methanobacteria | Methanobacteriales | Methanobacteriaceae | Methanobacterium | formicicum |
| SP00113 | Euryarchaeota | Thermococci | Thermococcales | Thermococcaceae | Thermococcus | paralvinellae |
| SP00114 | Euryarchaeota | Halobacteria | Natrialbales | Natrialbaceae | Halostagnicola | larsenii |
| SP00115 | Euryarchaeota | Halobacteria | Halobacteriales | Halobacteriaceae | Halobacterium | sp. DL1 |
| SP00116 | Thaumarchaeota | Nitrososphaeria | Nitrososphaerales | Nitrososphaeraceae | Nitrososphaera | evergladensis (C.) |
| SP00117 | Thaumarchaeota | Nitrososphaeria | Nitrososphaerales | Nitrososphaeraceae | Nitrososphaera | viennensis |
| SP00118 | Euryarchaeota | Thermococci | Thermococcales | Thermococcaceae | Thermococcus | eurythermalis |
| SP00119 | Euryarchaeota | Methanococci | Methanococcales | Methanocaldococcaceae | Methanocaldococcus | bathoardescens |
| SP00120 | Euryarchaeota | Methanomicrobia | Methanosarcinales | Methanosarcinaceae | Methanosarcina | thermophila |
| SP00121 | Euryarchaeota | Methanomicrobia | Methanosarcinales | Methanosarcinaceae | Methanosarcina | sp. WWM596 |
| SP00122 | Euryarchaeota | Methanomicrobia | Methanosarcinales | Methanosarcinaceae | Methanosarcina | sp. WH1 |
| SP00123 | Euryarchaeota | Methanomicrobia | Methanosarcinales | Methanosarcinaceae | Methanosarcina | sp. MTP4 |
| SP00124 | Euryarchaeota | Methanomicrobia | Methanosarcinales | Methanosarcinaceae | Methanosarcina | siciliae |
| SP00125 | Euryarchaeota | Methanomicrobia | Methanosarcinales | Methanosarcinaceae | Methanosarcina | lacustris |
| SP00126 | Euryarchaeota | Methanomicrobia | Methanosarcinales | Methanosarcinaceae | Methanosarcina | horonobensis |
| SP00127 | Euryarchaeota | Methanomicrobia | Methanosarcinales | Methanosarcinaceae | Methanococcoides | methylutens |
| SP00128 | Euryarchaeota | Methanomicrobia | Methanosarcinales | Methanosarcinaceae | Methanosarcina | vacuolata |
| SP00129 | Euryarchaeota | Methanomicrobia | Methanosarcinales | Methanosarcinaceae | Methanosarcina | sp. Kolksee |
| SP00131 | Euryarchaeota | Halobacteria | Halobacteriales | Haloarculaceae | Haloarcula | sp. CBA1115 |
| SP00132 | Euryarchaeota | Methanobacteria | Methanobacteriales | Methanobacteriaceae | Methanobrevibacter | millerae |
| SP00133 | Euryarchaeota | Halobacteria | Halobacteriales | Haloarculaceae | Halomicrobium | sp. ZPS1 |
| SP00134 | Euryarchaeota | Methanobacteria | Methanobacteriales | Methanobacteriaceae | Methanosphaera | stadtmanae |
| SP00135 | Euryarchaeota | Methanobacteria | Methanobacteriales | Methanobacteriaceae | Methanobacterium | congolense |

The Table details the archaeal species which had amplicon similarity/identity values ≥97% with at least one different taxon using the archaeal-specific and the bacterial and archaeal primer pairs analyzed in the present study. C.= candidatus; ID= species identifier.

Appendix Table 18. Oral archaea species with no amplicon similarity values ≥97% with different taxa.

| **ID** | **Phylum** | **Class** | **Order** | **Family** | **Genus** | **Species** |
| --- | --- | --- | --- | --- | --- | --- |
| SP00009 | C.Thermoplasmatota | Thermoplasmata | Thermoplasmatales | Thermoplasmataceae | Thermoplasma | volcanium |
| SP00011 | Euryarchaeota | Methanopyri | Methanopyrales | Methanopyraceae | Methanopyrus | kandleri |
| SP00014 | C.Thermoplasmatota | Thermoplasmata | Thermoplasmatales | Picrophilaceae | Picrophilus | torridus |
| SP00016 | Crenarchaeota | Thermoprotei | Sulfolobales | Sulfolobaceae | Sulfolobus | acidocaldarius |
| SP00019 | Euryarchaeota | Methanomicrobia | Methanomicrobiales | Methanospirillaceae | Methanospirillum | hungatei |
| SP00021 | Euryarchaeota | Halobacteria | Haloferacales | Haloferacaceae | Haloquadratum | walsbyi |
| SP00023 | Crenarchaeota | Thermoprotei | Thermoproteales | Thermofilaceae | Thermofilum | pendens |
| SP00025 | Euryarchaeota | Methanomicrobia | Methanomicrobiales | Methanocorpusculaceae | Methanocorpusculum | labreanum |
| SP00029 | Euryarchaeota | Methanomicrobia | Methanocellales | Methanocellaceae | Methanocella | arvoryzae |
| SP00035 | Crenarchaeota | Thermoprotei | Sulfolobales | Sulfolobaceae | Acidianus | hospitalis |
| SP00039 | Euryarchaeota | Halobacteria | Haloferacales | Halorubraceae | Halorubrum | lacusprofundi |
| SP00040 | Crenarchaeota | Thermoprotei | Sulfolobales | Sulfolobaceae | Sulfolobus | islandicus |
| SP00047 | Euryarchaeota | Methanomicrobia | Methanocellales | Methanocellaceae | Methanocella | paludicola |
| SP00054 | Euryarchaeota | Halobacteria | Haloferacales | Haloferacaceae | Haloferax | volcanii |
| SP00060 | Euryarchaeota | Halobacteria | Halobacteriales | Halobacteriaceae | Halalkalicoccus | jeotgali |
| SP00063 | Euryarchaeota | Methanomicrobia | Methanomicrobiales | Methanomicrobiaceae | Methanolacinia | petrolearia |
| SP00065 | Euryarchaeota | Methanobacteria | Methanobacteriales | Methanothermaceae | Methanothermus | fervidus |
| SP00066 | Euryarchaeota | Halobacteria | Haloferacales | Haloferacaceae | Halogeometricum | borinquense |
| SP00073 | Crenarchaeota | Thermoprotei | Sulfolobales | Sulfolobaceae | Metallosphaera | cuprina |
| SP00085 | Euryarchaeota | Methanomicrobia | Methanosarcinales | Methanotrichaceae | Methanothrix | harundinacea |
| SP00087 | Euryarchaeota | Methanomicrobia | Methanocellales | Methanocellaceae | Methanocella | conradii |
| SP00088 | Crenarchaeota | Thermoprotei | Fervidicoccales | Fervidicoccaceae | Fervidicoccus | fontis |
| SP00095 | Crenarchaeota | Thermoprotei | Acidilobales | Caldisphaeraceae | Caldisphaera | lagunensis |
| SP00103 | C.Thermoplasmatota | Thermoplasmata | Methanomassiliicoccales | C.Methanomethylophilaceae | C.Methanomethylophilus | alvus |
| SP00105 | C.Thermoplasmatota | Thermoplasmata | Methanomassiliicoccales | Methanomassiliicoccaceae | Methanomassiliicoccus | intestinalis(c.) |
| SP00107 | C.Thermoplasmatota | Thermoplasmata | Thermoplasmatales | Ferroplasmaceae | Ferroplasma | acidarmanus |
| SP00130 | C.Thermoplasmatota | Thermoplasmata | Methanomassiliicoccales | Methanomassiliicoccaceae | C. Methanoplasma | termitum |

The Table illustrates the archaeal species which had no amplicon similarity/identity values ≥97% with different taxa using the archaeal-specific and the bacterial and archaeal primer pairs analyzed in the present study. C.= candidatus; ID= species identifier.

Appendix Table 19. Pairs of oral archaeal species with amplicon similarity ≥97% using the analyzed primer pairs.

| **ID** | **Genus** | **Species** | **ID** | **Genus** | **Species** | **Freq** |
| --- | --- | --- | --- | --- | --- | --- |
| SP00005 | Methanocaldococcus | jannaschii | SP00043 | Methanocaldococcus | Fervens | 20 |
| SP00005 | Methanocaldococcus | jannaschii | SP00052 | Methanocaldococcus | sp. FS406-22 | 20 |
| SP00005 | Methanocaldococcus | jannaschii | SP00119 | Methanocaldococcus | Bathoardescens | 20 |
| SP00030 | Methanobrevibacter | smithii | SP00132 | Methanobrevibacter | Millerae | 20 |
| SP00043 | Methanocaldococcus | fervens | SP00052 | Methanocaldococcus | sp. FS406-22 | 20 |
| SP00043 | Methanocaldococcus | fervens | SP00119 | Methanocaldococcus | Bathoardescens | 20 |
| SP00052 | Methanocaldococcus | sp. FS406-22 | SP00119 | Methanocaldococcus | Bathoardescens | 20 |
| SP00006 | Pyrococcus | horikoshii | SP00074 | Pyrococcus | sp. NA2 | 19 |
| SP00006 | Pyrococcus | horikoshii | SP00080 | Pyrococcus | Yayanosii | 19 |
| SP00006 | Pyrococcus | horikoshii | SP00089 | Pyrococcus | sp. ST04 | 19 |
| SP00012 | Methanosarcina | acetivorans | SP00013 | Methanosarcina | Mazei | 19 |
| SP00012 | Methanosarcina | acetivorans | SP00017 | Methanosarcina | Barkeri | 19 |
| SP00012 | Methanosarcina | acetivorans | SP00120 | Methanosarcina | Thermophila | 19 |
| SP00012 | Methanosarcina | acetivorans | SP00121 | Methanosarcina | sp. WWM596 | 19 |
| SP00012 | Methanosarcina | acetivorans | SP00122 | Methanosarcina | sp. WH1 | 19 |
| SP00012 | Methanosarcina | acetivorans | SP00124 | Methanosarcina | Siciliae | 19 |
| SP00012 | Methanosarcina | acetivorans | SP00126 | Methanosarcina | Horonobensis | 19 |
| SP00012 | Methanosarcina | acetivorans | SP00128 | Methanosarcina | Vacuolata | 19 |
| SP00012 | Methanosarcina | acetivorans | SP00129 | Methanosarcina | sp. Kolksee | 19 |
| SP00013 | Methanosarcina | mazei | SP00120 | Methanosarcina | Thermophila | 19 |
| SP00013 | Methanosarcina | mazei | SP00124 | Methanosarcina | Siciliae | 19 |
| SP00013 | Methanosarcina | mazei | SP00126 | Methanosarcina | Horonobensis | 19 |
| SP00015 | Haloarcula | marismortui | SP00083 | Haloarcula | Hispanica | 19 |
| SP00015 | Haloarcula | marismortui | SP00131 | Haloarcula | sp. CBA1115 | 19 |
| SP00017 | Methanosarcina | barkeri | SP00120 | Methanosarcina | Thermophila | 19 |
| SP00121 | Methanosarcina | sp. WWM596 | SP00017 | Methanosarcina | Barkeri | 19 |
| SP00122 | Methanosarcina | sp. WH1 | SP00017 | Methanosarcina | Barkeri | 19 |
| SP00124 | Methanosarcina | siciliae | SP00017 | Methanosarcina | Barkeri | 19 |
| SP00126 | Methanosarcina | horonobensis | SP00017 | Methanosarcina | Barkeri | 19 |
| SP00017 | Methanosarcina | barkeri | SP00128 | Methanosarcina | Vacuolata | 19 |
| SP00017 | Methanosarcina | barkeri | SP00129 | Methanosarcina | sp. Kolksee | 19 |
| SP00020 | Methanococcoides | burtonii | SP00127 | Methanococcoides | Methylutens | 19 |
| SP00036 | Thermococcus | onnurineus | SP00041 | Thermococcus | Gammatolerans | 19 |
| SP00036 | Thermococcus | onnurineus | SP00081 | Thermococcus | sp. 4557 | 19 |
| SP00036 | Thermococcus | onnurineus | SP00091 | Thermococcus | Cleftensis | 19 |
| SP00036 | Thermococcus | onnurineus | SP00113 | Thermococcus | Paralvinellae | 19 |
| SP00041 | Thermococcus | gammatolerans | SP00081 | Thermococcus | sp. 4557 | 19 |
| SP00041 | Thermococcus | gammatolerans | SP00091 | Thermococcus | Cleftensis | 19 |
| SP00041 | Thermococcus | gammatolerans | SP00113 | Thermococcus | Paralvinellae | 19 |
| SP00042 | Thermococcus | sibiricus | SP00109 | Thermococcus | Litoralis | 19 |
| SP00044 | Halorhabdus | utahensis | SP00108 | Halorhabdus | Tiamatea | 19 |
| SP00045 | Halomicrobium | mukohataei | SP00133 | Halomicrobium | sp. ZPS1 | 19 |
| SP00067 | Thermococcus | barophilus | SP00113 | Thermococcus | Paralvinellae | 19 |
| SP00074 | Pyrococcus | sp. NA2 | SP00080 | Pyrococcus | Yayanosii | 19 |
| SP00074 | Pyrococcus | sp. NA2 | SP00089 | Pyrococcus | sp. ST04 | 19 |
| SP00080 | Pyrococcus | yayanosii | SP00089 | Pyrococcus | sp. ST04 | 19 |
| SP00081 | Thermococcus | sp. 4557 | SP00091 | Thermococcus | Cleftensis | 19 |
| SP00081 | Thermococcus | sp. 4557 | SP00113 | Thermococcus | Paralvinellae | 19 |
| SP00083 | Haloarcula | hispanica | SP00131 | Haloarcula | sp. CBA1115 | 19 |
| SP00091 | Thermococcus | cleftensis | SP00113 | Thermococcus | Paralvinellae | 19 |
| SP00120 | Methanosarcina | thermophila | SP00124 | Methanosarcina | Siciliae | 19 |
| SP00120 | Methanosarcina | thermophila | SP00126 | Methanosarcina | Horonobensis | 19 |
| SP00120 | Methanosarcina | thermophila | SP00128 | Methanosarcina | Vacuolata | 19 |
| SP00120 | Methanosarcina | thermophila | SP00129 | Methanosarcina | sp. Kolksee | 19 |
| SP00121 | Methanosarcina | sp. WWM596 | SP00122 | Methanosarcina | sp. WH1 | 19 |
| SP00121 | Methanosarcina | sp. WWM596 | SP00124 | Methanosarcina | Siciliae | 19 |
| SP00121 | Methanosarcina | sp. WWM596 | SP00125 | Methanosarcina | Lacustris | 19 |
| SP00121 | Methanosarcina | sp. WWM596 | SP00126 | Methanosarcina | Horonobensis | 19 |
| SP00121 | Methanosarcina | sp. WWM596 | SP00128 | Methanosarcina | Vacuolata | 19 |
| SP00121 | Methanosarcina | sp. WWM596 | SP00129 | Methanosarcina | sp. Kolksee | 19 |
| SP00122 | Methanosarcina | sp. WH1 | SP00124 | Methanosarcina | Siciliae | 19 |
| SP00122 | Methanosarcina | sp. WH1 | SP00125 | Methanosarcina | Lacustris | 19 |
| SP00122 | Methanosarcina | sp. WH1 | SP00126 | Methanosarcina | Horonobensis | 19 |
| SP00122 | Methanosarcina | sp. WH1 | SP00128 | Methanosarcina | Vacuolata | 19 |
| SP00122 | Methanosarcina | sp. WH1 | SP00129 | Methanosarcina | sp. Kolksee | 19 |
| SP00124 | Methanosarcina | siciliae | SP00126 | Methanosarcina | Horonobensis | 19 |
| SP00124 | Methanosarcina | siciliae | SP00128 | Methanosarcina | Vacuolata | 19 |
| SP00124 | Methanosarcina | siciliae | SP00129 | Methanosarcina | sp. Kolksee | 19 |
| SP00126 | Methanosarcina | horonobensis | SP00128 | Methanosarcina | Vacuolata | 19 |
| SP00126 | Methanosarcina | horonobensis | SP00129 | Methanosarcina | sp. Kolksee | 19 |
| SP00128 | Methanosarcina | vacuolata | SP00129 | Methanosarcina | sp. Kolksee | 19 |
| SP00006 | Pyrococcus | horikoshii | SP00036 | Thermococcus | Onnurineus | 18 |
| SP00017 | Methanosarcina | barkeri | SP00125 | Methanosarcina | Lacustris | 18 |
| SP00036 | Thermococcus | onnurineus | SP00067 | Thermococcus | Barophilus | 18 |
| SP00036 | Thermococcus | onnurineus | SP00080 | Pyrococcus | Yayanosii | 18 |
| SP00036 | Thermococcus | onnurineus | SP00089 | Pyrococcus | sp. ST04 | 18 |
| SP00036 | Thermococcus | onnurineus | SP00109 | Thermococcus | Litoralis | 18 |
| SP00037 | Desulfurococcus | amylolyticus | SP00068 | Desulfurococcus | Mucosus | 18 |
| SP00041 | Thermococcus | gammatolerans | SP00067 | Thermococcus | Barophilus | 18 |
| SP00042 | Thermococcus | sibiricus | SP00067 | Thermococcus | Barophilus | 18 |
| SP00042 | Thermococcus | sibiricus | SP00113 | Thermococcus | Paralvinellae | 18 |
| SP00067 | Thermococcus | barophilus | SP00081 | Thermococcus | sp. 4557 | 18 |
| SP00067 | Thermococcus | barophilus | SP00091 | Thermococcus | Cleftensis | 18 |
| SP00067 | Thermococcus | barophilus | SP00109 | Thermococcus | Litoralis | 18 |
| SP00081 | Thermococcus | sp. 4557 | SP00109 | Thermococcus | Litoralis | 18 |
| SP00091 | Thermococcus | cleftensis | SP00109 | Thermococcus | Litoralis | 18 |
| SP00109 | Thermococcus | litoralis | SP00113 | Thermococcus | Paralvinellae | 18 |
| SP00116 | Nitrososphaera | evergladensis (c.) | SP00117 | Nitrososphaera | Viennensis | 18 |
| SP00012 | Methanosarcina | acetivorans | SP00125 | Methanosarcina | Lacustris | 17 |
| SP00017 | Methanosarcina | barkeri | SP00013 | Methanosarcina | Mazei | 17 |
| SP00121 | Methanosarcina | sp. WWM596 | SP00013 | Methanosarcina | Mazei | 17 |
| SP00122 | Methanosarcina | sp. WH1 | SP00013 | Methanosarcina | Mazei | 17 |
| SP00013 | Methanosarcina | mazei | SP00128 | Methanosarcina | Vacuolata | 17 |
| SP00013 | Methanosarcina | mazei | SP00129 | Methanosarcina | sp. Kolksee | 17 |
| SP00018 | Natronomonas | pharaonis | SP00102 | Natronomonas | Moolapensis | 17 |
| SP00036 | Thermococcus | onnurineus | SP00042 | Thermococcus | Sibiricus | 17 |
| SP00036 | Thermococcus | onnurineus | SP00074 | Pyrococcus | sp. NA2 | 17 |
| SP00041 | Thermococcus | gammatolerans | SP00042 | Thermococcus | Sibiricus | 17 |
| SP00041 | Thermococcus | gammatolerans | SP00109 | Thermococcus | Litoralis | 17 |
| SP00042 | Thermococcus | sibiricus | SP00081 | Thermococcus | sp. 4557 | 17 |
| SP00042 | Thermococcus | sibiricus | SP00091 | Thermococcus | Cleftensis | 17 |
| SP00042 | Thermococcus | sibiricus | SP00111 | Palaeococcus | Pacificus | 17 |
| SP00049 | Haloterrigena | turkmenica | SP00092 | Natrinema | sp. J7-2 | 17 |
| SP00092 | Natrinema | sp. J7-2 | SP00098 | Natrinema | Pellirubrum | 17 |
| SP00124 | Methanosarcina | siciliae | SP00125 | Methanosarcina | Lacustris | 17 |
| SP00125 | Methanosarcina | lacustris | SP00126 | Methanosarcina | Horonobensis | 17 |
| SP00005 | Methanocaldococcus | jannaschii | SP00046 | Methanocaldococcus | Vulcanius | 16 |
| SP00046 | Methanocaldococcus | vulcanius | SP00052 | Methanocaldococcus | sp. FS406-22 | 16 |
| SP00048 | Archaeoglobus | profundus | SP00051 | Ferroglobus | Placidus | 16 |
| SP00070 | Thermoproteus | uzoniensis | SP00084 | Thermoproteus | Tenax | 16 |
| SP00094 | Nitrososphaera | gargensis (c.) | SP00116 | Nitrososphaera | evergladensis (c.) | 16 |
| SP00094 | Nitrososphaera | gargensis (c.) | SP00117 | Nitrososphaera | Viennensis | 16 |
| SP00120 | Methanosarcina | thermophila | SP00125 | Methanosarcina | Lacustris | 16 |
| SP00125 | Methanosarcina | lacustris | SP00128 | Methanosarcina | Vacuolata | 16 |
| SP00125 | Methanosarcina | lacustris | SP00129 | Methanosarcina | sp. Kolksee | 16 |
| SP00013 | Methanosarcina | mazei | SP00125 | Methanosarcina | Lacustris | 15 |
| SP00036 | Thermococcus | onnurineus | SP00111 | Palaeococcus | Pacificus | 15 |
| SP00041 | Thermococcus | gammatolerans | SP00080 | Pyrococcus | Yayanosii | 15 |
| SP00041 | Thermococcus | gammatolerans | SP00089 | Pyrococcus | sp. ST04 | 15 |
| SP00041 | Thermococcus | gammatolerans | SP00111 | Palaeococcus | Pacificus | 15 |
| SP00049 | Haloterrigena | turkmenica | SP00098 | Natrinema | Pellirubrum | 15 |
| SP00069 | Methanobacterium | lacus | SP00076 | Methanobacterium | Paludis | 15 |
| SP00080 | Pyrococcus | yayanosii | SP00091 | Thermococcus | Cleftensis | 15 |
| SP00080 | Pyrococcus | yayanosii | SP00113 | Thermococcus | Paralvinellae | 15 |
| SP00089 | Pyrococcus | sp. ST04 | SP00113 | Thermococcus | Paralvinellae | 15 |
| SP00091 | Thermococcus | cleftensis | SP00111 | Palaeococcus | Pacificus | 15 |
| SP00120 | Methanosarcina | thermophila | SP00121 | Methanosarcina | sp. WWM596 | 15 |
| SP00120 | Methanosarcina | thermophila | SP00122 | Methanosarcina | sp. WH1 | 15 |
| SP00006 | Pyrococcus | horikoshii | SP00113 | Thermococcus | Paralvinellae | 14 |
| SP00027 | Methanoculleus | marisnigri | SP00093 | Methanoculleus | Bourgensis | 14 |
| SP00043 | Methanocaldococcus | fervens | SP00046 | Methanocaldococcus | Vulcanius | 14 |
| SP00046 | Methanocaldococcus | vulcanius | SP00119 | Methanocaldococcus | Bathoardescens | 14 |
| SP00056 | Methanocaldococcus | infernus | SP00119 | Methanocaldococcus | Bathoardescens | 14 |
| SP00109 | Thermococcus | litoralis | SP00111 | Palaeococcus | Pacificus | 14 |
| SP00006 | Pyrococcus | horikoshii | SP00041 | Thermococcus | Gammatolerans | 13 |
| SP00006 | Pyrococcus | horikoshii | SP00067 | Thermococcus | Barophilus | 13 |
| SP00006 | Pyrococcus | horikoshii | SP00091 | Thermococcus | Cleftensis | 13 |
| SP00036 | Thermococcus | onnurineus | SP00118 | Thermococcus | Eurythermalis | 13 |
| SP00041 | Thermococcus | gammatolerans | SP00074 | Pyrococcus | sp. NA2 | 13 |
| SP00041 | Thermococcus | gammatolerans | SP00118 | Thermococcus | Eurythermalis | 13 |
| SP00052 | Methanocaldococcus | sp. FS406-22 | SP00056 | Methanocaldococcus | Infernus | 13 |
| SP00067 | Thermococcus | barophilus | SP00074 | Pyrococcus | sp. NA2 | 13 |
| SP00067 | Thermococcus | barophilus | SP00080 | Pyrococcus | Yayanosii | 13 |
| SP00067 | Thermococcus | barophilus | SP00089 | Pyrococcus | sp. ST04 | 13 |
| SP00074 | Pyrococcus | sp. NA2 | SP00091 | Thermococcus | Cleftensis | 13 |
| SP00074 | Pyrococcus | sp. NA2 | SP00113 | Thermococcus | Paralvinellae | 13 |
| SP00076 | Methanobacterium | paludis | SP00135 | Methanobacterium | Congolense | 13 |
| SP00080 | Pyrococcus | yayanosii | SP00118 | Thermococcus | Eurythermalis | 13 |
| SP00089 | Pyrococcus | sp. ST04 | SP00091 | Thermococcus | Cleftensis | 13 |
| SP00091 | Thermococcus | cleftensis | SP00118 | Thermococcus | Eurythermalis | 13 |
| SP00113 | Thermococcus | paralvinellae | SP00118 | Thermococcus | Eurythermalis | 13 |
| SP00006 | Pyrococcus | horikoshii | SP00118 | Thermococcus | Eurythermalis | 12 |
| SP00069 | Methanobacterium | lacus | SP00135 | Methanobacterium | Congolense | 12 |
| SP00074 | Pyrococcus | sp. NA2 | SP00118 | Thermococcus | Eurythermalis | 12 |
| SP00089 | Pyrococcus | sp. ST04 | SP00118 | Thermococcus | Eurythermalis | 12 |
| SP00067 | Thermococcus | barophilus | SP00118 | Thermococcus | Eurythermalis | 11 |
| SP00109 | Thermococcus | litoralis | SP00118 | Thermococcus | Eurythermalis | 11 |
| SP00006 | Pyrococcus | horikoshii | SP00109 | Thermococcus | Litoralis | 10 |
| SP00024 | Hyperthermus | butylicus | SP00082 | Pyrolobus | Fumarii | 10 |
| SP00067 | Thermococcus | barophilus | SP00111 | Palaeococcus | Pacificus | 10 |
| SP00074 | Pyrococcus | sp. NA2 | SP00109 | Thermococcus | Litoralis | 10 |
| SP00080 | Pyrococcus | yayanosii | SP00109 | Thermococcus | Litoralis | 10 |
| SP00089 | Pyrococcus | sp. ST04 | SP00109 | Thermococcus | Litoralis | 10 |
| SP00111 | Palaeococcus | pacificus | SP00113 | Thermococcus | Paralvinellae | 10 |
| SP00004 | Aeropyrum | pernix | SP00110 | Aeropyrum | Camini | 9 |
| SP00033 | Methanoregula | boonei | SP00097 | Methanoregula | Formicica | 9 |
| SP00049 | Haloterrigena | turkmenica | SP00053 | Natrialba | Magadii | 9 |
| SP00081 | Thermococcus | sp. 4557 | SP00118 | Thermococcus | Eurythermalis | 9 |
| SP00042 | Thermococcus | sibiricus | SP00080 | Pyrococcus | Yayanosii | 8 |
| SP00043 | Methanocaldococcus | fervens | SP00056 | Methanocaldococcus | Infernus | 8 |
| SP00006 | Pyrococcus | horikoshii | SP00042 | Thermococcus | Sibiricus | 7 |
| SP00008 | Halobacterium | salinarum | SP00115 | Halobacterium | sp. DL1 | 7 |
| SP00010 | Pyrobaculum | aerophilum | SP00028 | Pyrobaculum | Arsenaticum | 7 |
| SP00010 | Pyrobaculum | aerophilum | SP00084 | Thermoproteus | Tenax | 7 |
| SP00042 | Thermococcus | sibiricus | SP00118 | Thermococcus | Eurythermalis | 7 |
| SP00080 | Pyrococcus | yayanosii | SP00081 | Thermococcus | sp. 4557 | 7 |
| SP00111 | Palaeococcus | pacificus | SP00118 | Thermococcus | Eurythermalis | 7 |
| SP00006 | Pyrococcus | horikoshii | SP00081 | Thermococcus | sp. 4557 | 6 |
| SP00006 | Pyrococcus | horikoshii | SP00111 | Palaeococcus | Pacificus | 6 |
| SP00007 | Methanococcus | maripaludis | SP00031 | Methanococcus | Vannielii | 6 |
| SP00010 | Pyrobaculum | aerophilum | SP00070 | Thermoproteus | Uzoniensis | 6 |
| SP00028 | Pyrobaculum | arsenaticum | SP00070 | Thermoproteus | Uzoniensis | 6 |
| SP00028 | Pyrobaculum | arsenaticum | SP00084 | Thermoproteus | Tenax | 6 |
| SP00042 | Thermococcus | sibiricus | SP00089 | Pyrococcus | sp. ST04 | 6 |
| SP00071 | Archaeoglobus | veneficus | SP00104 | Archaeoglobus | Sulfaticallidus | 6 |
| SP00074 | Pyrococcus | sp. NA2 | SP00111 | Palaeococcus | Pacificus | 6 |
| SP00080 | Pyrococcus | yayanosii | SP00111 | Palaeococcus | Pacificus | 6 |
| SP00081 | Thermococcus | sp. 4557 | SP00111 | Palaeococcus | Pacificus | 6 |
| SP00089 | Pyrococcus | sp. ST04 | SP00111 | Palaeococcus | Pacificus | 6 |
| SP00003 | Pyrobaculum | oguniense | SP00010 | Pyrobaculum | Aerophilum | 5 |
| SP00003 | Pyrobaculum | oguniense | SP00028 | Pyrobaculum | Arsenaticum | 5 |
| SP00003 | Pyrobaculum | oguniense | SP00070 | Thermoproteus | Uzoniensis | 5 |
| SP00003 | Pyrobaculum | oguniense | SP00084 | Thermoproteus | Tenax | 5 |
| SP00005 | Methanocaldococcus | jannaschii | SP00056 | Methanocaldococcus | Infernus | 5 |
| SP00026 | Staphylothermus | marinus | SP00057 | Staphylothermus | Hellenicus | 5 |
| SP00042 | Thermococcus | sibiricus | SP00074 | Pyrococcus | sp. NA2 | 5 |
| SP00074 | Pyrococcus | sp. NA2 | SP00081 | Thermococcus | sp. 4557 | 5 |
| SP00076 | Methanobacterium | paludis | SP00112 | Methanobacterium | Formicicum | 5 |
| SP00081 | Thermococcus | sp. 4557 | SP00089 | Pyrococcus | sp. ST04 | 5 |
| SP00123 | Methanosarcina | sp. MTP4 | SP00013 | Methanosarcina | Mazei | 4 |
| SP00026 | Staphylothermus | marinus | SP00090 | Thermogladius | Calderae | 4 |
| SP00045 | Halomicrobium | mukohataei | SP00083 | Haloarcula | Hispanica | 4 |
| SP00045 | Halomicrobium | mukohataei | SP00131 | Haloarcula | sp. CBA1115 | 4 |
| SP00057 | Staphylothermus | hellenicus | SP00068 | Desulfurococcus | Mucosus | 4 |
| SP00057 | Staphylothermus | hellenicus | SP00082 | Pyrolobus | Fumarii | 4 |
| SP00057 | Staphylothermus | hellenicus | SP00090 | Thermogladius | Calderae | 4 |
| SP00078 | Halopiger | xanaduensis | SP00092 | Natrinema | sp. J7-2 | 4 |
| SP00078 | Halopiger | xanaduensis | SP00098 | Natrinema | Pellirubrum | 4 |
| SP00083 | Haloarcula | hispanica | SP00133 | Halomicrobium | sp. ZPS1 | 4 |
| SP00112 | Methanobacterium | formicicum | SP00135 | Methanobacterium | Congolense | 4 |
| SP00123 | Methanosarcina | sp. MTP4 | SP00128 | Methanosarcina | Vacuolata | 4 |
| SP00123 | Methanosarcina | sp. MTP4 | SP00129 | Methanosarcina | sp. Kolksee | 4 |
| SP00131 | Haloarcula | sp. CBA1115 | SP00133 | Halomicrobium | sp. ZPS1 | 4 |
| SP00012 | Methanosarcina | acetivorans | SP00123 | Methanosarcina | sp. MTP4 | 3 |
| SP00015 | Haloarcula | marismortui | SP00045 | Halomicrobium | Mukohataei | 3 |
| SP00015 | Haloarcula | marismortui | SP00133 | Halomicrobium | sp. ZPS1 | 3 |
| SP00044 | Halorhabdus | utahensis | SP00045 | Halomicrobium | Mukohataei | 3 |
| SP00044 | Halorhabdus | utahensis | SP00133 | Halomicrobium | sp. ZPS1 | 3 |
| SP00045 | Halomicrobium | mukohataei | SP00108 | Halorhabdus | Tiamatea | 3 |
| SP00051 | Ferroglobus | placidus | SP00071 | Archaeoglobus | Veneficus | 3 |
| SP00053 | Natrialba | magadii | SP00092 | Natrinema | sp. J7-2 | 3 |
| SP00053 | Natrialba | magadii | SP00098 | Natrinema | Pellirubrum | 3 |
| SP00053 | Natrialba | magadii | SP00114 | Halostagnicola | Larsenii | 3 |
| SP00055 | Methanohalophilus | mahii | SP00127 | Methanococcoides | Methylutens | 3 |
| SP00069 | Methanobacterium | lacus | SP00112 | Methanobacterium | Formicicum | 3 |
| SP00108 | Halorhabdus | tiamatea | SP00133 | Halomicrobium | sp. ZPS1 | 3 |
| SP00001 | Ignisphaera | aggregans | SP00024 | Hyperthermus | Butylicus | 2 |
| SP00001 | Ignisphaera | aggregans | SP00026 | Staphylothermus | Marinus | 2 |
| SP00001 | Ignisphaera | aggregans | SP00057 | Staphylothermus | Hellenicus | 2 |
| SP00001 | Ignisphaera | aggregans | SP00082 | Pyrolobus | Fumarii | 2 |
| SP00003 | Pyrobaculum | oguniense | SP00086 | Pyrobaculum | Ferrireducens | 2 |
| SP00005 | Methanocaldococcus | jannaschii | SP00075 | Methanotorris | Igneus | 2 |
| SP00007 | Methanococcus | maripaludis | SP00058 | Methanococcus | Voltae | 2 |
| SP00010 | Pyrobaculum | aerophilum | SP00086 | Pyrobaculum | Ferrireducens | 2 |
| SP00015 | Haloarcula | marismortui | SP00044 | Halorhabdus | Utahensis | 2 |
| SP00015 | Haloarcula | marismortui | SP00108 | Halorhabdus | Tiamatea | 2 |
| SP00017 | Methanosarcina | barkeri | SP00123 | Methanosarcina | sp. MTP4 | 2 |
| SP00018 | Natronomonas | pharaonis | SP00045 | Halomicrobium | Mukohataei | 2 |
| SP00018 | Natronomonas | pharaonis | SP00133 | Halomicrobium | sp. ZPS1 | 2 |
| SP00020 | Methanococcoides | burtonii | SP00055 | Methanohalophilus | Mahii | 2 |
| SP00020 | Methanococcoides | burtonii | SP00079 | Methanosalsum | Zhilinae | 2 |
| SP00024 | Hyperthermus | butylicus | SP00026 | Staphylothermus | Marinus | 2 |
| SP00024 | Hyperthermus | butylicus | SP00034 | Ignicoccus | Hospitalis | 2 |
| SP00024 | Hyperthermus | butylicus | SP00057 | Staphylothermus | Hellenicus | 2 |
| SP00024 | Hyperthermus | butylicus | SP00090 | Thermogladius | Calderae | 2 |
| SP00026 | Staphylothermus | marinus | SP00082 | Pyrolobus | Fumarii | 2 |
| SP00028 | Pyrobaculum | arsenaticum | SP00086 | Pyrobaculum | Ferrireducens | 2 |
| SP00034 | Ignicoccus | hospitalis | SP00082 | Pyrolobus | Fumarii | 2 |
| SP00043 | Methanocaldococcus | fervens | SP00075 | Methanotorris | Igneus | 2 |
| SP00044 | Halorhabdus | utahensis | SP00083 | Haloarcula | Hispanica | 2 |
| SP00044 | Halorhabdus | utahensis | SP00131 | Haloarcula | sp. CBA1115 | 2 |
| SP00045 | Halomicrobium | mukohataei | SP00102 | Natronomonas | Moolapensis | 2 |
| SP00049 | Haloterrigena | turkmenica | SP00078 | Halopiger | Xanaduensis | 2 |
| SP00049 | Haloterrigena | turkmenica | SP00114 | Halostagnicola | Larsenii | 2 |
| SP00050 | Methanobrevibacter | ruminantium | SP00106 | Methanobrevibacter | sp. AbM4 | 2 |
| SP00052 | Methanocaldococcus | sp. FS406-22 | SP00075 | Methanotorris | Igneus | 2 |
| SP00055 | Methanohalophilus | mahii | SP00079 | Methanosalsum | Zhilinae | 2 |
| SP00062 | Methanothermobacter | marburgensis | SP00112 | Methanobacterium | Formicicum | 2 |
| SP00062 | Methanothermobacter | marburgensis | SP00135 | Methanobacterium | Congolense | 2 |
| SP00068 | Desulfurococcus | mucosus | SP00090 | Thermogladius | Calderae | 2 |
| SP00075 | Methanotorris | igneus | SP00119 | Methanocaldococcus | Bathoardescens | 2 |
| SP00079 | Methanosalsum | zhilinae | SP00127 | Methanococcoides | Methylutens | 2 |
| SP00082 | Pyrolobus | fumarii | SP00090 | Thermogladius | Calderae | 2 |
| SP00083 | Haloarcula | hispanica | SP00108 | Halorhabdus | Tiamatea | 2 |
| SP00084 | Thermoproteus | tenax | SP00086 | Pyrobaculum | Ferrireducens | 2 |
| SP00092 | Natrinema | sp. J7-2 | SP00114 | Halostagnicola | Larsenii | 2 |
| SP00096 | Natronobacterium | gregoryi | SP00114 | Halostagnicola | Larsenii | 2 |
| SP00102 | Natronomonas | moolapensis | SP00133 | Halomicrobium | sp. ZPS1 | 2 |
| SP00108 | Halorhabdus | tiamatea | SP00131 | Haloarcula | sp. CBA1115 | 2 |
| SP00121 | Methanosarcina | sp. WWM596 | SP00123 | Methanosarcina | sp. MTP4 | 2 |
| SP00122 | Methanosarcina | sp. WH1 | SP00123 | Methanosarcina | sp. MTP4 | 2 |
| SP00123 | Methanosarcina | sp. MTP4 | SP00124 | Methanosarcina | Siciliae | 2 |
| SP00123 | Methanosarcina | sp. MTP4 | SP00125 | Methanosarcina | Lacustris | 2 |
| SP00123 | Methanosarcina | sp. MTP4 | SP00126 | Methanosarcina | Horonobensis | 2 |
| SP00001 | Ignisphaera | aggregans | SP00034 | Ignicoccus | Hospitalis | 1 |
| SP00002 | Methanolobus | psychrophilus | SP00055 | Methanohalophilus | Mahii | 1 |
| SP00004 | Aeropyrum | pernix | SP00024 | Hyperthermus | Butylicus | 1 |
| SP00004 | Aeropyrum | pernix | SP00026 | Staphylothermus | Marinus | 1 |
| SP00004 | Aeropyrum | pernix | SP00034 | Ignicoccus | Hospitalis | 1 |
| SP00004 | Aeropyrum | pernix | SP00057 | Staphylothermus | Hellenicus | 1 |
| SP00004 | Aeropyrum | pernix | SP00082 | Pyrolobus | Fumarii | 1 |
| SP00010 | Pyrobaculum | aerophilum | SP00064 | Vulcanisaeta | Distributa | 1 |
| SP00018 | Natronomonas | pharaonis | SP00044 | Halorhabdus | Utahensis | 1 |
| SP00018 | Natronomonas | pharaonis | SP00108 | Halorhabdus | Tiamatea | 1 |
| SP00018 | Natronomonas | pharaonis | SP00115 | Halobacterium | sp. DL1 | 1 |
| SP00020 | Methanococcoides | burtonii | SP00100 | Methanomethylovorans | Hollandica | 1 |
| SP00022 | Methanothrix | thermoacetophila | SP00072 | Methanothrix | Soehngenii | 1 |
| SP00024 | Hyperthermus | butylicus | SP00068 | Desulfurococcus | Mucosus | 1 |
| SP00024 | Hyperthermus | butylicus | SP00110 | Aeropyrum | Camini | 1 |
| SP00026 | Staphylothermus | marinus | SP00034 | Ignicoccus | Hospitalis | 1 |
| SP00026 | Staphylothermus | marinus | SP00037 | Desulfurococcus | Amylolyticus | 1 |
| SP00026 | Staphylothermus | marinus | SP00061 | Acidilobus | Saccharovorans | 1 |
| SP00026 | Staphylothermus | marinus | SP00068 | Desulfurococcus | Mucosus | 1 |
| SP00028 | Pyrobaculum | arsenaticum | SP00064 | Vulcanisaeta | Distributa | 1 |
| SP00030 | Methanobrevibacter | smithii | SP00050 | Methanobrevibacter | Ruminantium | 1 |
| SP00031 | Methanococcus | vannielii | SP00077 | Methanothermococcus | Okinawensis | 1 |
| SP00032 | Methanococcus | aeolicus | SP00077 | Methanothermococcus | Okinawensis | 1 |
| SP00034 | Ignicoccus | hospitalis | SP00057 | Staphylothermus | Hellenicus | 1 |
| SP00034 | Ignicoccus | hospitalis | SP00068 | Desulfurococcus | Mucosus | 1 |
| SP00034 | Ignicoccus | hospitalis | SP00090 | Thermogladius | Calderae | 1 |
| SP00034 | Ignicoccus | hospitalis | SP00110 | Aeropyrum | Camini | 1 |
| SP00037 | Desulfurococcus | amylolyticus | SP00057 | Staphylothermus | Hellenicus | 1 |
| SP00037 | Desulfurococcus | amylolyticus | SP00090 | Thermogladius | Calderae | 1 |
| SP00038 | Methanosphaerula | palustris | SP00093 | Methanoculleus | Bourgensis | 1 |
| SP00044 | Halorhabdus | utahensis | SP00102 | Natronomonas | Moolapensis | 1 |
| SP00045 | Halomicrobium | mukohataei | SP00115 | Halobacterium | sp. DL1 | 1 |
| SP00046 | Methanocaldococcus | vulcanius | SP00056 | Methanocaldococcus | Infernus | 1 |
| SP00049 | Haloterrigena | turkmenica | SP00096 | Natronobacterium | Gregoryi | 1 |
| SP00049 | Haloterrigena | turkmenica | SP00099 | Halovivax | Ruber | 1 |
| SP00049 | Haloterrigena | turkmenica | SP00101 | Natronococcus | Occultus | 1 |
| SP00051 | Ferroglobus | placidus | SP00118 | Thermococcus | Eurythermalis | 1 |
| SP00053 | Natrialba | magadii | SP00078 | Halopiger | Xanaduensis | 1 |
| SP00053 | Natrialba | magadii | SP00096 | Natronobacterium | Gregoryi | 1 |
| SP00053 | Natrialba | magadii | SP00099 | Halovivax | Ruber | 1 |
| SP00053 | Natrialba | magadii | SP00101 | Natronococcus | Occultus | 1 |
| SP00057 | Staphylothermus | hellenicus | SP00061 | Acidilobus | Saccharovorans | 1 |
| SP00059 | Methanohalobium | evestigatum | SP00079 | Methanosalsum | Zhilinae | 1 |
| SP00062 | Methanothermobacter | marburgensis | SP00069 | Methanobacterium | Lacus | 1 |
| SP00064 | Vulcanisaeta | distributa | SP00070 | Thermoproteus | Uzoniensis | 1 |
| SP00064 | Vulcanisaeta | distributa | SP00084 | Thermoproteus | Tenax | 1 |
| SP00068 | Desulfurococcus | mucosus | SP00082 | Pyrolobus | Fumarii | 1 |
| SP00069 | Methanobacterium | lacus | SP00134 | Methanosphaera | Stadtmanae | 1 |
| SP00070 | Thermoproteus | uzoniensis | SP00086 | Pyrobaculum | Ferrireducens | 1 |
| SP00078 | Halopiger | xanaduensis | SP00096 | Natronobacterium | Gregoryi | 1 |
| SP00078 | Halopiger | xanaduensis | SP00099 | Halovivax | Ruber | 1 |
| SP00078 | Halopiger | xanaduensis | SP00101 | Natronococcus | Occultus | 1 |
| SP00078 | Halopiger | xanaduensis | SP00114 | Halostagnicola | Larsenii | 1 |
| SP00092 | Natrinema | sp. J7-2 | SP00096 | Natronobacterium | Gregoryi | 1 |
| SP00092 | Natrinema | sp. J7-2 | SP00099 | Halovivax | Ruber | 1 |
| SP00092 | Natrinema | sp. J7-2 | SP00101 | Natronococcus | Occultus | 1 |
| SP00096 | Natronobacterium | gregoryi | SP00098 | Natrinema | Pellirubrum | 1 |
| SP00096 | Natronobacterium | gregoryi | SP00101 | Natronococcus | Occultus | 1 |
| SP00098 | Natrinema | pellirubrum | SP00099 | Halovivax | Ruber | 1 |
| SP00098 | Natrinema | pellirubrum | SP00101 | Natronococcus | Occultus | 1 |
| SP00098 | Natrinema | pellirubrum | SP00114 | Halostagnicola | Larsenii | 1 |
| SP00099 | Halovivax | ruber | SP00114 | Halostagnicola | Larsenii | 1 |
| SP00101 | Natronococcus | occultus | SP00114 | Halostagnicola | Larsenii | 1 |
| SP00102 | Natronomonas | moolapensis | SP00108 | Halorhabdus | Tiamatea | 1 |
| SP00102 | Natronomonas | moolapensis | SP00115 | Halobacterium | sp. DL1 | 1 |
| SP00115 | Halobacterium | sp. DL1 | SP00133 | Halomicrobium | sp. ZPS1 | 1 |
| ***TOTAL*** | | | | | | 3232 |

The Table details all the pairs of archaeal species which had amplicon similarity/identity ≥97% using the archaeal-specific and the bacterial and archaeal primer pairs analyzed in the present study. C.= candidatus; Freq.= frequency, number times that a pair of species had amplicon similarity/identity ≥97% in the different primer pairs; ID= species identifier.

Appendix Table 20. Genus of the pairs of archaeal species with amplicon similarity ≥97%.

| **Pair of genus** | **Frequency** |
| --- | --- |
| Methanosarcina\|Methanosarcina | 1034 |
| Thermococcus\|Thermococcus | 601 |
| Methanocaldococcus\|Methanocaldococcus | 221 |
| Pyrococcus\|Pyrococcus | 114 |
| Haloarcula\|Haloarcula | 57 |
| Methanobacterium\|Methanobacterium | 52 |
| Nitrososphaera\|Nitrososphaera | 50 |
| Methanobrevibacter\|Methanobrevibacter | 23 |
| Pyrobaculum\|Pyrobaculum | 23 |
| Halomicrobium\|Halomicrobium | 19 |
| Halorhabdus\|Halorhabdus | 19 |
| Methanococcoides\|Methanococcoides | 19 |
| Desulfurococcus\|Desulfurococcus | 18 |
| Natrinema\|Natrinema | 17 |
| Natronomonas\|Natronomonas | 17 |
| Thermoproteus\|Thermoproteus | 16 |
| Methanoculleus\|Methanoculleus | 14 |
| Aeropyrum\|Aeropyrum | 9 |
| Methanoregula\|Methanoregula | 9 |
| Methanococcus\|Methanococcus | 8 |
| Halobacterium\|Halobacterium | 7 |
| Archaeoglobus\|Archaeoglobus | 6 |
| Staphylothermus\|Staphylothermus | 5 |
| Methanothrix\|Methanothrix | 1 |
| ***TOTAL*** | 2359 |
| **Pair of different genus** | **Frequency** |
| Pyrococcus\|Thermococcus | 428 |
| Palaeococcus\|Thermococcus | 109 |
| Pyrobaculum\|Thermoproteus | 38 |
| Haloterrigena\|Natrinema | 32 |
| Palaeococcus\|Pyrococcus | 24 |
| Haloarcula\|Halomicrobium | 22 |
| Archaeoglobus\|Ferroglobus | 19 |
| Haloarcula\|Halorhabdus | 12 |
| Halomicrobium\|Halorhabdus | 12 |
| Hyperthermus\|Pyrolobus | 10 |
| Haloterrigena\|Natrialba | 9 |
| Halomicrobium\|Natronomonas | 8 |
| Halopiger\|Natrinema | 8 |
| Methanocaldococcus\|Methanotorris | 8 |
| Staphylothermus\|Thermogladius | 8 |
| Desulfurococcus\|Staphylothermus | 7 |
| Natrialba\|Natrinema | 6 |
| Pyrolobus\|Staphylothermus | 6 |
| Methanobacterium\|Methanothermobacter | 5 |
| Methanococcoides\|Methanohalophilus | 5 |
| Halorhabdus\|Natronomonas | 4 |
| Hyperthermus\|Staphylothermus | 4 |
| Ignisphaera\|Staphylothermus | 4 |
| Methanococcoides\|Methanosalsum | 4 |
| Desulfurococcus\|Thermogladius | 3 |
| Halostagnicola\|Natrialba | 3 |
| Halostagnicola\|Natrinema | 3 |
| Acidilobus\|Staphylothermus | 2 |
| Aeropyrum\|Hyperthermus | 2 |
| Aeropyrum\|Ignicoccus | 2 |
| Aeropyrum\|Staphylothermus | 2 |
| Halobacterium\|Halomicrobium | 2 |
| Halobacterium\|Natronomonas | 2 |
| Halopiger\|Haloterrigena | 2 |
| Halostagnicola\|Haloterrigena | 2 |
| Halostagnicola\|Natronobacterium | 2 |
| Halovivax\|Natrinema | 2 |
| Hyperthermus\|Ignicoccus | 2 |
| Hyperthermus\|Ignisphaera | 2 |
| Hyperthermus\|Thermogladius | 2 |
| Ignicoccus\|Pyrolobus | 2 |
| Ignicoccus\|Staphylothermus | 2 |
| Ignisphaera\|Pyrolobus | 2 |
| Methanococcus\|Methanothermococcus | 2 |
| Methanohalophilus\|Methanosalsum | 2 |
| Natrinema\|Natronobacterium | 2 |
| Natrinema\|Natronococcus | 2 |
| Pyrobaculum\|Vulcanisaeta | 2 |
| Pyrolobus\|Thermogladius | 2 |
| Thermoproteus\|Vulcanisaeta | 2 |
| Aeropyrum\|Pyrolobus | 1 |
| Desulfurococcus\|Hyperthermus | 1 |
| Desulfurococcus\|Ignicoccus | 1 |
| Desulfurococcus\|Pyrolobus | 1 |
| Ferroglobus\|Thermococcus | 1 |
| Halopiger\|Halostagnicola | 1 |
| Halopiger\|Halovivax | 1 |
| Halopiger\|Natrialba | 1 |
| Halopiger\|Natronobacterium | 1 |
| Halopiger\|Natronococcus | 1 |
| Halostagnicola\|Halovivax | 1 |
| Halostagnicola\|Natronococcus | 1 |
| Haloterrigena\|Halovivax | 1 |
| Haloterrigena\|Natronobacterium | 1 |
| Haloterrigena\|Natronococcus | 1 |
| Halovivax\|Natrialba | 1 |
| Ignicoccus\|Ignisphaera | 1 |
| Ignicoccus\|Thermogladius | 1 |
| Methanobacterium\|Methanosphaera | 1 |
| Methanococcoides\|Methanomethylovorans | 1 |
| Methanoculleus\|Methanosphaerula | 1 |
| Methanohalobium\|Methanosalsum | 1 |
| Methanohalophilus\|Methanolobus | 1 |
| Natrialba\|Natronobacterium | 1 |
| Natrialba\|Natronococcus | 1 |
| Natronobacterium\|Natronococcus | 1 |
| ***TOTAL*** | 873 |

Appendix Table 21. Families of the pairs of archaeal species with amplicon similarity ≥97%.

| **Pair of families** | **Frequency** |
| --- | --- |
| Thermococcaceae\|Thermococcaceae | 1276 |
| Methanosarcinaceae\|Methanosarcinaceae | 1067 |
| Methanocaldococcaceae\|Methanocaldococcaceae | 229 |
| Haloarculaceae\|Haloarculaceae | 170 |
| Natrialbaceae\|Natrialbaceae | 104 |
| Methanobacteriaceae\|Methanobacteriaceae | 81 |
| Thermoproteaceae\|Thermoproteaceae | 81 |
| Desulfurococcaceae\|Desulfurococcaceae | 63 |
| Nitrososphaeraceae\|Nitrososphaeraceae | 50 |
| Archaeoglobaceae\|Archaeoglobaceae | 25 |
| Methanomicrobiaceae\|Methanomicrobiaceae | 14 |
| Methanococcaceae\|Methanococcaceae | 10 |
| Pyrodictiaceae\|Pyrodictiaceae | 10 |
| Methanoregulaceae\|Methanoregulaceae | 9 |
| Halobacteriaceae\|Halobacteriaceae | 7 |
| Methanotrichaceae\|Methanotrichaceae | 1 |
| ***TOTAL*** | 3197 |
| **Pair of different families** | **Frequency** |
| Desulfurococcaceae\|Pyrodictiaceae | 27 |
| Haloarculaceae\|Halobacteriaceae | 4 |
| Acidilobaceae\|Desulfurococcaceae | 2 |
| Archaeoglobaceae\|Thermococcaceae | 1 |
| Methanomicrobiaceae\|Methanoregulaceae | 1 |
| ***TOTAL*** | 35 |

Appendix Table 22. Orders of the pairs of archaeal species with amplicon similarity ≥97%.

| **Pair of orders** | **Frequency** |
| --- | --- |
| Thermococcales\|Thermococcales | 1276 |
| Methanosarcinales\|Methanosarcinales | 1068 |
| Methanococcales\|Methanococcales | 239 |
| Halobacteriales\|Halobacteriales | 181 |
| Natrialbales\|Natrialbales | 104 |
| Desulfurococcales\|Desulfurococcales | 100 |
| Methanobacteriales\|Methanobacteriales | 81 |
| Thermoproteales\|Thermoproteales | 81 |
| Nitrososphaerales\|Nitrososphaerales | 50 |
| Archaeoglobales\|Archaeoglobales | 25 |
| Methanomicrobiales\|Methanomicrobiales | 24 |
| ***TOTAL*** | 3229 |
| **Pair of different orders** | **Frequency** |
| Acidilobales\|Desulfurococcales | 2 |
| Archaeoglobales\|Thermococcales | 1 |
| ***TOTAL*** | 3 |

Appendix Table 23. Classes of the pairs of archaeal species with amplicon similarity ≥97%.

| **Pair of classes** | **Frequency** |
| --- | --- |
| Thermococci\|Thermococci | 1276 |
| Methanomicrobia\|Methanomicrobia | 1092 |
| Halobacteria\|Halobacteria | 285 |
| Methanococci\|Methanococci | 239 |
| Thermoprotei\|Thermoprotei | 183 |
| Methanobacteria\|Methanobacteria | 81 |
| Nitrososphaeria\|Nitrososphaeria | 50 |
| Archaeoglobi\|Archaeoglobi | 25 |
| ***TOTAL*** | 3231 |
| **Pair of different classes** | **Frequency** |
| Archaeoglobi\|Thermococci | 1 |
| ***TOTAL*** | 1 |
